# An Automated HDX-MS Platform for *in situ* characterisation of Membrane Proteins

**DOI:** 10.64898/2026.03.06.710074

**Authors:** Charlotte Guffick, Juan Pablo Rincon Pabon, Damon Griffiths, Satomi Inaba-Inoue, Konstantinos Beis, Argyris Politis

## Abstract

The structural study of membrane proteins has traditionally relied on detergent-based extraction from cellular membranes. Although native-like reconstitution approaches have advanced, fully understanding membrane protein dynamics requires examining them within their native membrane environment. Hydrogen–deuterium exchange mass spectrometry (HDX-MS) is a powerful method for probing structural dynamics in reconstituted systems, but the presence of the lipid bilayer introduces considerable complexity, limiting broader adoption under physiological conditions. Here, we present the first fully automated HDX-MS platform incorporating a two-stage delipidation workflow. We applied this approach to monitor the dynamics of the ABC transporter MsbA in isolated inner membrane vesicles (IIMVs) from *Escherichia coli* through its ATPase cycle. IIMVs revealed distinct dynamic features within the nucleotide binding domains and substrate binding cavity, highlighting physiologically relevant motions not observed with detergent solubilised MsbA. This platform significantly advances HDX-MS and underscores the importance of studying membrane proteins in native lipid environments.

## Main

Rapid advances in structural biology and biophysical techniques have enabled the study of complex membrane bound macromolecular structures. A key challenge ahead is to probe the molecular structure and dynamics of these systems *in situ*. Hydrogen Deuterium Exchange coupled to Mass Spectrometry (HDX-MS) is a powerful analytical tool capable of reporting on conformational dynamics^1-4^, protein-protein/protein-small molecule interactions^5,6^, and folding/unfolding kinetics^7^, among others. As a bottom-up LC-MS technique, HDX-MS is more tolerant to complex samples when compared to other structural biology techniques, including exploring membrane protein dynamics within the context of their native-like lipid environment^8,9^. This capability is particularly enhanced when HDX-MS is coupled with post labelling delipidation strategies, which remove lipids that would otherwise interfere with downstream LC-MS analysis causing reduced enzymatic digestion efficiency, peptide ionisation suppression, increased spectral complexity, and impaired chromatographic performance^10-12^.While a few studies have demonstrated *in situ* HDX-MS of integral membrane proteins (IMPs)^13-17^, these approaches vary widely in methodology, including intensive sample clean up, precipitation of the protein, failure to cover transmembrane domains or focused on protein dense outer membranes.

The sensitivity of HDX relies on precise and robust experimental practices. The introduction of automation to HDX has improved the accuracy and efficiency of experimental handling. To date, the main strategy for lipid removal using automated methodologies involves the use of Zirconium Dioxide (ZrO_2_)-beads. ZrO_2_ binds the negatively charged head groups of phospholipids for effective sample clean up via filtration, or using a column packed with immobilised beads^9,10,12^. Although effective, this approach remains laborious, with reproducibility issues and non-specific binding resulting in low protein recovery. While automated lipid removal with ZrO_2_-bead filtration expanded this approach to simple protein-lipid samples, the use of this approach alone has drawbacks and limitations when handling complex lipid samples, such as isolated cell membrane vesicles for *in situ* HDX analysis, as it fails to remove other contaminants such as non-phospholipid membrane components, excess detergent used for membrane solubilisation, and other chaotropic agents that can be detrimental downstream. Adaption of an LC-MS system to include size exclusion chromatography (SEC) either prior to or after an online protease column has been shown to be highly effective for lipid removal^11^. The SEC column removes lipids and contaminants that are retained on column for much longer than both intact proteins and peptides. Previously, a manually operated valve was used to control the selective injection of protein and not lipid into the downstream LC-MS system. This had added benefits of being able to remove additional quench buffer contaminants e.g. urea. However, effective column regeneration and full automation of pre-digestion SEC lipid removal remained bottlenecks to widespread application to complex biological systems, such as lipid dense inner bacterial membranes.

As with all Gram-negative bacteria, the membrane envelope of *Escherichia coli (E. coli)* is composed of a double membrane. Unlike the outer membrane, that has a high protein: lipid ratio and extreme asymmetric lipid composition, the inner membrane of *E. coli* mainly consists of phospholipids (phosphatidylethanolamine (PE), phosphatidylglycerol (PG) and cardiolipin) with a 3:1 distribution of PE on each leaflet, depending on cell shape^18^. This dynamic, lipid rich membrane holds transporters of many families, the two most common being the ATP–binding cassette (ABC) and Major Facilitator Superfamily (MFS) transporters. Numerous studies have shown the importance of the lipids in the membrane in mechanism of both MFS transporters, e.g. sugar transporter / glucose transporter homolog XylE, and ABC transporters, e.g. MsbA, highlighting the necessity to characterise such transporters in the native lipid context to confirm or oppose existing models derived from work in mimetic reconstitution environments^19,20^. Both MsbA and XylE proteins have been well biochemically characterised and serve as model systems for their mammalian homologs.

MsbA is an essential homodimeric ABC exporter responsible for flipping core lipid A - the precursor of lipopolysaccharide (LPS) - across the cytoplasmic membrane for outer membrane biogenesis in Gram-negative bacteria. MsbA consists of two identical half-transporters; each comprising a six-helix transmembrane domain (TMD) and a cytoplasmic nucleotide binding domain (NBD). ATP binding and hydrolysis at the NBDs drive large-scale conformational changes that are coupled to the TMDs. These changes reorient a large cavity between inward facing (IF) and outward-facing (OF) conformations with dimerized NBDs sandwiching two ATPs at the dimer interface^21,22^. MsbA has been used as a model system to characterise the conformations along the transport cycle using structural (cryo-EM and X-ray crystallography), biophysical (EPR) and biochemical (mutagenesis and transport assays) studies. These studies have been performed in either detergent or lipid mimetics (nanodiscs) or whole cells, yet they all display inconsistent results ^21-24^.

To address the physiological relevance of transporter conformation it is necessary to study these systems within their native membranes. Here, we present the first fully automated HDX-MS platform that utilises a dual-mode delipidation strategy, for removal of lipid and other contaminants prior to LC-MS analysis. Our platform relies on ZrO_2_ delipidation, performed similarly as described in Anderson *et al*.^12^, and subsequent SEC delipidation performed on-line using a novel three valve and three LC pumps configuration, on a commercially available system. While the ability to use standalone ZrO_2_ or SEC increases system versatility, the use of ZrO_2_ and SEC in tandem, henceforth referred to as dual-mode delipidation (DMD) (**Fig. 1**), substantially increases lipid removal efficiency beyond either ZrO_2_ or SEC alone, with minimal impact on deuterium back-exchange measurements. These advancements allowed probing the dynamic flexibility of the ABC transporter MsbA within physiological membrane, capturing previously unreported conformations at both the TMDs and NBDs. With our advancements to the classical HDX-MS membrane protein workflow we provide a robust and detailed approach to directly compare conformational dynamics within native membranes to lipid mimetics, that can be applied to a range of IMP superfamilies. We envision that more widespread adoption of this approach will allow more robust and reproducible analysis of integral membrane proteins in native lipid environments by HDX-MS in the near future.

**Figure 1.**
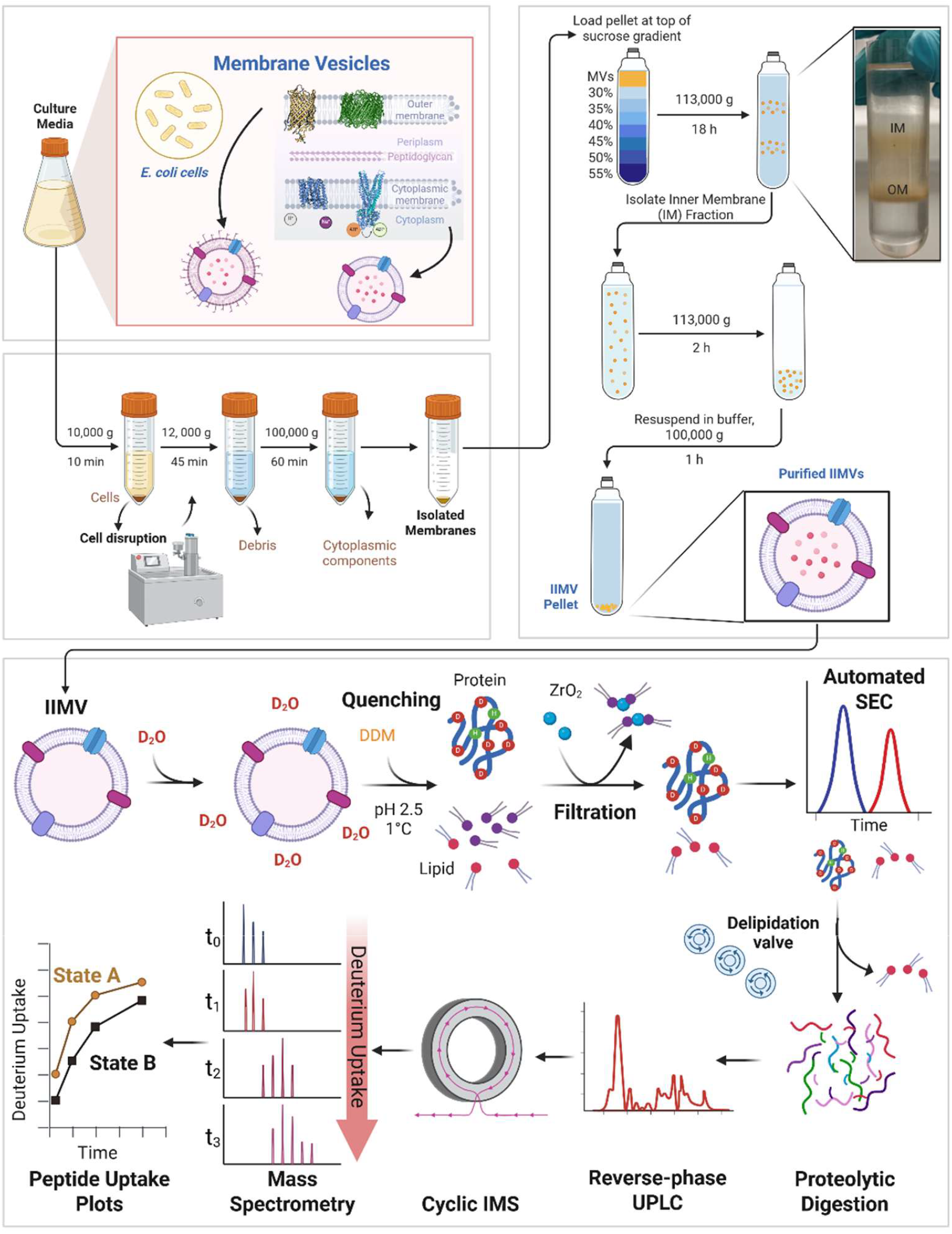
In-situ HDX-MS workflow with integrated dual-mode-delipidation. Protein-lipid samples (i.e., IIMVs) are first incubated in deuterated buffer for deuterium incorporation. The exchange reaction is then quenched by dropping pH and temperature to 2.5 and 0-1°C, respectively, which is coupled to DDM-mediated disassembly. ZrO_2_ beads are then added to selectively bind to phospholipids, with ZrO_2_-lipid particulates being subsequently removed via membranous filtration. The filtrate is then subjected to rapid size exclusion chromatography (SEC), which allows a delipidation valve to direct faster eluting protein for proteolytic digestion and later eluting lipid to waste. Peptides are then analysed via LC-MS to detect deuterium uptake at peptide level resolution and compared between states.

## Results

### Fully Automated Dual-Mode Delipidation

To perform fully automated HDX-MS on lipid dense protein samples we reconfigured a dual head LEAP HDX robot with a filtration system (Trajan Scientific, USA) to perform all sample handling and mixing, ZrO_2_- and SEC-mediated delipidation, without affecting analytical throughput (**Fig. 1**). This primarily involved adaption of the fluidic system to allow for fully automated SEC delipidation and subsequent SEC column cleaning/regeneration (see **Methods, Fig. S1**). To demonstrate the capabilities of the newly developed DMD-HDX-MS workflow and to compare its delipidation efficiency and protein recovery to autonomous BSA-washed ZrO_2_ (wZrO_2_) or SEC delipidation HDX-MS workflows, we employed 1-palmitoyl-2-oleoyl-sn-glycero-3-phosphocholine (POPC) and Bovine Carbonic Anhydrase (BCA) as model systems (**Fig. 2**). Quantification of residual POPC and BCA recovery in samples after delipidation was achieved via direct MS measurements. While these analytes differ from those expected in isolated membranes samples, their simplicity allowed determination of the systems overall efficiency.

**Figure 2.**
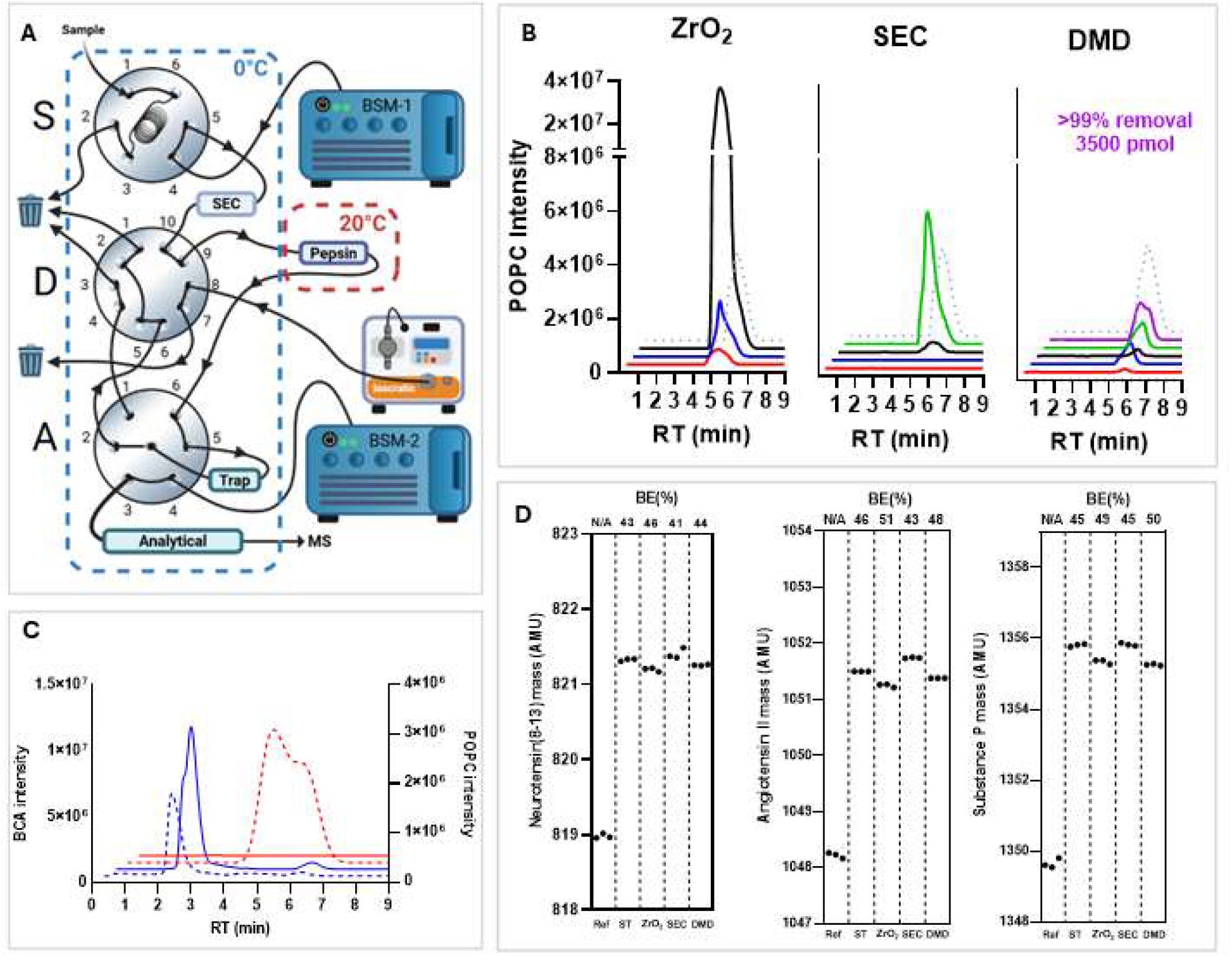
Characterisation of automated ZrO_2_, SEC and DMD delipidation. (**A**) Schematic of Trajan and Waters LC fluidic system with valve reconfiguration for fully automated DMD. The sample (S), delipidation (D) and analytical (A) naming conventions are shown with LC and waste bottle connections. The full 8-step valve cycle is shown in **Fig S1**. (**B**) Extracted ion chromatograms (EICs) of 10 (red), 100 (blue), 1000 (black), 2500 (green) and 3500 (purple) pmol POPC mass experiments after delipidation via wZrO_2_, SEC and DMD. Dashed lines represent 50 pmol of POPC injected without delipidation. Post-delipidation POPC quantification was achieved using the detector calibration curve in **Fig S2**. (**C**) EICs from BCA (blue) and POPC (red) intact mass experiments. Dashed lines represent EICs pulled from the same experiment (60 pmol BCA + 3500 pmol POPC) and block lines represent EICs pulled from the same experiment (60 pmol BCA). (**D**) Nested plots showing ion mass (Da) per delipidation type over triplicate experiment. The delipidation approach used and calculated average back-exchange (%) are specified at the bottom and top of each channel, respectively.

Overall, all three delipidation methods were able to remove the model lipid POPC up to a different degree (**Fig. 2B and Table S1**). wZrO_2_ delipidation alone removed up to 100 pmol of POPC with 73% efficiency, while SEC mediated delipidation successfully removed 98% of 2500 pmol POPC. As expected, DMD workflow outperformed both individual approaches, removing 99% of 3500 pmol POPC, making this approach the only one able to handle samples with high lipid content. Similarly, protein recovery using 60pmol BCA was calculated for all three methods, resulting in 107± 24% recovery for wZrO_2_, 61± 4 % for SEC and 48± 1% for the DMD workflow (**Fig. S3**). To ensure the automated HDX-MS platform provided selective removal of lipids, 60 pmol BCA and 3500 pmol POPC was analysed together using DMD and compared to 60 pmol BCA alone without delipidation (**Fig. 2C**). Both delipidation efficiency and protein recovery remained relatively unchanged (97% efficiency and 44% recovery) as in initial experiments. All in all, these results demonstrate that automated DMD can remove large amounts of lipids while maintaining protein quantities sufficient for subsequent LC-MS analyses. We note that the quantity of ZrO_2_ was not optimised to improve lipid removal here, instead we used the established concentration for lipid removal from nanodisc samples^8^.

When introducing additional steps into the HDX-MS workflow, it is critical to assess its suitability through quantification of hydrogen/deuterium back-exchange^25,26^. Compared to the system without any delipidation steps, that gave a back exchange measurements of 44± 2%, standalone SEC and ZrO_2_ resulted in an average of 42± 3% and 48± 2% respectively using three model peptides (**Fig. 2D**). When combined in DMD, resulting back exchange was 47± 3%. While back-exchange increased compared to no delipidation, the difference in back exchanges is under <5% and remain within previously established community recommendations^26^.

Overall, the fully automated DMD HDX-MS workflow enables highly efficient lipid removal while maintaining sufficient protein recovery for downstream LC-MS analysis and introducing only minimal increases in back-exchange, demonstrating its suitability for robust *in situ* HDX-MS applications.

### Peptide positive identifications using DMD workflow in Isolated Membrane Vesicles

To assess the DMD capabilities, positive peptide identifications via HDMS^E^ fragmentation using detergent solubilised protein and isolated inner membrane vesicles (IIMVs) were compared under identical LC-MS conditions. To ensure workflow robustness, separate preparations of IIMVs from *E. coli* containing overexpressed MsbA or XylE were utilised. IIMVs were harvested directly from fractionated cell pellets (**Fig. 1**), using high-pressure lysis to invert the membrane orientation, as such the cytoplasmic protein domains were most exposed to external buffers.

Successful peptide identifications rely on achieving sufficient target protein concentrations, efficient delipidation and effective membrane solubilisation under quench conditions, among others. Coupling of DMD and HDX-MS for IMPs in complex lipid environments requires measurement of protein:lipid ratios to ensure detectable levels of target protein whilst maintaining efficient delipidation. MsbA represented ∼6.5% of the total protein within the isolated membranes (**Fig. S4**) while quantification of total lipid content gave a lipid concentration of ∼0.23 mM (**Fig. S6**). Similarly, protein and lipid content of XylE IIMVs gave ∼22% of the expressed membrane protein and ∼0.75 mM of total lipid concentration, respectively (**Fig. S5-6**).

Peptide identification of non-deuterated samples under DMD-LC-MS conditions was optimised for target protein: lipid: detergent molar ratios determined at ∼1:200:5000, with a maximal lipid load of ∼3700 pmol. High number of positive identifications was achieved; 98.3% coverage across 263 peptides and 93.9% coverage across 100 peptides for MsbA and XylE respectively (**Fig. S7**). In comparison to IIMV data, MsbA purified from the membranes into DDM, subjected to DMD-LC-MS produced a sequence coverage of 97.1% coverage with 424 peptides, suggesting that the decrease in peptide identifications results from increased MS spectral crowding and lower peptide peak intensities in the IIMVs rather from sample loss during DMD. These data demonstrate the reproducibility, reliability and adaptability of the presented DMD platform, that is not restricted to a specific membrane protein family.

### *In vitro* vs *in situ* dynamics of the ABC transporter MsbA

For rigorous validation of our DMD platform, we monitored the conformational dynamics of the MsbA transporter, in its apo and inhibited post-hydrolytic state. As a type IV ABC transporter, MsbA undergoes large structural rearrangements during its ATP hydrolysis cycle. Such rearrangements result in significant changes in protein dynamics that can be well followed by HDX-MS^15,27-29^.

We carried out HDX-MS on MsbA IIMVs and DDM-solubilised MsbA in the absence of nucleotides (apo; only Mg^2+^) and in the presence of ATP + vanadate (Vi). Following manual data curation of deuterated peptides effective sequence coverage of the MsbA in IIMVs was 60.3% across 64 peptides with 1.93 redundancy (**Fig. S7**). Coverage was retained for the vast majority of TMD and NBD features were, although part of the α-helical subdomain of the NBD and most of transmembrane helix (TMH) 2 and 4 were lost.

*In vitro* MsbA solubilised in DDM in its apo state exhibited the lowest levels of deuterium uptake in the TMHs, with TMH-1, -3 and -5 exhibiting almost no deuterium uptake at all (**Fig. 3**). The overall pattern of intrinsic dynamics of MsbA in IIMVs was highly comparable to the results observed for MsbA in DDM in the apo state (**Fig. 3**). The major differences in dynamics within the TMDs appear at the cytoplasmic extensions of TMH1, TMH3 and TMH5 and around the lipid substrate binding site, where the detergent protein surprisingly has greater uptake of deuterium. The apo wide IF structure of MsbA, while observed in IIMVs by Galazzo *et al*^*19*^, is found in addition to narrower IF species. It is likely this mixture is present in our HDX samples contributing to reduced uptake in IIMVs, but they do not exist distinctly enough to result in multimodal distributions.

**Figure 3.**
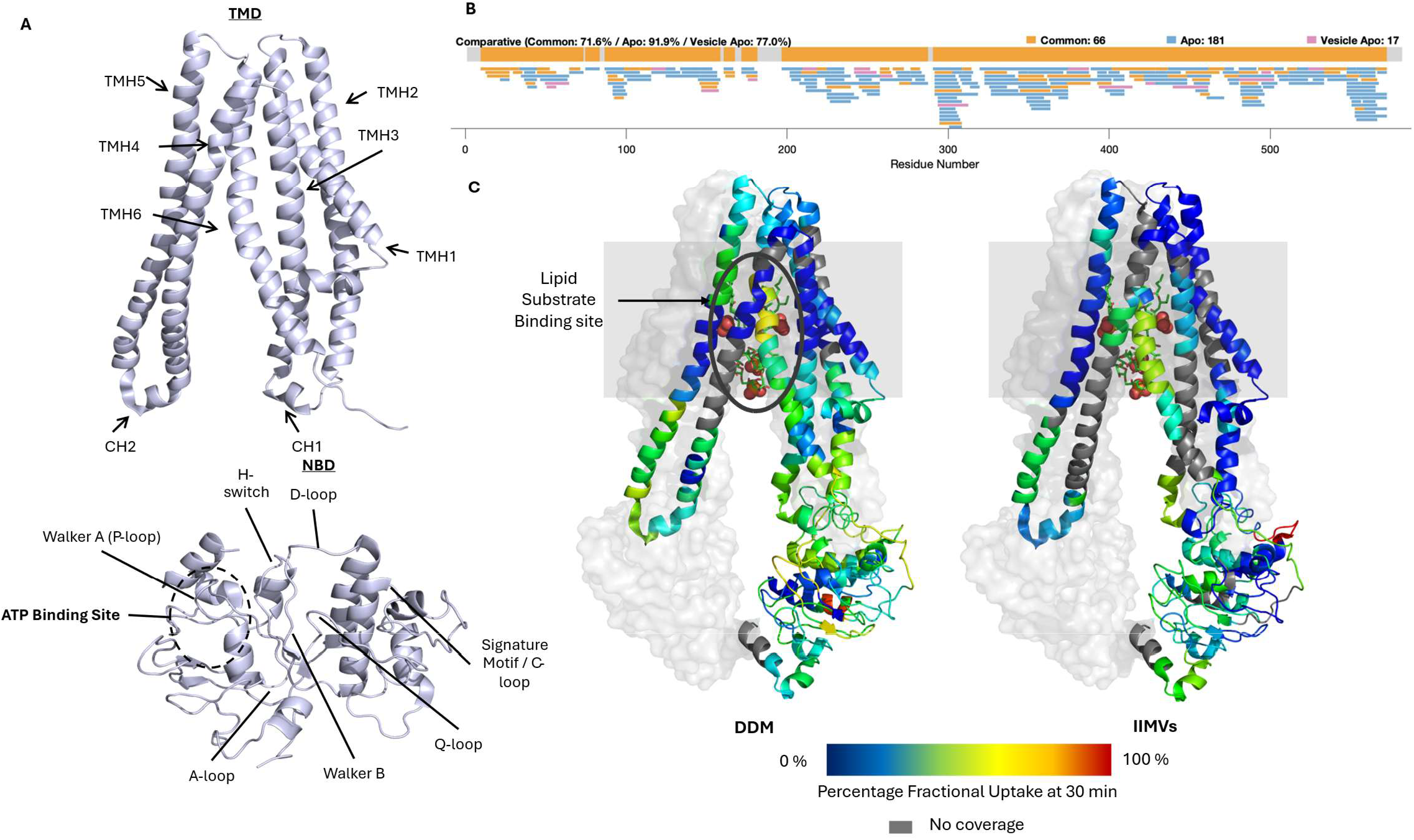
Peptide mapping and Intrinsic Dynamics of MsbA in detergent and IIMV lipid environments under DMD-HDX-MS conditions. **(A)** MsbA domain structure in the IF_narrow_ conformation highlighting domain features. The transmembrane domain (TMD) is shown on top and the nucleotide binding domain (NBD) shown below. **(B)** Peptide coverage maps of apo MsbA in DDM and IIMV. **(C)** Percentage Deuterium Uptake relative to maximum theoretical uptake mapped onto the structure of MsbA IF_narrow_ conformation (PDB: 5TV4^21^) with one monomer represented as helical cartoons and the other as a surface structure. Lipid substrate is shown as sticks. Regions without sequence coverage are coloured grey. Percentage fractional deuterium uptake from MsbA DDM and IIMV HDX-MS at 30 min timepoint is heat-mapped onto the structure. Deuterium uptake plots of peptides can be found in **Supplementary Info**.

In line with homologous studies, addition of ATP + Vi resulted in protection from exchange across the transporter with a large majority of the DDM solubilised protein showing reduced deuterium uptake. HDX-MS *in situ* corroborated this stabilisation of the transporter upon addition of ADP*Vi seen in DDM, particularly at the NBD dimerization interface and the core TMHs (**Fig. 4B and 5**). Additionally, stabilisation of both coupling helices (CHs) supports NBD dimerization driving TMD conformation change through stabilisation of an NBD-TMD communication relay at the CHs. Deprotection was also observed within the extracellular domain, however only between TMH 1 and 2. In contrast to the DDM, IIMVs show a larger number of deprotected peptides at the surface of the NBDs and within the membrane of TMH4 and TMH5 that appear not to be associated with NBD dimerization. Due to the presence of the lipid bilayer changing the overall dynamics of the protein, direct comparison of peptides between DDM and IIMV data was minimised. Six peptides were selected for direct comparison that demonstrated similar uptake behaviour in the apo state, two from the TMD and 4 from the NBD. Deuterium uptake plots from the NBDs demonstrate that at the ATP binding site, including peptides containing the Walker A, Signature motif and the H-loop, protection from exchange following ADP*Vi binding is clearly observed both in DDM and IIMVs, as anticipated (**Fig. S9**).

**Figure 4.**
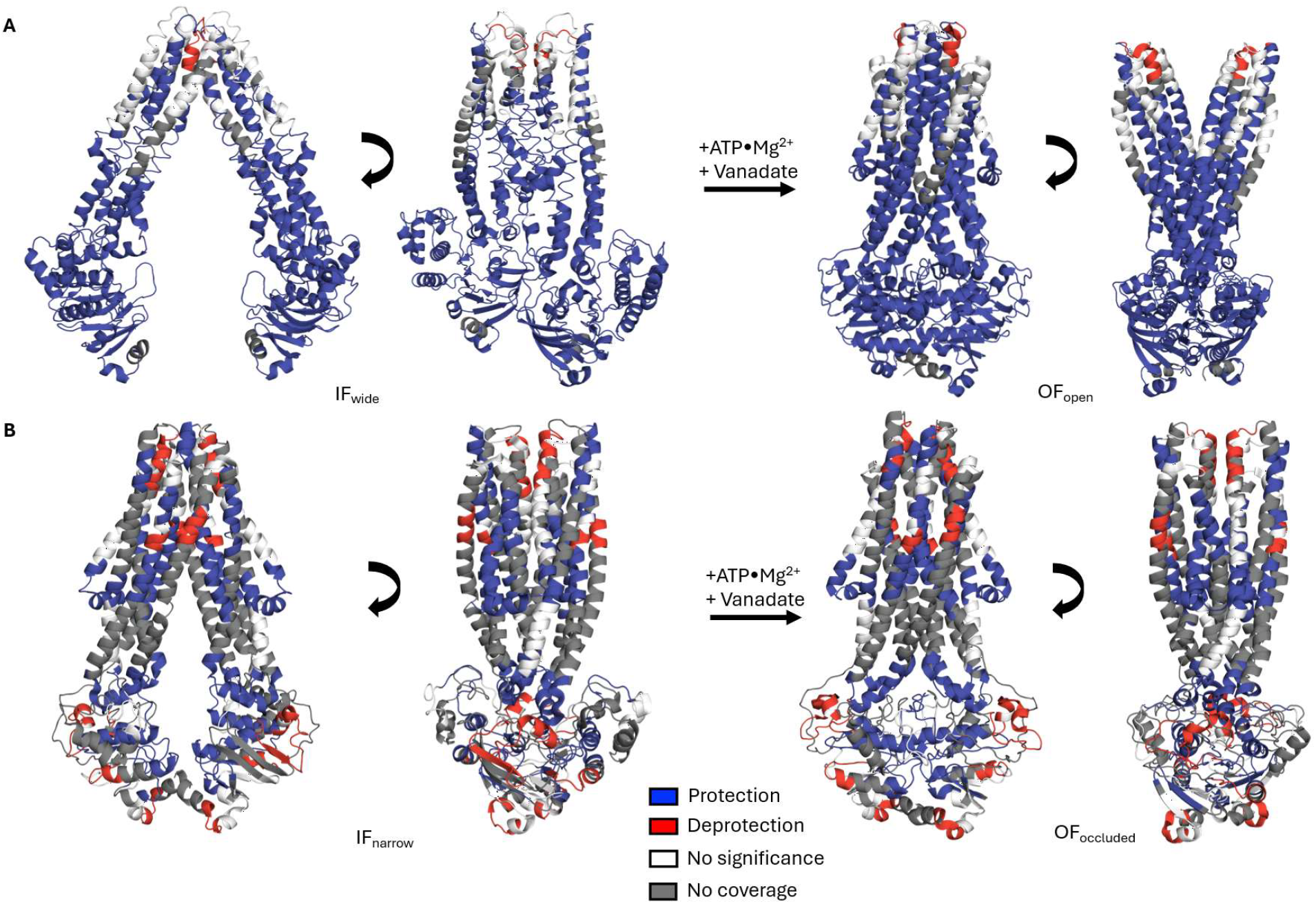
ATP hydrolysis induced changes in MsbA Dynamics in vitro and in situ. (**A**) MsbA ΔHDX of apo and ADP*Vi trapped states in detergent micelles, mapped onto the conformational transitions between inward-facing (IF) (PDB:9FUQ^23^) and outward-facing (OF) (PDB: 3B60^22^) structures determined in DDM. IF-wide structure is depicted as the apo DDM cryo-EM structure while OF-open structure is depicted with the DDM AMP-PNP bound crystal structure, which maps identically to the DDM ADP*Vi bound structure but with higher resolution. ADP*Vi for formed following incubation with ATP and vanadate. Hydrolysis of ATP to ADP releases a phosphate that is replaced by vanadate, trapping the structure in a post-hydrolytic state. (B) MsbA ΔHDX of apo and ADP*Vi trapped states from Isolated Inner Membrane Vesicles (IIMVs). Differential uptake is mapped onto the IF-narrow structure (PDB:5TV4) and the OF-occluded (PDB:5TTP^21^) from MSP1E3D1 nanodiscs with E. coli polar lipid extract bilayers. In all panels, peptides showing less deuterium uptake (protected from exchange) are shown in blue while peptides showing increased uptake (deprotected) are shown in red. Peptides with no significant difference in uptake are shown in white while regions where peptides were not recovered are shown in grey. Data represents four technical replicates for all states.

**Figure 5.**
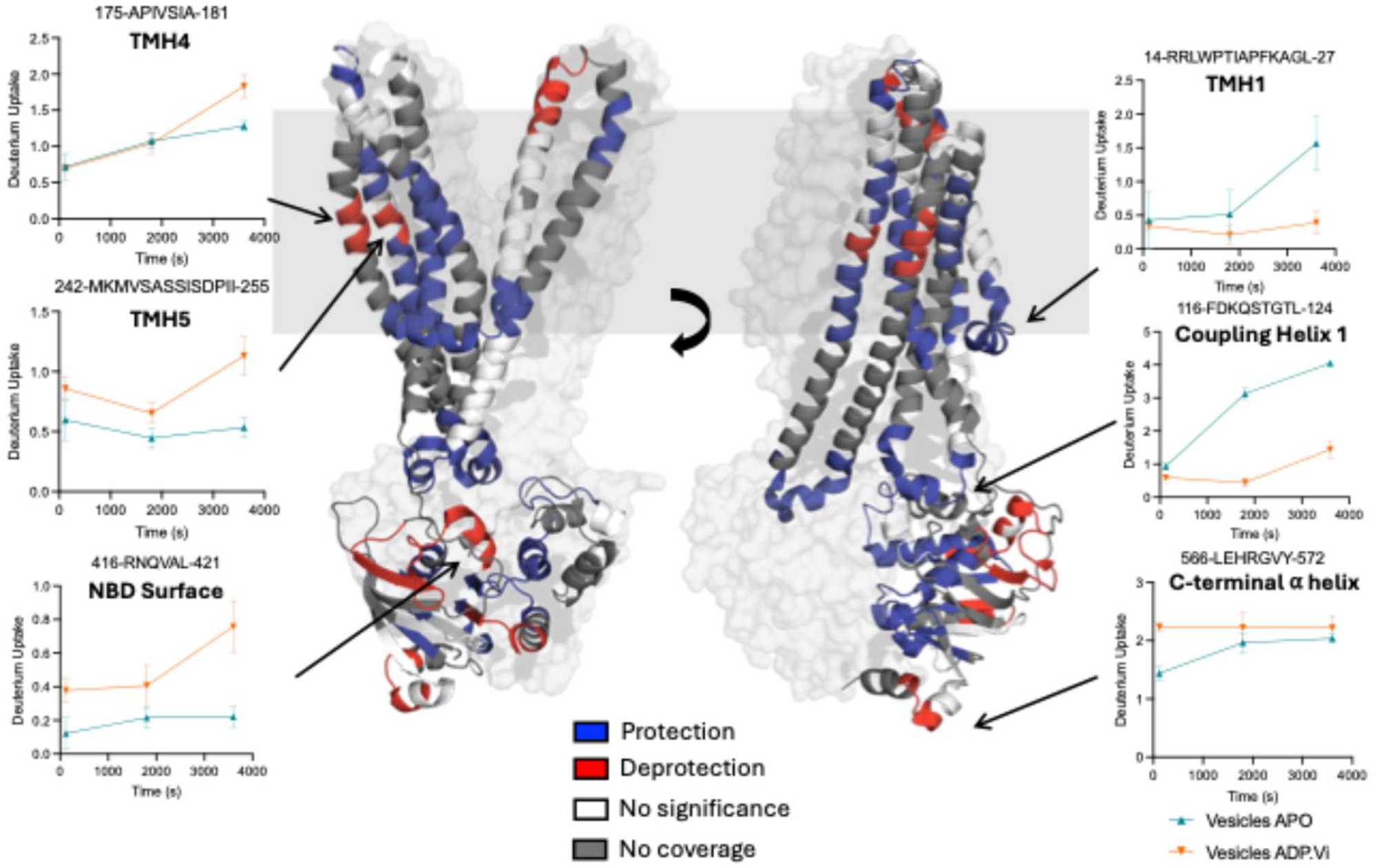
Membrane embedded MsbA has an open OF post-hydrolytic conformation. ΔHDX shown on OF_open_ (PDB: 3B60^22^) conformation. For clarity, one monomer is shown as a transparent surface. Statistically significant differences from Apo vs ADP*Vi ΔHDX-MS are mapped onto both models with protection (blue), deprotection (red), no change (white), no coverage (grey). The same data is represented in Fig. 4. All uptake plots can be found in **Supporting Data**.

## Discussion

*In situ* HDX-MS study of membrane proteins has employed a broad array of approaches with success in isolated outer membrane vesicles, mitochondrial membranes and within cell^13,14,16,17,30^. However, these manual methodologies are labour intensive, introduce opportunity for measurement error between replicates and have low coverage of transmembrane peptides, increasing the likelihood of obscuring subtle differences in dynamics across the whole protein. As a result, it is not unsurprising that *in situ* studies are underrepresented in the literature, despite introduction over a decade ago. Additionally, coverage of integral membrane proteins has remained a challenge across the field. Improved instrumentation and automation platforms have transformed the feasibility of IMP HDX-MS^31,32^.Here, we report the first fully automated HDX-MS platform utilising a two-stage ZrO_*2*_ and SEC-mediated delipidation in tandem for the analyses of complex protein-lipid assemblies. We envision this platform will increase the feasibility and reliability of running HDX-MS with mixed protein-lipid samples, particularly those with increased complexity and high protein: lipid ratios. The ability to use either standalone delipidation approach increases system versatility and allows better tailoring of delipidation approaches on a sample-by-sample basis. Our ∼70% coverage over the entire MsbA transporter under HDX-MS is significant improvement on previous studies. Further optimisation of DMD could further improve delipidation efficiency beyond what is demonstrated in this publication, along with being tailored for more complex membranes e.g. isolated mammalian membranes. For example, the adaption of the system to nano-flow LC is likely to both improve the coverage and reduce the sample requirement therefore reducing the lipid content for removal^33^. For samples not requiring handling of high lipid content we recommend the SEC delipidation workflow over wZrO_*2*_ due to greater robustness and reliability when automating data collection. Moreover, we recognize that sample preparation parameters, such as target protein expression and the protein: lipid ratio, have a significant influence on data quality and experimental outcome. As a result, we recommend establishing standardised reporting of these values in future publication methods, as exemplified in this work, to aid experimental optimisation processes in the future.

The work presented here demonstrates that careful platform design can decipher protein dynamics *in situ* that are not observable in detergent or lipid mimetic reconstituted environments. MsbA sensitivity to reconstitution environment has allowed the dynamic nature of the IF conformation to be probed by cryo-EM, EPR, smFRET and *in cellulo* nanobody-assisted EPR^19,23,34,35^. Contrastingly, rearrangements of the extracellular loops of the TMD cannot be resolved with distance measurements and have failed to be captured with other biophysical techiniques^36^. Debate still remains on the physiological relevance of an OF_occluded_ with fully dimerized NBDs and a closed periplasmic TMD or the OF_open_ with dimerised NBDs but wide opening to the periplasm found in differing reconstitution environments^21,22^ (**Fig. 4)**. In line with previous studies, our platform showcases large changes in conformational dynamics during ATP hydrolysis, both *in vitro* and *in situ*, with stabilisation of the transporter seen across the protein, particularly around the interface of the nucleotide binding domains, the CHs and extending into the substrate binding core (**Fig. 4 & 5**). This dramatic stabilisation is particularly obvious within DDM, which may reflect the broad distribution of conformations and wider opening of the central cavity seen structurally in the apo state (**Fig. 4A**). Our data from DDM and IIMVs find deprotection from exchange extending into the TMD cavity between TMH5 and TMH6, at p<0.01, in the DDM state and deprotection between TMH1 and TMH2, at p<0.01, in IIMVs, both suggesting an OF_open_ conformation. The TMH1-2 loop has been identified as a key feature of multidrug resistant type IV ABC exporters, varying in size and shape to allow for diverse substrate release, while TMH6 rearrangements in MsbA are hypothesised to encourage substrate release. Similar increases in helical flexibility have been observed for MsbA homologs, BmrA and P-gp upon stabilisation of the OF conformation. In particular, residues 45-56, showing higher deuterium uptake in the IIMV ADP*Vi bound state, are overlapping with deprotection observed in nanodisc BmrA stabilised by Vi trapping^37^. The number of peptides showing increased deuteration is lower in MsbA compared to both BmrA and P-gp, only one or two peptides show increase in either TMH1 or TMH6 while homologous proteins shown increased exchange at TMH1, 2, 3, 4 and 6^28,29,37,38^. These data suggests that while the fully occluded conformation seen in MsbA nanodisc structures is unlikely, both in DDM and IIMVs, the degree of opening under our HDX conditions may not be as wide as shown in structures from Ward *et al*. and Zhang *et al*. due to the extension of protection into the TMD cavity (**Fig. 4**)^22,24^.

The differences observed between DDM and IIMV data, may be associated with different OF conformations that can be attributed to the presence of lipid, both in stabilising the transporter but also acting as a substrate for MsbA. Assuming that within IIMVs some MsbA may be substrate bound, the change in protein dynamics following ATP hydrolysis may also represent a post-hydrolysis and post-lipid translocation state. We cannot differentiate under our current experimental parameters between apo and lipid bound MsbA within IIMVs. The presence of substrate results in stimulation of MsbA ATP hydrolysis activity and while a lipid bilayer may stabilise the apo state, it may also result in a more dynamic substrate binding cavity, which may explain the unexpected increased uptake in the TMD within IIMVs^39^.

*in situ* HDX-MS provides peptide level resolution into conformational dynamics, unobservable in other techniques. While tracking the conformational dynamics of dimerization and transition to an OF state is clear, the surface of the NBDs that forms no interface, i.e. neither intra- or inter-monomer interactions, shows unexpected increases in deuterium uptake upon stabilisation by Vi inhibition solely in the IIMVs (**Fig. 4 & 5**). This deprotection extends to destabilisation of the dynamic C-terminal α-helix in the ADP*Vi bound state within the IIMVs (**Fig. 4 & 5**). The C-terminal α-helix is believed to be a stabilizing structural feature of ABC exporters that may interact with the Walker A domain during ATPase activity^40-42^. The presence of the membrane does not appear alter dynamics associated with nucleotide hydrolysis at conserved ABC motifs (**Fig. S9**). In the post-hydrolytic state, perhaps accompanying substrate translocation, destabilization at the C-terminus of the NBDs may encourage NBD separation following ATP hydrolysis and substrate release to return the transporter to an IF conformation that results in increased dynamics at the NBD outer surfaces. Additionally, as *in situ* studies are carried out in the presence of the inner membrane proteome, interactions partners present in the membrane may reduce uptake in these regions in an apo state, potentially involved in regulating futile ATP hydrolysis. These features have not previously been observed in any *in vitro* studies however, further exploration is needed to capture the full *in situ* biochemistry of MsbA.

The fully automated DMD platform presented here is a robust platform able to identify and follow a high number of peptides from IMPs, with limited back exchange and no chromatographic fouling. Probing the dynamics of an ABC transporter *in situ* highlighted previously uncharacterised dynamics within the NBDs and validated the physiological presence of an OF_open_ conformation. This platform is adaptable to further development with sub-zero or nano-flow LC that may enhance peptide recovery and allow expansion to proteins that express at lower levels while also improving recovery of deuterated peptides. Thus, the platform represents a notable advancement in automated LC-MS methodology with potential to enable the high throughput analysis of IMPs in complex lipid environments.

## Methods

All materials and additional experimental details can be found in the Supporting Information.

### HDX-MS

HDX-MS experiments were performed on a SELECT SERIES cIM QTOF (Waters) mass spectrometer using single pass ion mobility as described previously^32,43^. All sample handling was carried out using a dual head Trajan LEAP HDX automation system equipped with lipid filtration system.

For apo states, deuteration labelling was initiated by diluting 5 μL of either 28.4 μM MsbA in 0.2% DDM or 2.34 ± 0.71 mg/mL MsbA IIMVs in 95 μL of D_2_O labelling buffer (20 mM Tris-HCl, 150 mM NaCl, pD 7.0). Samples were labelled for 2, 30 and 60 minutes at 20°C. Subsequently reactions were quenched by the addition of 100 μL of quench buffer (100 mM Glycine-HCl, 0.2% DDM, 4 M Urea, pH 2.5) precooled to 1°C. Following two mixing strokes, sample was transferred to the filter base of the nanofilter vial (Thomson Instrument Company, Carlsbad, CA), and wZrO_2_ mediated delipidation was performed at 4°C. Each base contained 10 μL of pre-dispensed 300 mg/ml ZrO_2_ coated silica beads in 0.1% formic acid in water, pre-washed with BSA as previously described^44^. Sample was filtered and 190 μL of filtrate was injected into a 250 μL sample loop. For ADP/Vi states, purified protein or IIMV stocks were diluted to the same protein concentrations as above with equilibration buffer (20 mM Tris-HCl, 150 mM NaCl, pD 7.0) supplemented with 10 mM ATP, 2 mM Vanadate and 10 mM MgCl_2_.

Sample then underwent SEC delipidation at 0°C using a Waters BEH SEC guard column (125 Å, 1.7 μm, 4.6 mm x 30 mm, 1–80 K) followed by on-line digestion at 20°C with a self-packed column (30 × 2.1 mm) containing immobilized porcine pepsin (Thermo Scientific, Leicestershire, UK) using 0.1% formic acid in water (mobile phase A) at 200 μL/min. After 160 sec, SEC valve switching diverted the lipid elution to waste, while maintaining flow to complete protein digestion and peptide trapping and further desalting for 50 sec on a Waters BEH C18 (2.1 × 5 mm, 1.7 μm) VanGuard pre-column (Waters Corporation, Wilmslow, UK) at 200 μL/min. Peptides were eluted and separated on a Waters BEH C18 analytical column (1.0 × 100 mm, 130Å, 1.7 μm) with a 13-40% linear gradient of mobile phase B (0.1% formic in acetonitrile) over 8 min at 40 μL/min followed by immediate increase to 85% B, maintained there for two minutes before returning to 8% B. To avoid excessive carry-over, an additional 5 minutes at 85% B were used after each gradient separation. All chromatography steps were performed at 0°C to decrease back-exchange. Peptide masses were measured using positive ESI HDMS acquisition.

For non-deuterated controls, equilibration buffer was used in place of labelling buffer with identical chromatography, but MS data was collected using HDMS^E^ with a collision energy ramping from 25– 45 V. Identical workflow was used for XylE IIMVs experiments where a stock solution of 11.0 mg/mL IIMVs was used. All HDX-MS experiments were performed in quadruplicate (n=4).

To minimise peptide and lipid carry-over, each sample injection was accompanied by three 100 μL injections of pepsin wash (100mM K_2_HPO_4_, 2 M GdnHCl, 5% Acetonitrile, pH 2.5) and two 100 μL injections of SEC wash (6 M GdnHCl, 25% MeOH, pH 2.5). Prior to pepsin washes, the SEC column was washed with 25% methanol for 3 min at 200 μL/min. Additionally, each sample run was followed by a cleaning run. Each cleaning run consisted of 3 sawtooth gradients (8-85%) through the trap and analytical column, while simultaneously the SEC column was washed with four 2 min cycles of 20 or 25% 50:50 IPA:H_2_O containing 0.1% fos-choline-12 for a total of 25 minutes per cleaning run. An additional two 100 μL injections of SEC wash over the SEC column and three 100 μL injections of pepsin wash over the SEC and pepsin column were also used during the cleaning gradient. Peptide carry over was assessed by immediately running a blank after an HDX-MS experiment. Carry-over was considered acceptable if peptide intensity was below 10% of the previous experiment.

Resulting peptides from HDMS^E^ were identified using ProteinLynx Global Server (v 3.0.3 from Waters™) and filtered using LARS 2.0^45^ using the following parameters: minimum intensity of 1000, minimum sequence length of 4, maximum sequence length of 25, minimum products of 3, minimum products per amino acid 0.11, minimum score of 6.62 and maximum MH^+^ error of 5 ppm. Final peptide lists from DDM and IIMV peptide mapping were combined and imported into HDExaminer (v 3.5.0 RC5) for manual curation. Statistical analysis was carried out with Deuteros 2.0^46^ using hybrid significance tests with a significant level of 0.01^47^. Figures were prepared using PyMOL Molecular Graphics System (Version 3.0 Schrödinger, LLC). To allow access to the HDX data of this study, HDX data summary table (**Supplementary file HDX Table S1**), HDX uptake data table (**Supplementary file HDX Table S2**) and all uptake plots (**Supplementary file Uptake plots**) are included in the supporting information as per consensus guidelines^26^. Additionally, all mass spectrometry data have been deposited to the ProteomeXchange Consortium via the PRIDE partner repository with the dataset identifier XXXXXX.

## Supporting information

Uptake Plots - HDX Data

## Acknowledgements

D.G. and A.P. acknowledge receiving PhD studentship funding from The University of Manchester BBSRC DTP. A.P. was also supported by an EPSRC Research Fellowship (EP/V0117151/1) and BBSRC grants (BB/V006487/2, BB/Y004981/1, and BB/X018326/1). The authors acknowledge the MRC Equipment Grant MR/X013030/1. AP acknowledges support by the European Union via the Horizon Europe ERA Chair “MASSTRUCT” Project, number id 101183630.

## Authorship contribution statement

D.G. J.P.R.P., C.G. and A.P. conceptualised and designed the research. C.G., D.G. and J.P.R.P. set up instrumentation and designed experiments. J.P.R.P and D.G. performed intact mass and back-exchange evaluation. C.G. performed all bottom-up HDX-MS experiments for MsbA. J.P.R.P carried out peptide mapping for XylE. S.I. and K.B. provided IIMV MsbA samples. C.G. performed MsbA purification and performed protein and phospholipid quantification assays. J.P.R.P. and C.G. analysed and curated all data. C.G. wrote the paper with input from all authors. A.P. supervised and acquired funds for the research.

## Declaration of competing interest

The authors declare no completing interest.

## Data availability

## Notes

### Competing Interest Statement

The authors have declared no competing interest.

