## Supplementary File for "An Automated HDX-MS Platform for *in situ* characterisation of Membrane Proteins"

### Supplementary Materials and Methods

#### Materials

1-palmitoy-2-oleoyl-glycero-3-phosphocholine (POPC) and n-Dodecyl-β-d-maltoside (DDM) were purchased from Avanti Polar Lipids (Alabaster, AL, USA). Bovine carbonic anhydrase (BCA), bovine serum albumin (BSA), Optima grade 0.1% formic acid in water and 0.1% formic acid in acetonitrile blends were purchased from Fisher Scientific (Leicestershire, U.K.). All other reagents were purchased from Merck Life Science (Gillingham, U.K.) unless otherwise noted.

#### MsbA expression, IIMV preparation and purification

WT MsbA from *E.* coli was overexpressed in *E. coli* BL21 (DE3) cells transformed using a pET-28b vector containing a C-terminal His10-tag and a Tobacco Etch Virus (TEV) protease site. The cells were cultured in LB at 37°C until OD600 reached 1.0, followed by induction with 0.5 mM IPTG. The cells were grown for a further 16 hours at 25°C. The cells were then harvested by centrifugation at 5,000 x g for 10 min at 4°C. The harvested cells were resuspended in 20 mM Tris-HCl pH 8.0, 20 mM NaCl, 0.1 mg DNase, 1 mM $MgCl_2$, 1 mM 4-benzenesulfonyl fluoride hydrochloride and 5% glycerol at 4°C, and then disrupted using a cell disruptor at 25 kpsi (2 passes) at 4°C. Cell debris and unbroken cells were removed by spinning the lysate at 12,000 x g for 30 min at 4°C. Total membranes were then collected by ultracentrifugation at 100,000 x g for 1 hour and resuspended in membrane buffer (MB) (20 mM Tris-HCl pH 7.5 and 0.5 mM EDTA) at 4°C. To isolate the IIMVs from the total membrane extract a sucrose gradient was prepared in 65 mL ultracentrifuge tubes containing 10 mL layers of: 55, 50, 45, 40, 35, and 30% (w/w) sucrose in MB, and 3.5 ml of total membranes (supplemented with 25% (w/w) sucrose) was layered on top (**Fig. 1**). After ultracentrifugation at 113,000 x g for 18 hours at 4°C with minimal acceleration and no breaking, the inner membrane (IM) fraction from the 35-40% gradient was collected and back diluted in 5 CV of MB. The isolate IM vesicles (IIMVs) were then harvested by ultracentrifugation at 113,000 x g for 2 hours and at 4°C and resuspended in 20 mM Tris-HCl (pH 7.0), 150 mM NaCl. Pellets were washed a final time in 20 mM Tris-HCl (pH 7.0), 150 mM NaCl, 10% (v/v) glycerol before storage. The presence of MsbA inside the IIMVs was verified by SDS-PAGE and western blot. Aliquots were then prepared and stored at -80°C.

Purification of MsbA from IIMVs was achieved by solubilising total membranes at 5 mg/mL in 1% DDM overnight at 4 °C. The extraction was centrifuged at 100,000 x *g* for 25 min, and the resulting supernatant was supplemented with 10 mM imidazole prior to purification by immobilized metal affinity chromatography. The extraction containing solubilized MsbA was loaded onto a column packed with 2.5 mL Ni-NTA resin pre-equilibrated in Wash A buffer (20 mM Tris-HCl, 150 mM NaCl, 10 mM imidazole, 10% (v/v) glycerol, pH 8 and supplemented with 2 x the critical micelle concentration (CMC) of DDM). After the loading, the column was washed with 5 column volumes (CV) of Wash B DDM buffer (20 mM Tris, 150 mM NaCl, 25 mM imidazole, 10% (v/v) glycerol, 0.05% DDM, pH 7.0). The immobilized protein was eluted with the addition of 2 CV of Elution buffer (20 mM Tris, 150 mM NaCl, 250 mM imidazole, 10% (v/v) glycerol, 0.05% DDM, pH 7.0). The pooled protein was concentrated using a centrifugal concentrator (Millipore, 100 kDa) prior to injection onto a Superdex 200 Increase 10/300 GL (Cytiva) column equilibrated with 20 mM Tris-HCl (pH 7.0), 150 mM NaCl, 10% (v/v) glycerol and 0.2% DDM. MsbA containing fractions were pooled, concentrated and flash frozen in $LN_2$ and stored at -80°C before use.

#### XylE expression and IIMV preparation

XylE was expressed as previously described[1] and IIMV preparation was carried out as described for MsbA.

#### Protein Quantification in IIMVs

Protein concentration in *E. coli* IMVs containing MsbA or XylE was determined using the Pierce BCA Protein Assay kit (ThermoFisher Scientific) according to manufactures instructions. A standard curve was prepared with BSA of known concentrations. Expression levels of MsbA and XylE in IIMVs was determined by Western blotting. 15, 10 and 5 µg of IIMV was loaded onto a 12% acrylamide gel. A standard curve was prepared with detergent solubilised protein of known concentration. Expression levels were determined from in gel band intensities relative to total loaded protein following transfer to PVDF membranes and using 1:5000 dilution of Anti-His-HRP conjugated antibody (Jackson ImmunoResearch). Intensities were analysed in ImageLab (BioRad). Protein concentration within IIMVs was estimated at 0.48 mg/mL and 2.42 mg/mL for MsbA and XylE respectively.

**Lipid quantification in IIMVs**

The lipid content of isolated membrane samples was determined using the sulpho-phospho-vanillin (SPV) method following manufactures protocols for colorimetric lipid quantification (Cell Bio Labs). Briefly, 15 µL of lipid source (either IIMV or purified lipid standard) was heated to 90°C with 150 µL of 18M sulfuric acid for 10 mins in glass coated 96-well plates. Following cooling on ice for 5 mins, 100 µL of reaction mixture was combined 1:1 with acidic vanillin. The vanillin was left to react for 15 mins at 37°C prior to reading absorbance at OD 540nm. Standards were prepared with suppled purified lipid standards between 300-4.7 mg/dL in DMSO. Standard curves were generated in GraphPad Prism v10.2.2 with non-linear curve fitting using the Pade (1,1) approximant equation. Standard curves were generated in triplicate fit with an $R^2$ of 0.9578. Unknown IIMV concentrations were interpolated to give 0.23 ± 0.8 and 0.75 ± 0.4 mM for MsbA and XylE respectively (n=3). As IIMVs were isolated from *E. coli* the average MW of lipid was estimated at 820 g/mol based on relative levels of phospholipid in total lipid extracts.

**Instrumentation and Automated SEC Delipidation**

To build the DMD platform, experiments were performed using two ACQUITY I-Class UPLC BSMs and a Reaxus MX isocratic pump coupled to a SELECT SERIES cIMS QTOF mass spectrometer (Waters Corporation, Wilmslow, UK). Sample handling, ZrO2 delipidation, and LC valve management were operated by a dual head LEAP HDX automation system with refrigerated HDX manager and lipid filtration system (Trajan, North Carolina, USA). The refrigerated HDX manager contained a three-valve set-up which was reconfigured to allow fully automated on-line SEC separations while column regeneration could be performed simultaneous to the analytical LC gradient.

**Delipidation Valve Set Up**

The original configuration of the HDX manager system consisted of 3 valves (one 6-port, 2 position, sample injection valve; one 10-port, 2-position switching valve; and a 6-port selector valve) which supports back-flushing functionality as previously described[2]. These were connected to two ACQUITY I-Class BSMs pumps named loading and gradient pumps, alongside an isocratic ReaXus MX class pump. To implement the automated dual mode delipidation workflow, these valves were reconfigured, which we now refer to as the sample injection, delipidation, and analytical valves, respectively (**Fig.2**). The developed 8-step valve position cycle for automated SEC delipidation was as follows: In step 1, the sample injection valve is held in the ON position, allowing the robot to load sample into the sample loop post-$ZrO_2$ delipidation (**Fig. S1, step 1**). In step 2, the sample injection valve is switched OFF, allowing the loading pump to flow sample through the SEC column (**Fig. S1, step 2**). Here, the delipidation valve is kept OFF, allowing faster eluting proteins to be directed to the protease and trap columns for digestion and desalting, respectively, while slower eluting lipids are temporarily retained on the SEC column. After a user-specified time window (e.g., 160s), step 3 is triggered by switching the delipidation valve from the OFF to ON position (**Fig. S1, step 3**). Here, loading pump continues providing flow through the SEC column to waste, allowing later eluting lipids to be discarded, while the isocratic pump provides flow through the protease and trap columns to continue digestion/desalting. After trapping, step 4 is triggered by switching the analytical valve from position 2 to 3, allowing the gradient pump to elute trapped peptides to separate them in the analytical column (**Fig. S1, step 4**). Here, the delipidation valve is kept in the ON position, so that the SEC column can be washed with 25% methanol using the loading pump (**Fig. S1, step 5**). After three minutes (**Fig. S1, step 6**), the

loading pump goes back to 100% 0.1% FA in $H_2O$, allowing the robot to inject the SEC wash into the sample loop. The sample valve position is alternated between OFF and ON positions to direct SEC wash through the SEC column. Once 240 sec has elapsed following the final SEC wash injection (giving time for all SEC wash to be flushed to waste and preventing damage to the protease), step 5 is triggered by switching the delipidation valve to the OFF position so that pepsin wash can be injected into the sample loop and sample valve position alternated between OFF and ON to direct pepsin wash through both the SEC and protease columns (**Fig. S1, step 7**). Steps 4 - 7 are carried out during the analytical gradient. By doing so, both SEC and protease column washing are achieved simultaneously to the analytical gradient, thereby maintaining analytical throughput relative to experiments using only $ZrO_2$-mediated delipidation. After the analytical gradient has elapsed, the analytical valve is switched back to position 2 allow the system to equilibrate for the next sample injection (**Fig. S1, step 8**).
Similar valve configuration was used for the blank runs where the SEC column is washed with 20-25% 50:50 IPA:$H_2O$ containing 0.1% fos-choline-12 using the solvent selection valve of the I-Class loading pump.

**$ZrO_2$ beads pre-treatment**

To reduce non-specific interactions[3], 300mg of $ZrO_2$ coated silica beads (Sigma Aldrich) were incubated overnight with a 3% Bovine Serum Albumin (BSA) solution at 4°C. Beads were later washed five times with 0.1% formic acid in water and diluted to a final concentration of 300 mg/mL treated beads (w$ZrO_2$) were immediately used for delipidation purposes.

**POPC stock preparation**

1-palmitoyl-2-oleoyl-glycero-3-phosphocholine (POPC) (Avanti Polar Lipids) was prepared from 25mg/mL chloroform stocks. Chloroform was removed by a continuous flow of $N_2$ to generate a lipid film. POPC films were hydrated to 4 mg/mL in 100mM KPi (pH7) through sonication and 3x freeze-thaw cycles. Formed multilamellar vesicles were flash frozen with liquid $N_2$ and stored at -20°C until use. Upon use, aliquots were slowly thawed on ice and vortexed to ensure resuspension of lipid prior to further dilution to working concentration.

**DMD characterisation**

Delipidation efficiency was evaluated using POPC as a model system. The Trajan HDX manager was equipped with an Agilent ZORBAX 300SB-C3 (2.1 x 12.5mm, 5 µm) guard column. POPC stock solutions were further diluted to 200 µL in 0.1% formic acid in water and injected into a 250 µL sample loop. Samples were then trapped and desalted for 3 min at 200 µL/min using mobile phase A (0.1% formic acid in $H_2O$), followed by an 8 to 85% gradient of mobile phase B (0.1% formic acid in ACN) over 3 mins at room temperature. A 9 min washing step held at 85% B was then subsequently used to prevent carryover. Calibration curve (n=3) (1-50pmol) of POPC (**Fig. S2**) resulted in a correlation coefficient $R^2$=0.9974 and served as basis for quantification.

For $ZrO_2$-mediated delipidation, 10 µL of pre-treated $ZrO_2$ beads in 0.1% formic acid in water were pre-dispensed into the outer base of chilled nanofilter vials (Thomson Instrument Company). POPC stocks were then diluted up to 200 µL in 0.1% formic acid in water, followed by mixing with w$ZrO_2$ in the nanofilter base and filtering (Thomson Instrument Company), as previously described[4]. 195 µL of filtrate was then injected into the 250 µL sample loop for LC-MS analysis, resulting in "nominal" POPC loads of 10, 100, and 1000 pmol. For SEC-mediated delipidation, the above LC-MS method was performed but with a Waters BEH SEC guard column (125 Å, 1.7 µm, 4.6 x 30 mm, 1–80 K) installed between the sample injection and delipidation valves as described in the delipidation valve set up. Similarly, POPC stocks were diluted up to 200 µL in 0.1% formic acid in water, followed by injection of 195 µL to provide effective POPC loads of 10, 100, 1000 and 2500 pmol, respectively. For DMD experiments, a similar workflow was employed as describe above. POPC stocks were diluted to 200 µL in 0.1% formic acid in water, mixed with 10 µL of w$ZrO_2$ and filtered. 195 µL of the filtrate were injected for subsequent SEC-mediated delipidation, achieving "nominal" POPC loads of 10, 100, 1000,

2500 and 3500 pmol. All experiments were performed in triplicate (n=3) and used the same LC-MS workflow. Residual POPC and delipidation efficiency was determined using the initial POPC calibration curve (**Table S1**).

For protein recovery experiments, bovine carbonic anhydrase (BCA) was used as a model system. BCA stock solution was diluted with 95 µL equilibration buffer (20mM Tris, 150mM NaCl, pH:7.0) and quenched with 100 µL pre-cooled quench buffer (100mM Glycine, 4M Urea, 0.2% DDM, pH 2.3) mimicking an HDX experiment. Samples resulting in "nominal" amount of 60 pmol were injected into the MS with no delipidation, with standalone $wZrO_2$, with SEC delipidation, or using the DMD workflow. Protein recovery was calculated by comparing peak areas to the non-delipidated sample (**Fig. S3**). All intact mass experiments were performed in triplicate (n=3).

All extracted ion chromatograms (EIC) were generated on MassLynx (v. 4.2) by using the full width half height (FWHM) of the monoisotopic peak of monomeric POPC ion, and the FWHM of the intact charge state distribution for BCA. EIC peak areas were then integrated on MassLynx with a peak-to-peak amplitude of 2000.

**Deuterium/hydrogen back-exchange evaluation**

To evaluate system back exchange, 20 pmol Neurotensin 8-13 (RRPYIL), 20 pmol Angiotensin II (DRVYIHPF), and 20 pmol Substance P (RPKPQQFFGLM-NH2) were diluted up to 100 µL with labelling buffer (20 mM HEPES, pD 7.0) and incubated for 30 mins at room temperature to achieve maximal deuteration. The HDX reaction was then quenched by 1:1 dilution with quench buffer (100 mM $K_2HPO_4$, pH 2.5). LC-MS experiments were then run with the standard method using no delipidation (ST), SEC delipidation, $ZrO_2$ delipidation and with DMD, all in triplicate. Back-exchange was manually calculated using the centroid mass for each isotopic distribution as described elsewhere[5].

### Supplementary Figure and Tables

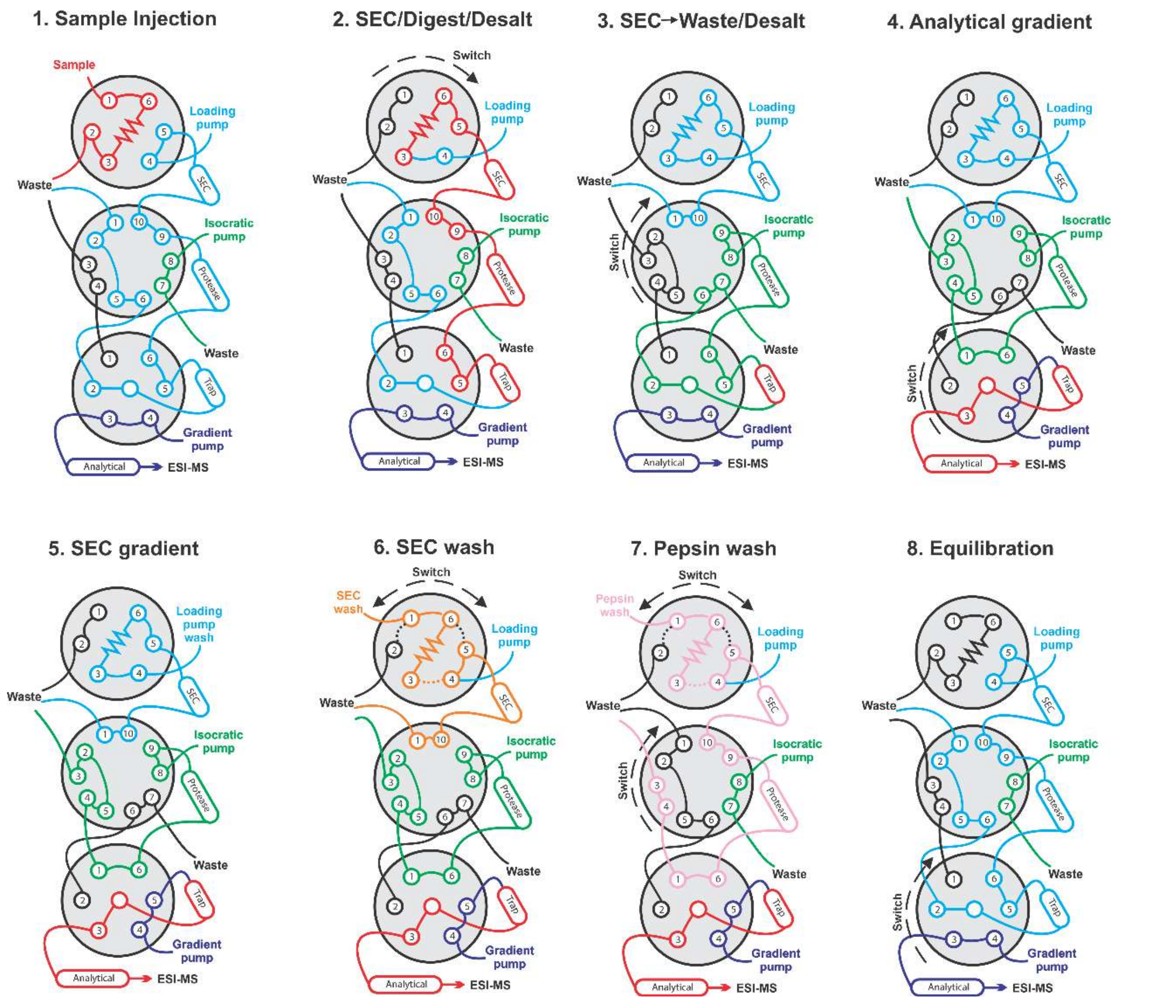

**Figure S1 – Overview of valve reconfiguration on Trajan HDX management system for automated on-line SEC-delipidation and cleaning**

Each experiment consists of an 8-step cycle involving valve position changes and mobile phase gradients. Red indicates the protein/peptide location at each step, blue indicates the loading pump flow path, green indicates the isocratic pump flow path, purple indicates the gradient pump flow path, and black indicates no flow. See methods for full details.

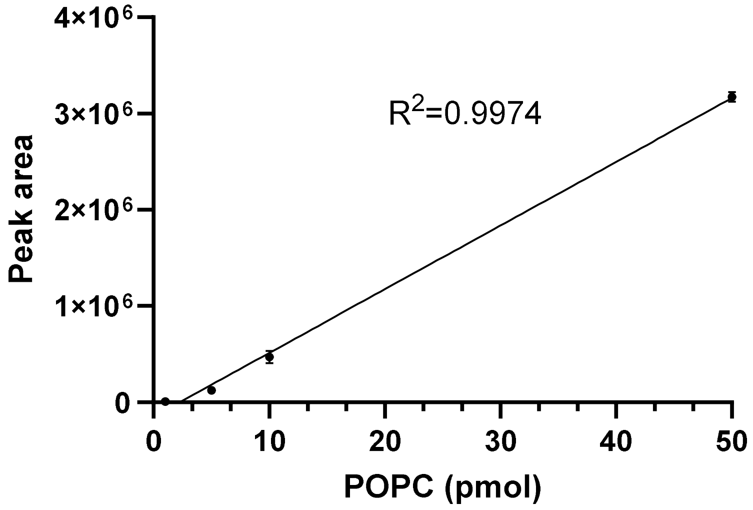

**Figure S2 – POPC calibration curve**
Detector calibration curve of POPC (1-50 pmol) with a correlation coefficient $R^2$=0.9974. EICs were generated using monoisotopic POPC ion and integrated to obtain peak area. Error bars represent the standard deviation of the measurements (n=3).

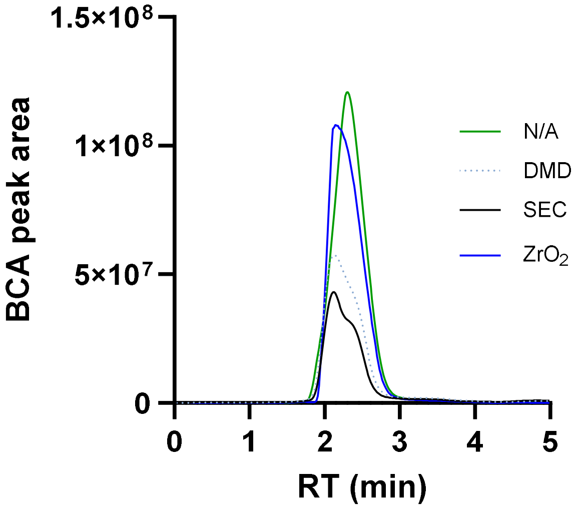

**Figure S3 – BCA protein recovery**
Protein recovery was calculated using BCA as a model system. Representative EIC of 60 pmol BCA following the standard method (N/A) without delipidation (green), SEC delipidation (black), $ZrO_2$ delipidation (blue), and using DMD (dotted blue). All experiments were performed in triplicate. Protein recovery resulted in ~107.6% for $ZrO_2$, ~61.2% for SEC and ~48.4% for the DMD workflow compared to the standard method without delipidation (ST) workflow.

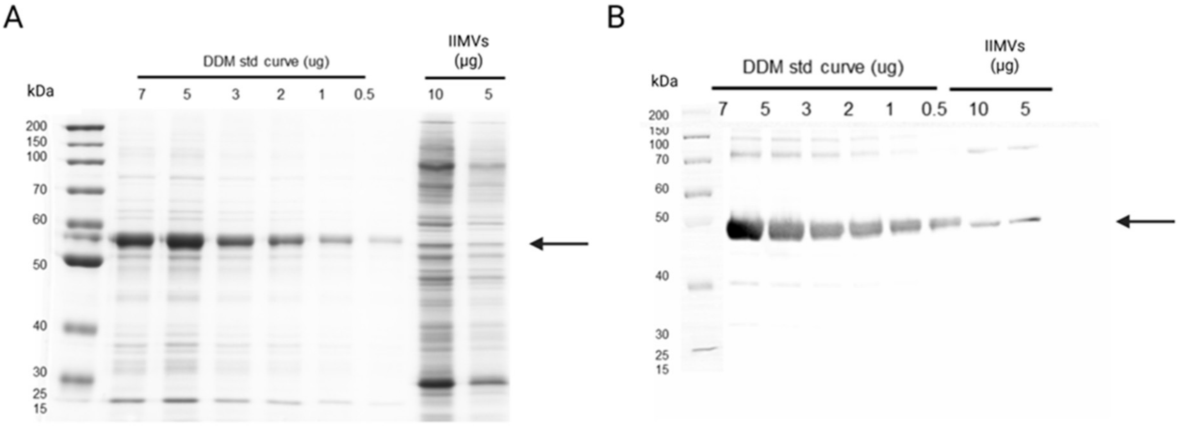

**Figure S4 – MsbA expression in *E. coli* IIMVs.**
(A) A representative Coomassie stained SDS-PAGE of MsbA (indicated with an arrow) containing IIMVs against a standard curve of MsbA in DDM. Intensities were calculated in ImageLab (BioRad) to quantify expression levels. (B) Confirmation of MsbA (indicated with an arrow) at the relative front taken from IIMVs in Coomassie gels was performed through western blot using 1:5000 dilution of Anti-His-HRP conjugated antibody (Jackson ImmunoResearch).

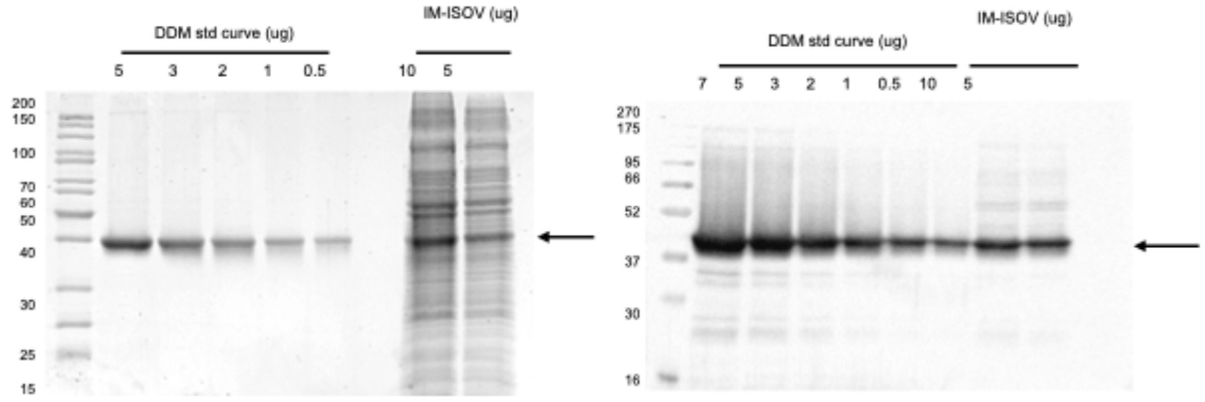

**Figure S5 – XylE expression in *E. coli* IIMVs.**
(A) A representative Coomassie stained SDS-PAGE of XylE (indicated with an arrow) containing IIMVs against a standard curve of XylE in DDM. Intensities were calculated in ImageLab (BioRad) to quantify expression levels. (B) Confirmation of XylE (indicated with an arrow) at the relative front taken from IIMVs in Coomassie gels was performed through western blot using 1:5000 dilution of Anti-His-HRP conjugated antibody (Jackson ImmunoResearch).

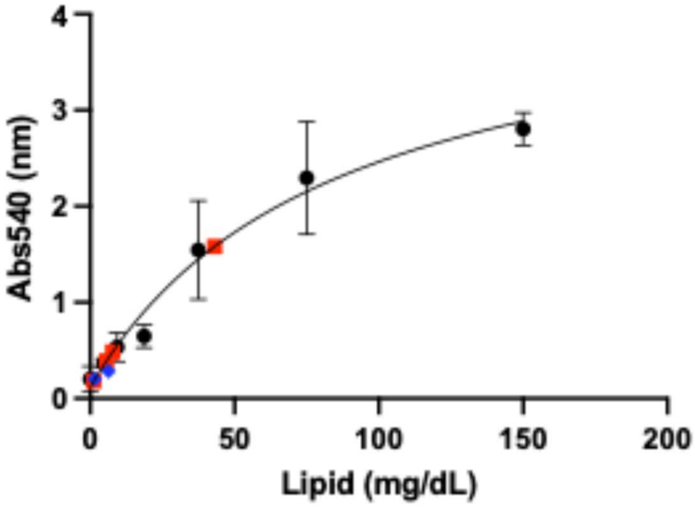

**Figure S6 – Lipid quantitation in *E. coli* IIMVs.**

Lipid quantification was carried out using SPV quantification (Cell Bio Labs). A standard curve was generated with the lipid standard (black) and the MsbA IIMVs (red) were diluted 1:1, 1:5, 1:10 and 1:20 in DMSO for quantification while the XylE IIMVs (blue) were diluted 1:50 and 1:100. Standard curves were generated in GraphPad Prism v10.2.2 with non-linear curve fitting using the Pade (1,1) approximant equation. Non-linear standard curve fitted is anticipated for SPV quantification. Standard curves were generated in triplicate fit with an $R^2$ of 0.9827. All measurements were performed in triplicate.

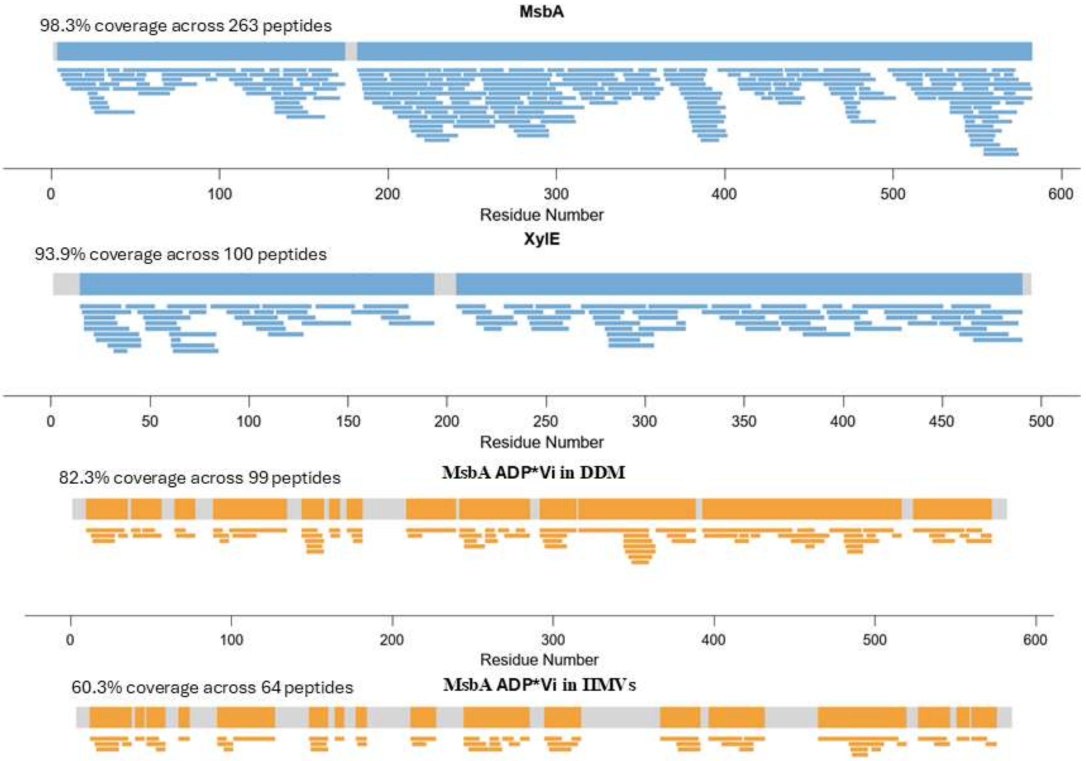

**Figure S7 – MsbA and XylE peptide identifications and effective peptide coverage maps**

In blue, positive peptide identifications for MsbA and XylE using IIMVs samples following a DMD workflow. 263 positive peptides identifications representing 98.3% sequence coverage and 100 positive identifications for a 93.9% sequence coverage were obtained for MsbA and XylE IIMVs respectively. All experiments were performed in triplicate (n=3) In orange, effective HDX peptide coverage map after manual curation. 99 peptides resulting in 82.3% sequence coverage for MsbA ADP*Vi in DDM and 64 peptides and 60.3% sequence coverage for MsbA ADP*Vi in IIMVs.

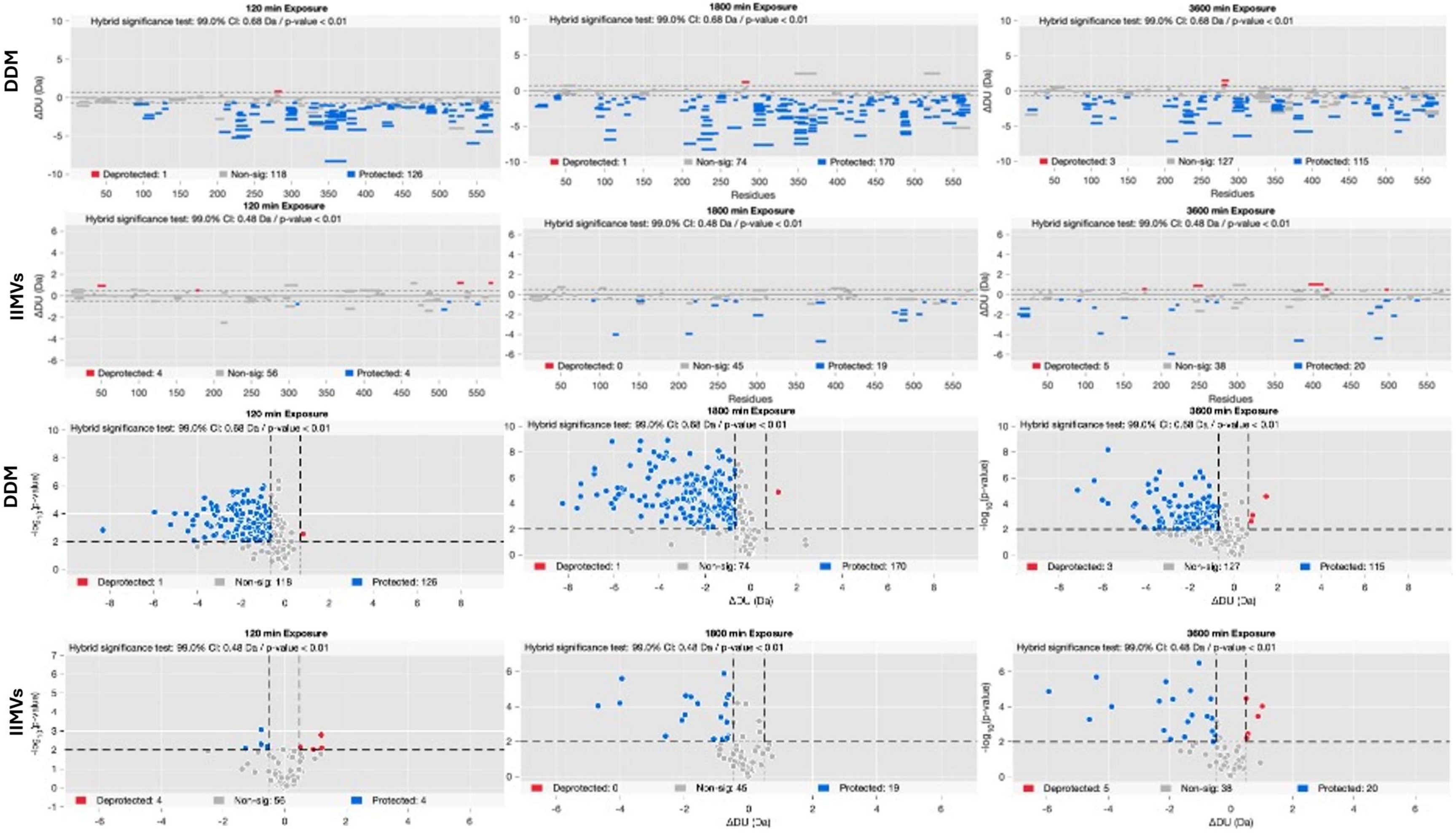

**Figure S8 – Woods and Volcano plots obtained from differential HDX-MS in DDM and IIMV environments.**

Top panels represent woods plots of statistically protected or deprotected peptides and non-significant peptides along the length of the protein sequence. Each bar represents a single peptide with peptide length indicated by the bar length. Bottom panels represent Volcano plot representation of the same data where each dot is equivalent to a peptide. All data represents hybrid statistical testing. Lower deuterium uptake (protected) peptides are shown in blue and increased deuterium uptake (deprotected) are shown in red.

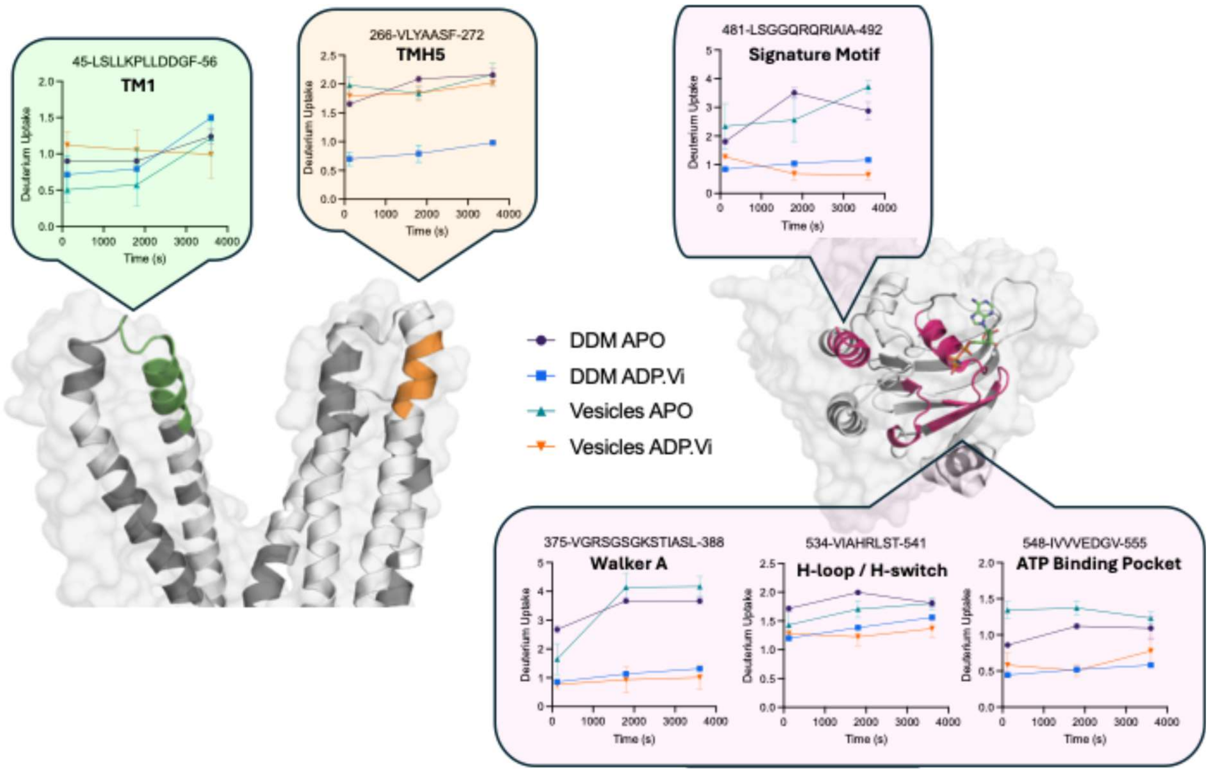

**Figure S9 – E. coli membranes alter the dynamics of MsbA Transmembrane Domain.** Comparison of deuteration uptake between DDM and Vesicle MsbA for selected peptides with similar deuteration levels in the apo state (non-significant differences in uptake). Relative HDX differences between 1min and 60 min between the DDM purified and vesicle states in the ADP*Vi trapped state (p-value < 0.01) are mapped onto the MsbA cryo-EM structure in the OF state. Regions more protected in ADP*Vi bound vs APO are shown in orange and pink, and regions less protected are shown in green, while other protein regions are while. Coverage is shown in white representing peptide identification from ADP*Vi bound vesicles. No coverage is shown in grey.

| | Delipidation workflow | | | | | |
|---|---|---|---|---|---|---|
| | $ZrO_2$ | | SEC | | DMD | |
| initial POPC amount (pmol) | post delipidation amount (pmol) | delipidation efficiency | post delipidation amount (pmol) | delipidation efficiency | post delipidation amount (pmol) | delipidation efficiency |
| 10 | 8.2 | 17.6 | 1.7 | 82.9 | 2.5 | 74.7 |
| 100 | 27.0 | 73.0 | 1.6 | 98.4 | 10.6 | 89.4 |
| 1000 | >50 | - | 3.2 | 99.7 | 4.1 | 99.6 |
| 2500 | - | - | 40.8 | 98.4 | 12.1 | 99.5 |
| 3500 | - | - | - | - | 25.4 | 99.3 |

n=3

**Table S1 – Delipidation workflows efficiency.**
Delipidation efficiency was calculated using POPC as a model system. Post delipidation amount of POPC was calculated using a POPC calibration curve. All experiments were performed in triplicate (n=3).
