## Supplementary material for "An Automated HDX-MS Platform for *in situ* characterisation of Membrane Proteins": Uptake Plots - HDX Data

MSBA 10-27: WQTFRRLWPTIAPFKAGL (#1)

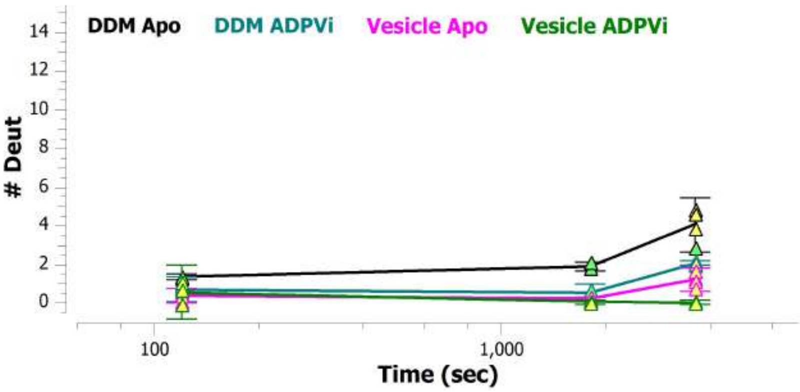


MSBA 101-108: RRRLFGHM (#18)

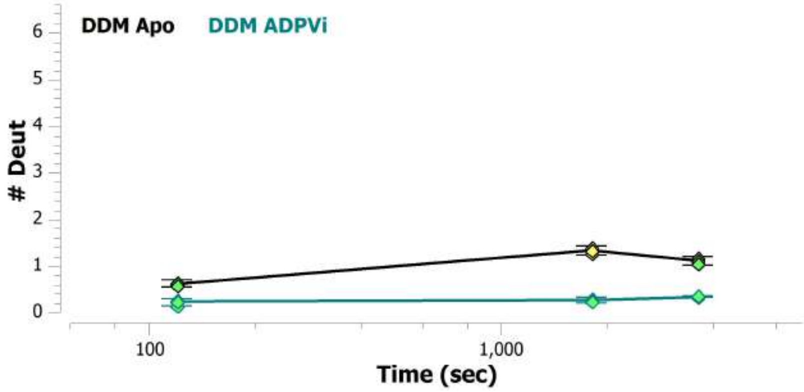


MSBA 101-115: RRRLFGHMMGMPVSF (#18)

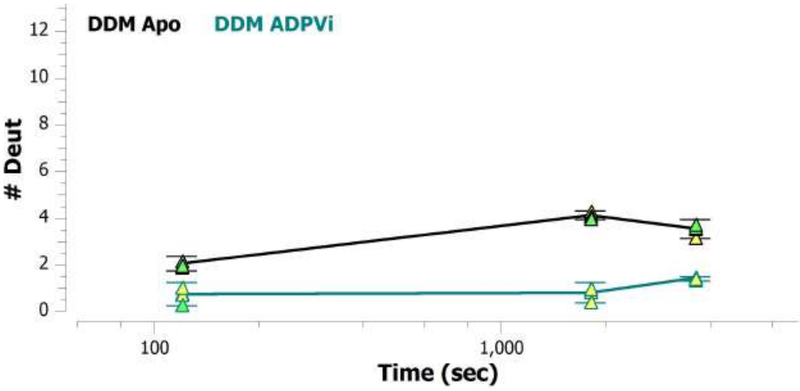


MSBA 109-115: MGMPVSF (#19)

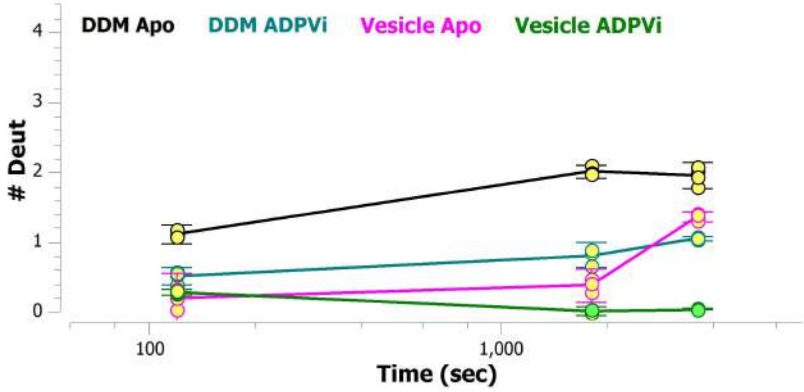


MSBA 116-124: FDKQSTGTL (#20)

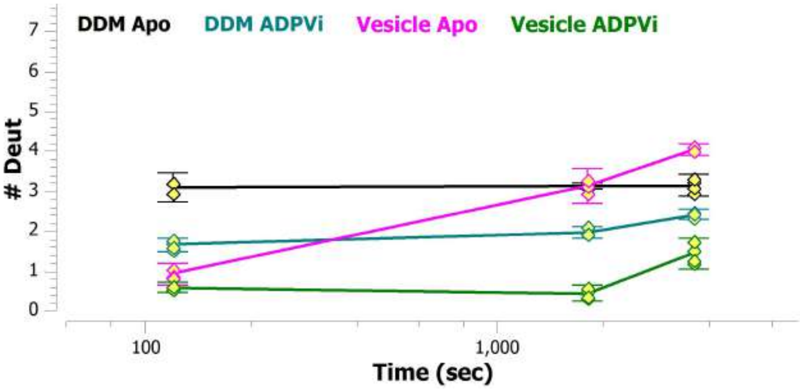


MSBA 116-125: FDKQSTGTLL (#19)

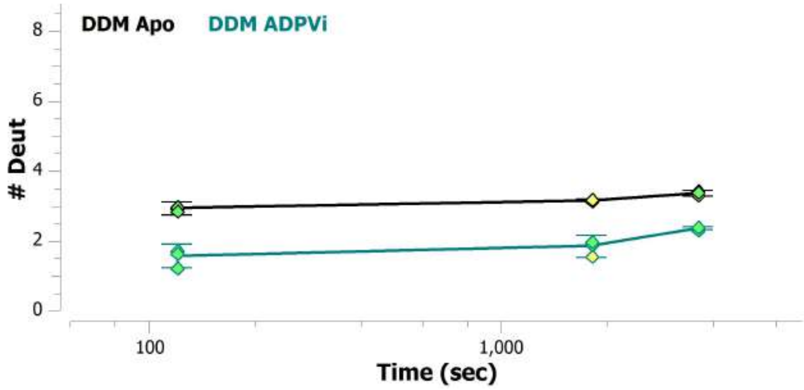

MSBA 125-130: LSRITY (#20)

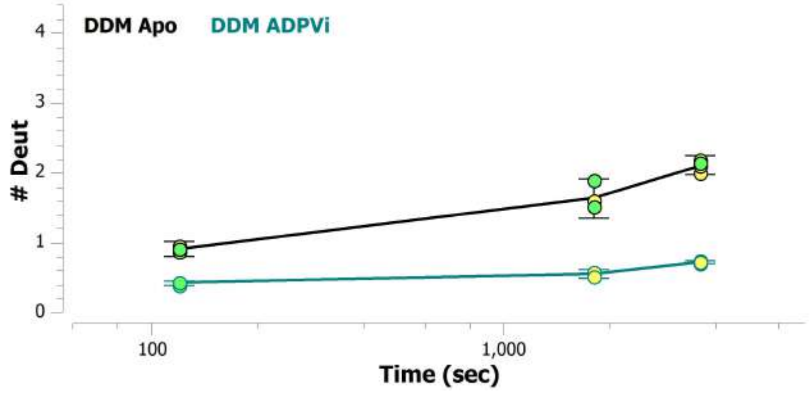


MSBA 125-134: LSRITYDSEQ (#21)

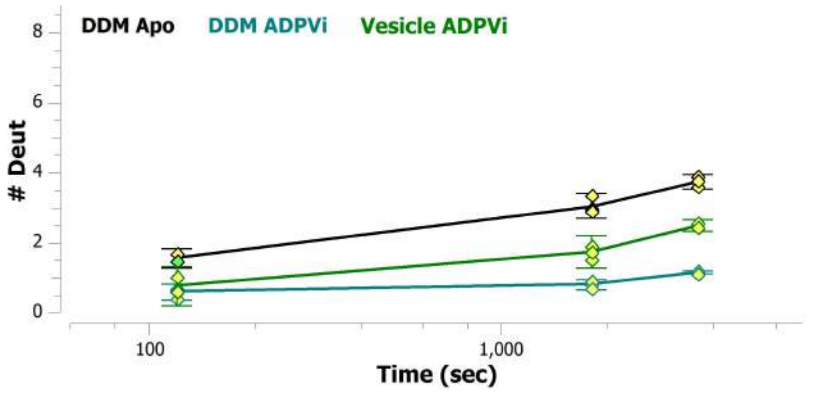


MSBA 126-133: SRITYDSE (#21)

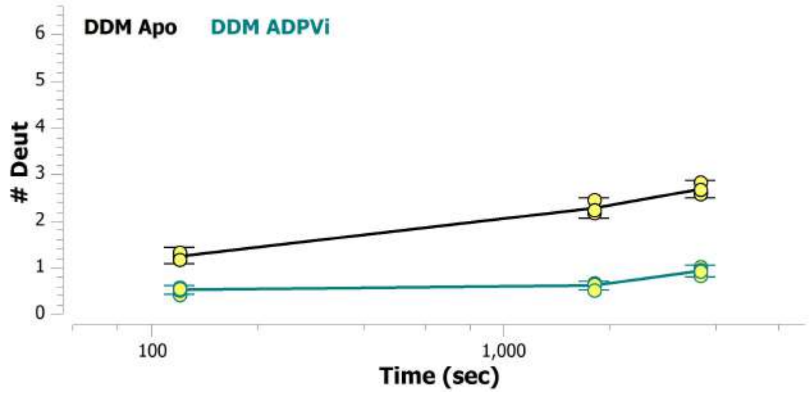


MSBA 13-27: FRRLWPTIAPFKAGL (#2)

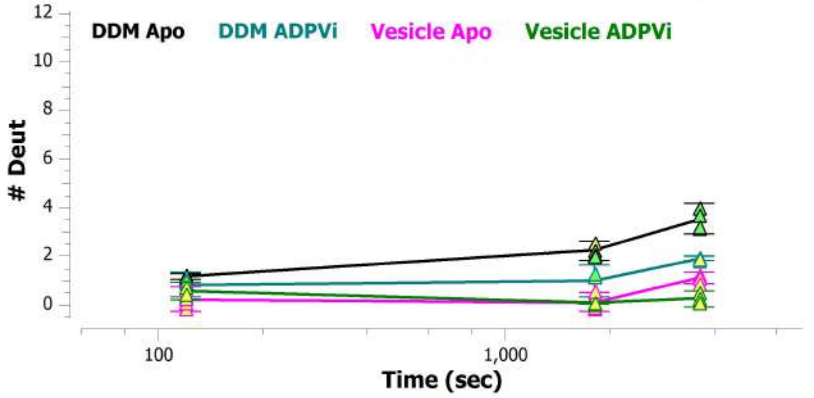


MSBA 134-143: QVASSSSGAL (#22)

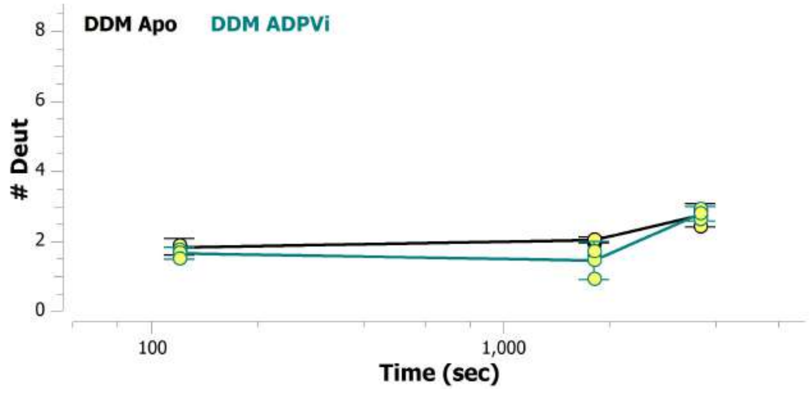


MSBA 135-143: VASSSGAL (#23)

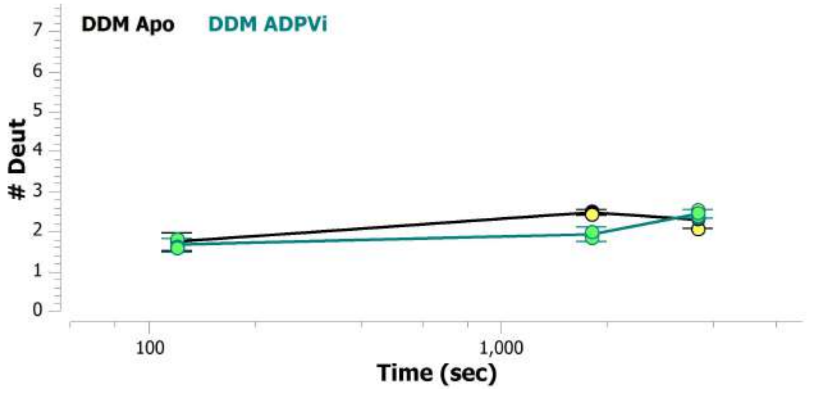

MSBA 135-145: VASSSSGALIT (#24)

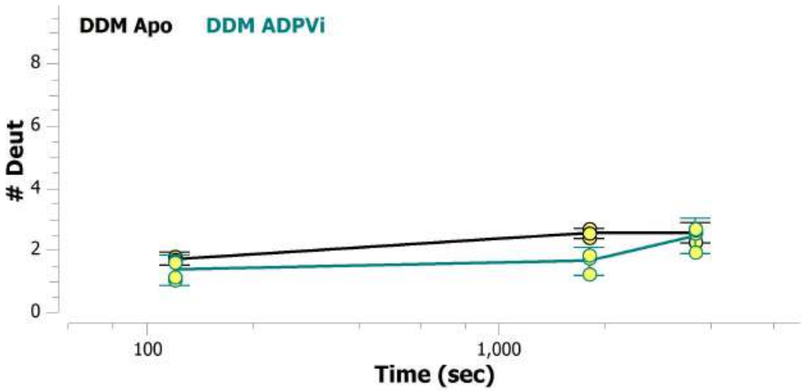


MSBA 14-27: RRLWPTIAPFKAGL (#3)

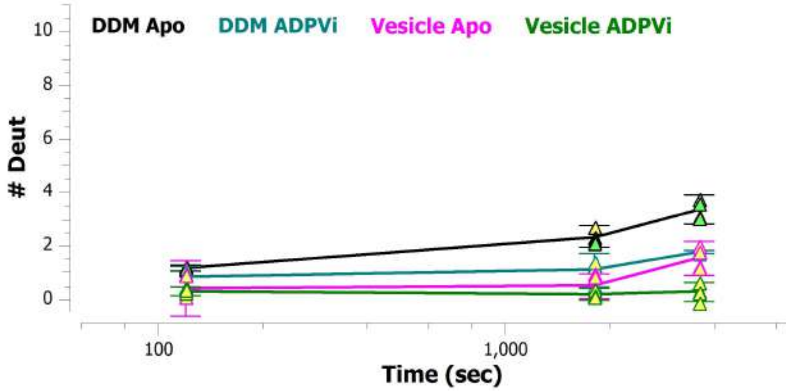


MSBA 144-153: ITVVREGASI (#22)

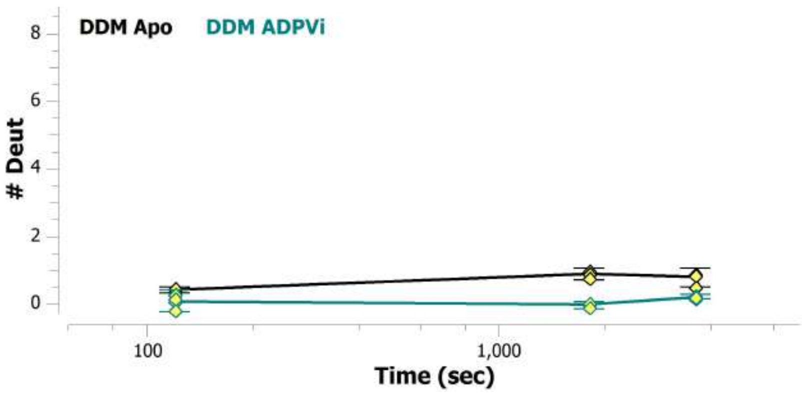


MSBA 144-156: ITVVREGASIIGL (#23)

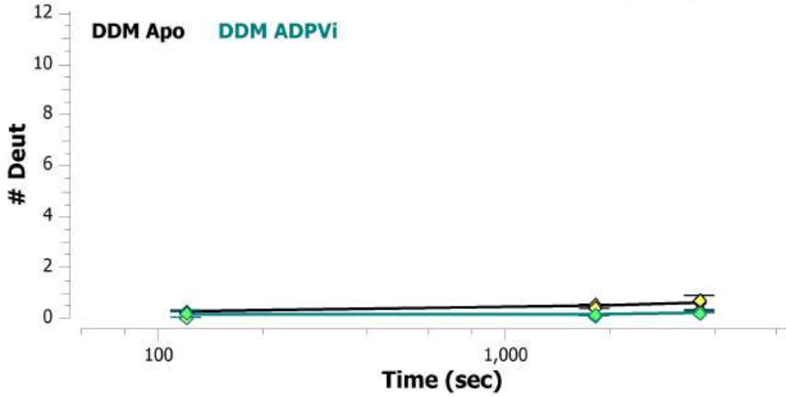


MSBA 145-158: TVVREGASIIGLFI (#25)

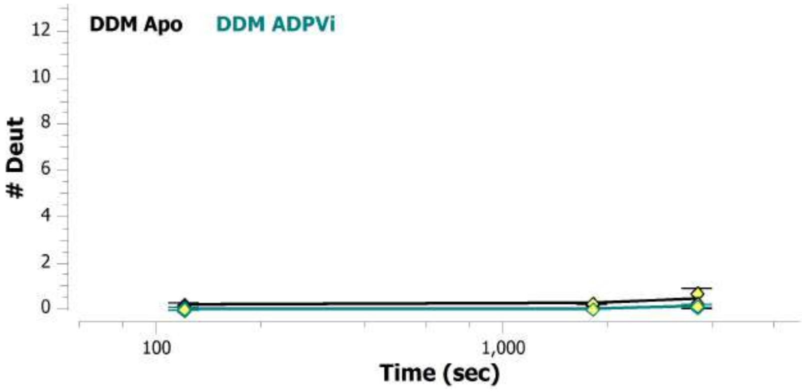


MSBA 146-156: VVREGASIIGL (#26)

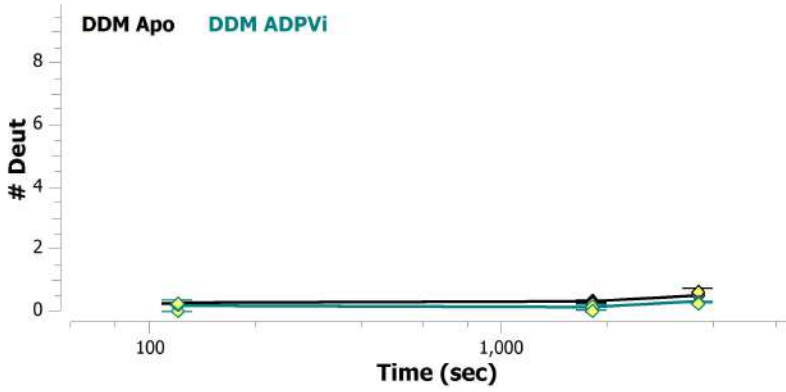

MSBA 146-157: VVREGASIIGLF (#24)

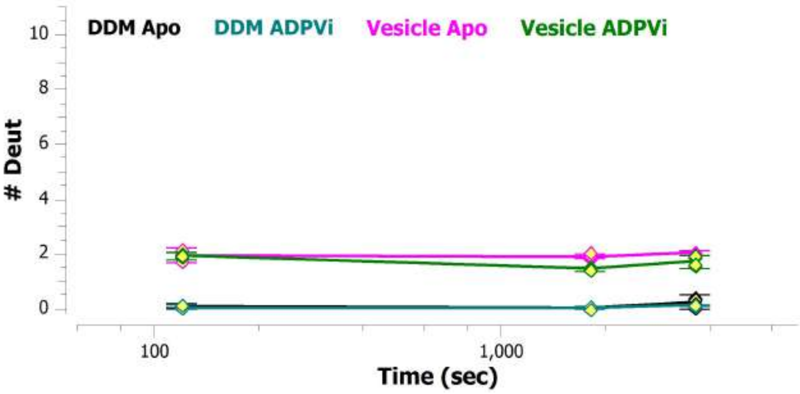


MSBA 147-156: VREGASIIGL (#25)

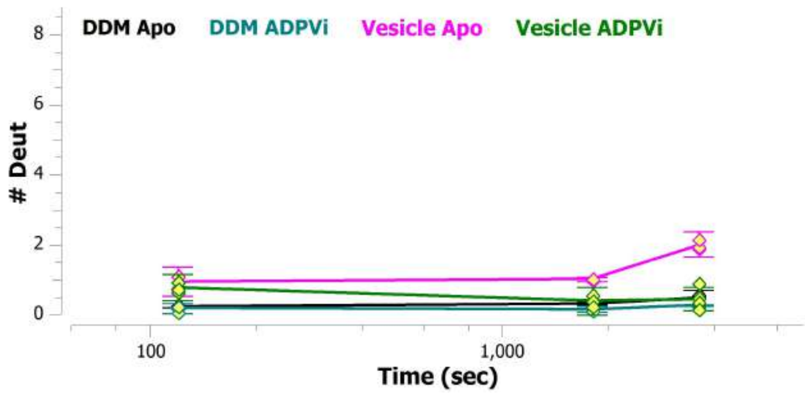


MSBA 147-157: VREGASIIGLF (#26)

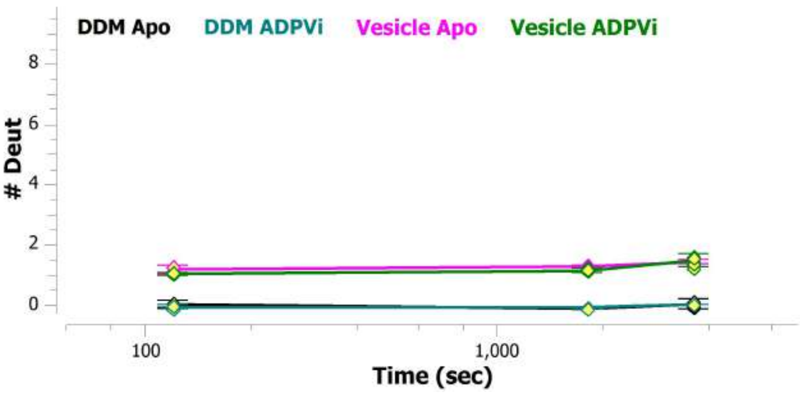


MSBA 161-166: FYYSWQ (#27)

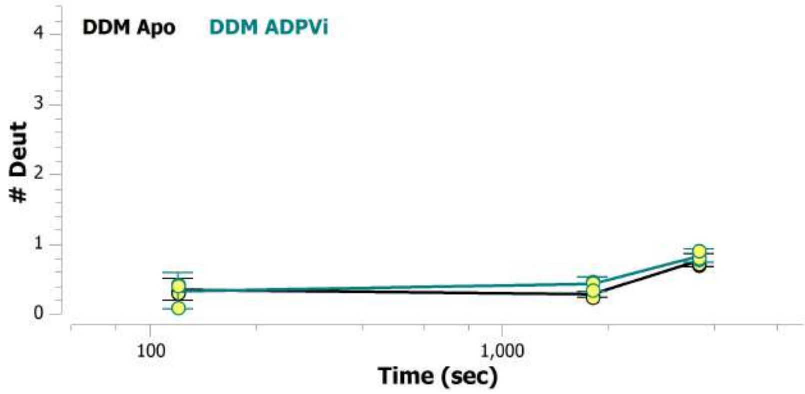


MSBA 161-167: FYYSWQL (#27)

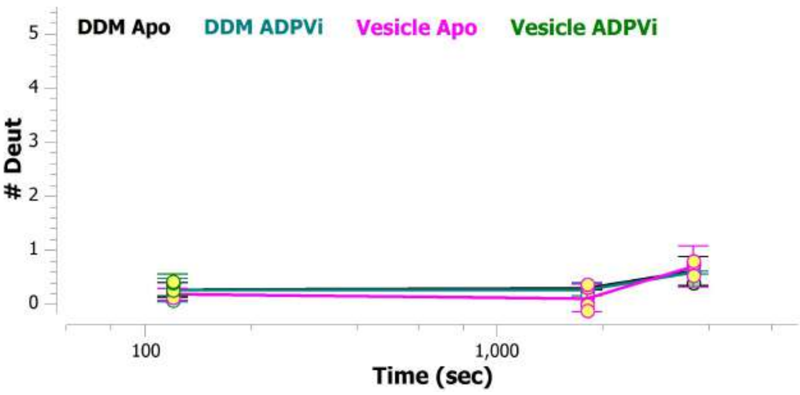


MSBA 162-167: YYSWQL (#28)

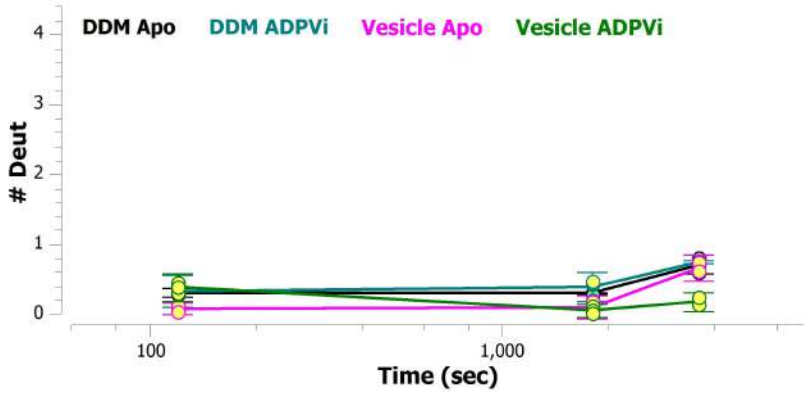

MSBA 172-179: IVLAPIVS (#28)

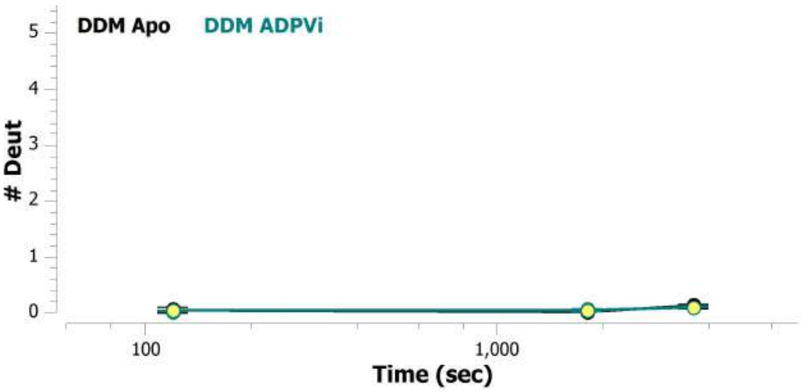


MSBA 172-181: IVLAPIVSIA (#29)

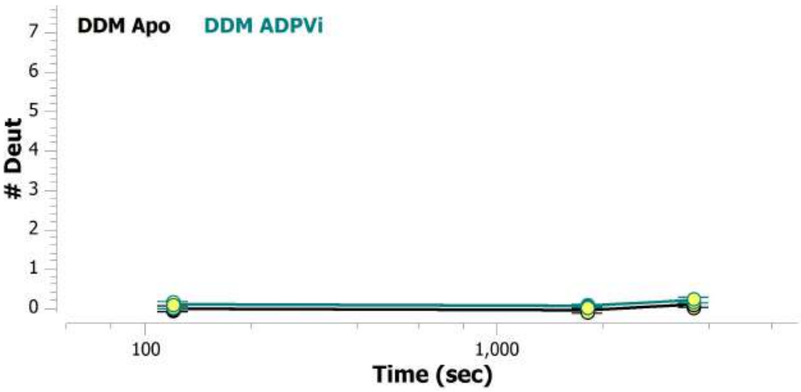


MSBA 175-181: APIVSIA (#30)

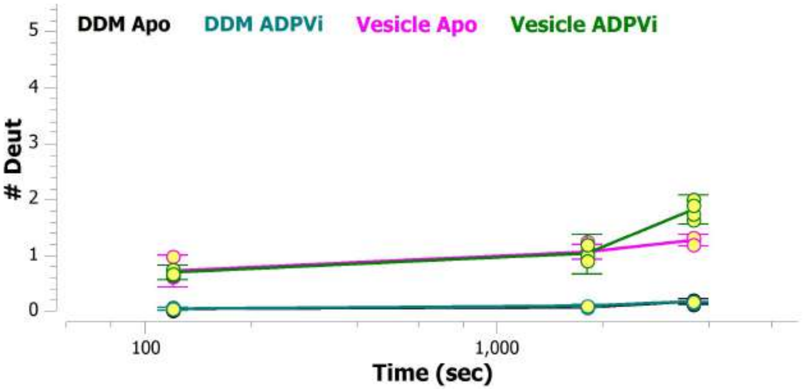


MSBA 176-181: PIVSIA (#31)

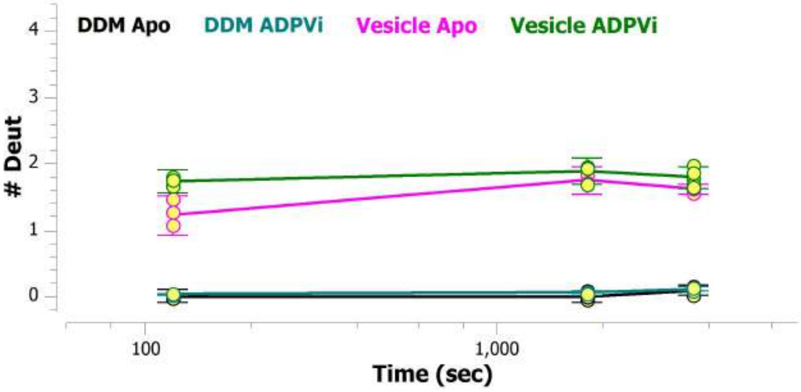


MSBA 197-209: QNTMGQVTTSAEQ (#29)

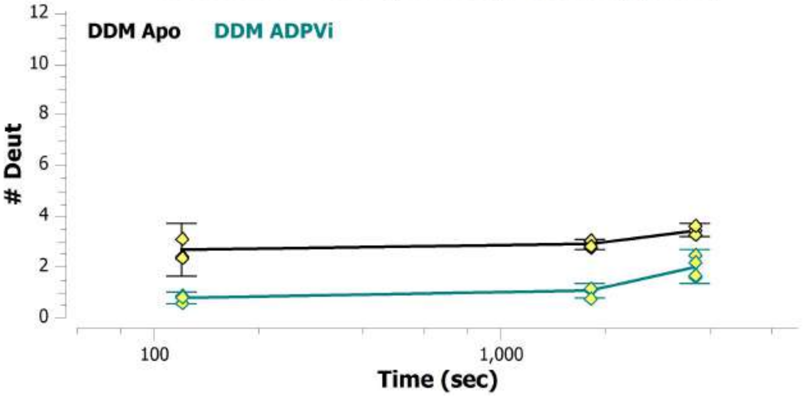


MSBA 202-208: QVTTSAE (#30)

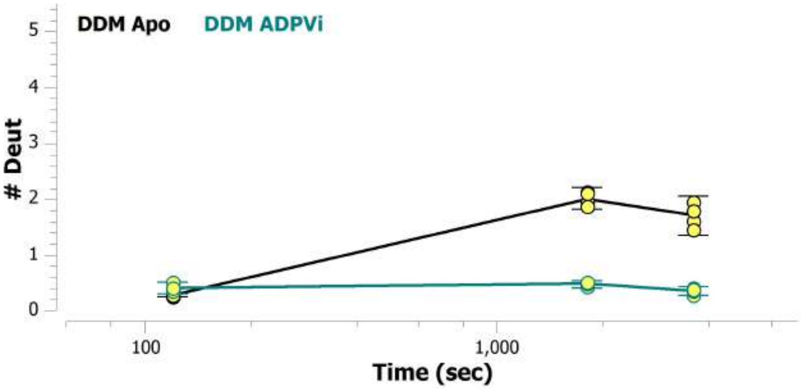

MSBA 202-218: QVTTSAEQMLKGHKEVL (#31)

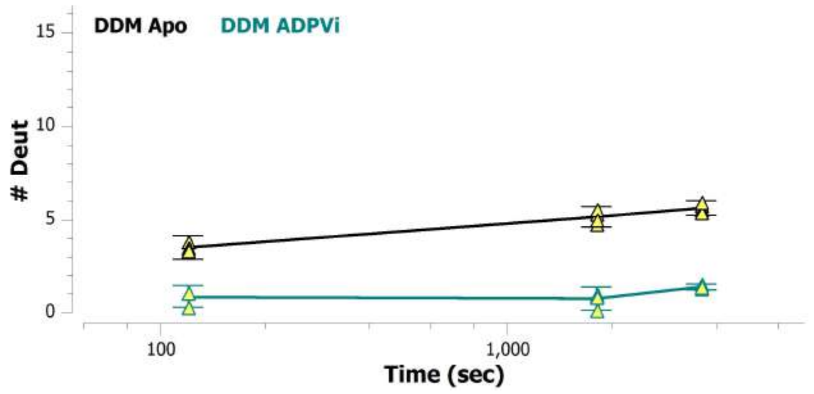


MSBA 203-208: VTTSAE (#32)

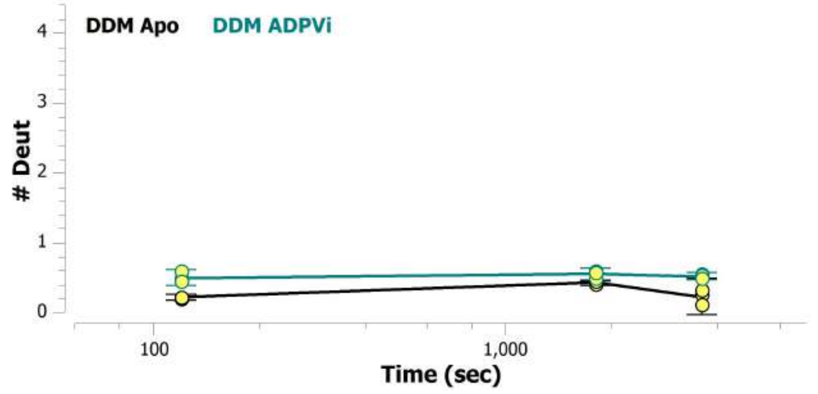


MSBA 208-218: EQMLKGHKEVL (#33)

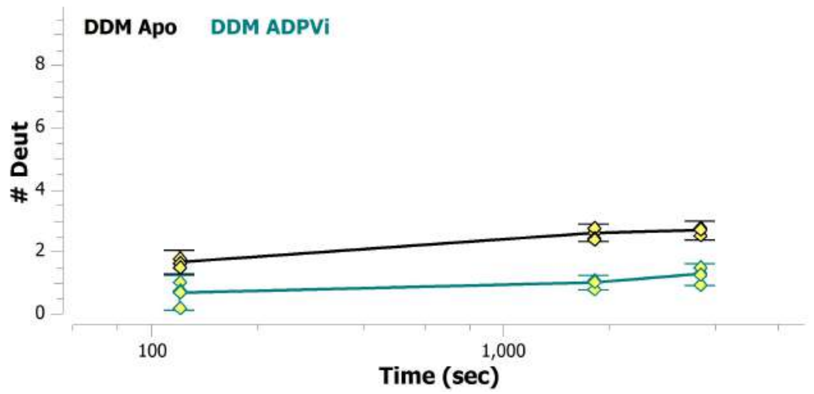


MSBA 209-218: QMLKGHKEVL (#32)

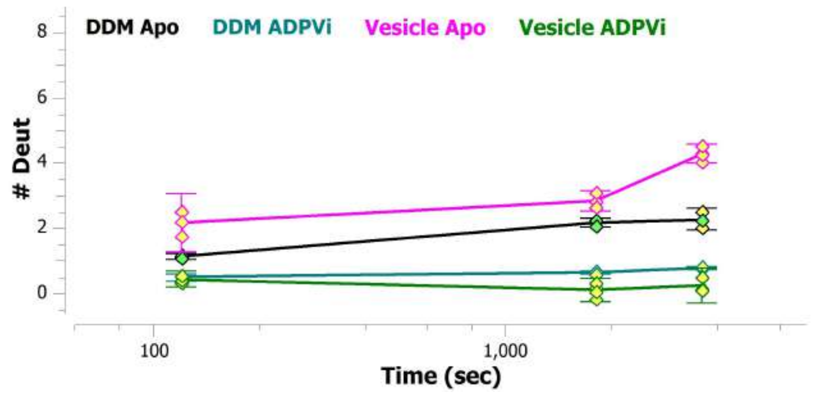


MSBA 210-218: MLKGHKEVL (#33)

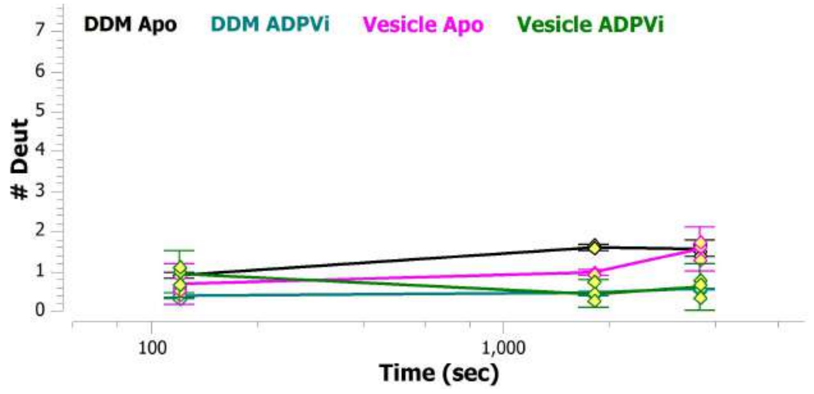


MSBA 211-218: LKGHKEVL (#34)

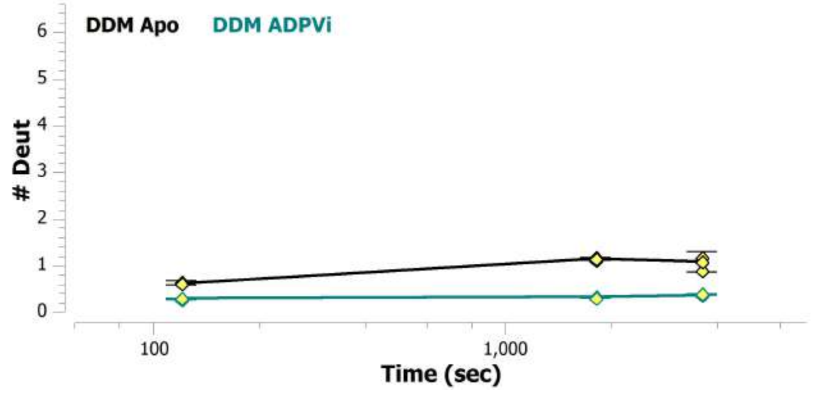

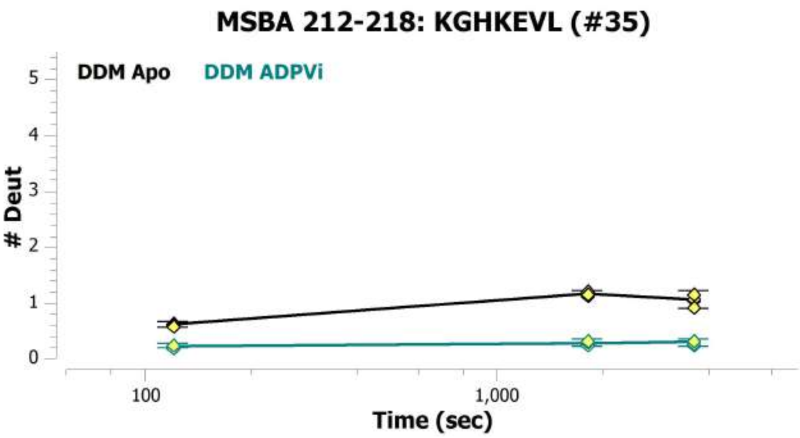
MSBA 212-218: KGHKEVL (#35)
DDM Apo
DDM ADPVi
### Deut
0
1
2
3
4
5
100
1,000
Time (sec)

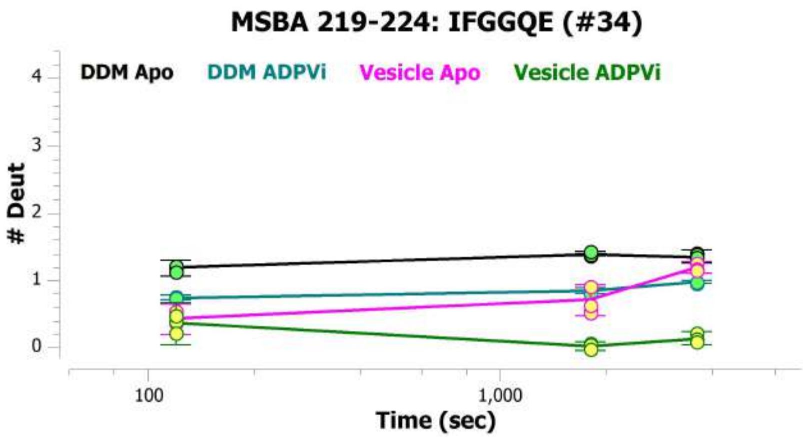
MSBA 219-224: IFGGQE (#34)
DDM Apo
DDM ADPVi
Vesicle Apo
Vesicle ADPVi
### Deut
0
1
2
3
4
100
1,000
Time (sec)

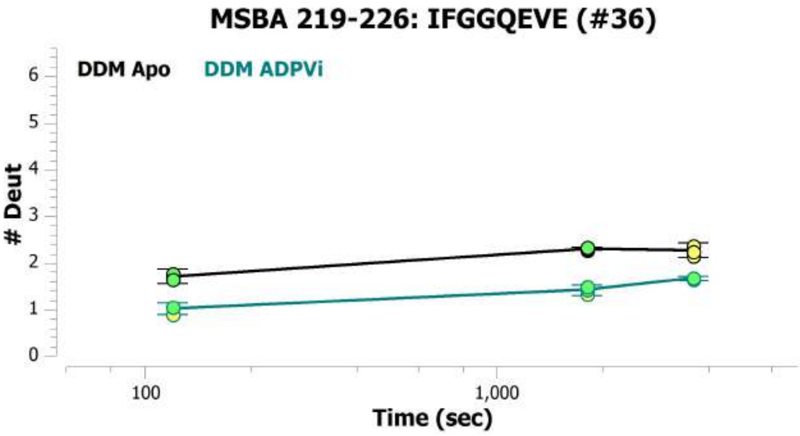
MSBA 219-226: IFGGQEVE (#36)
DDM Apo
DDM ADPVi
### Deut
0
1
2
3
4
5
6
100
1,000
Time (sec)

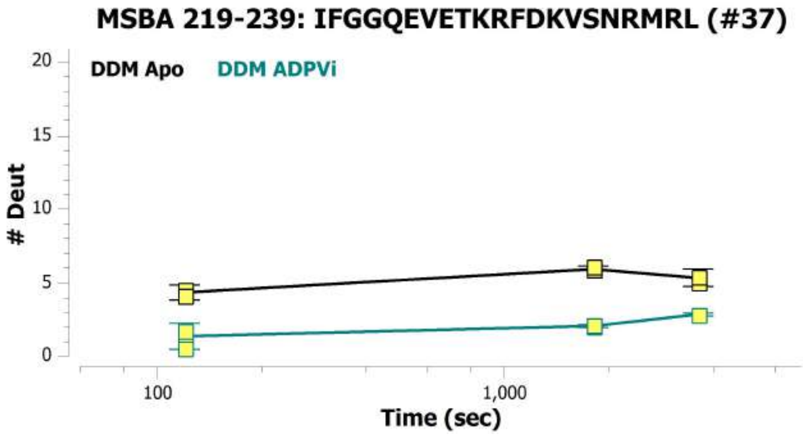
MSBA 219-239: IFGGQEVETKRFDKVSNRMRL (#37)
DDM Apo
DDM ADPVi
### Deut
0
5
10
15
20
100
1,000
Time (sec)

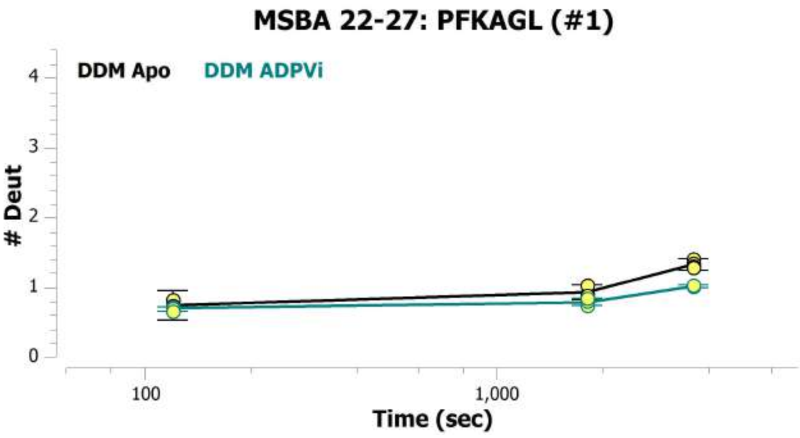
MSBA 22-27: PFKAGL (#1)
DDM Apo
DDM ADPVi
### Deut
0
1
2
3
4
100
1,000
Time (sec)

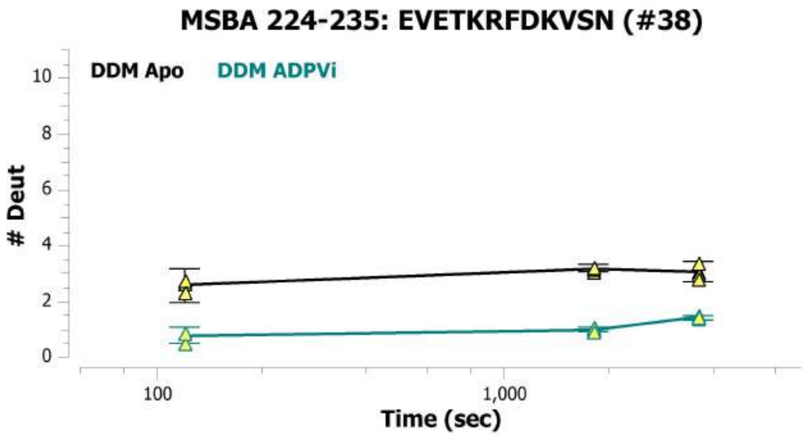
MSBA 224-235: EVETKRFDKVSN (#38)
DDM Apo
DDM ADPVi
### Deut
0
2
4
6
8
10
100
1,000
Time (sec)

##### MSBA 224-247: EVETKRFDKVSNRMRLQGMKMVSA (#39)

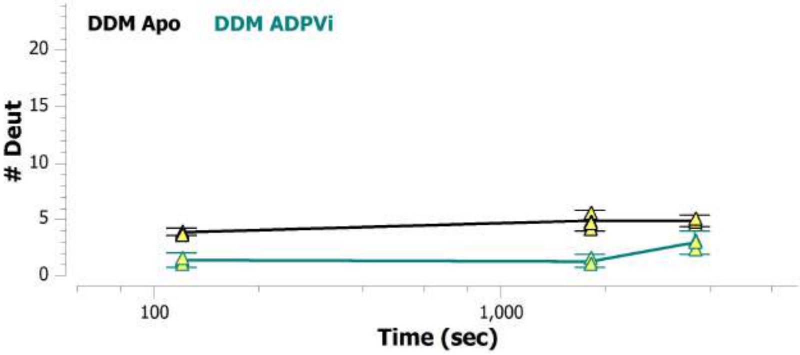


##### MSBA 225-235: VETKRFDKVSN (#40)

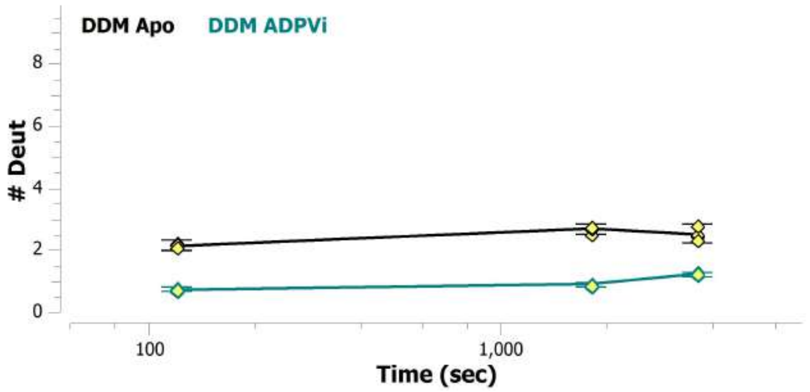


##### MSBA 225-239: VETKRFDKVSNRMRL (#35)

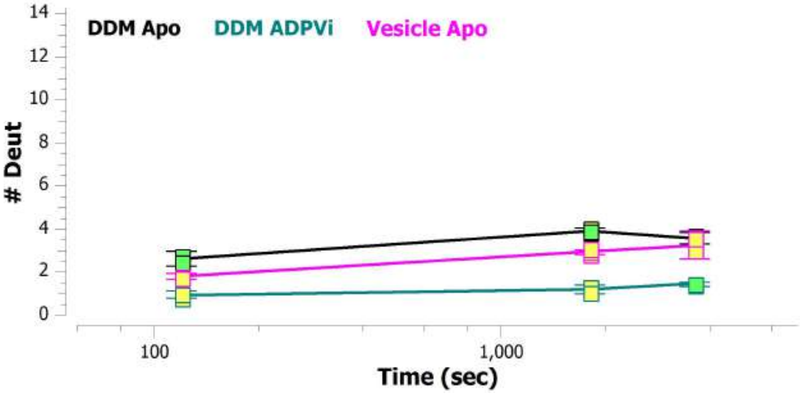


##### MSBA 225-241: VETKRFDKVSNRMRLQG (#41)

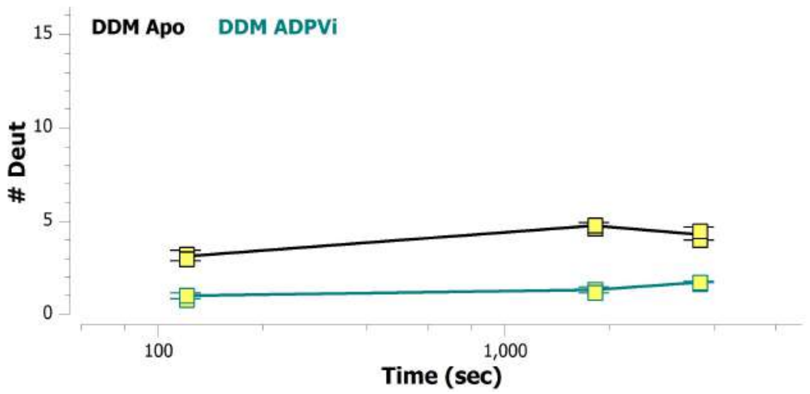


##### MSBA 225-244: VETKRFDKVSNRMRLQGMKM (#42)

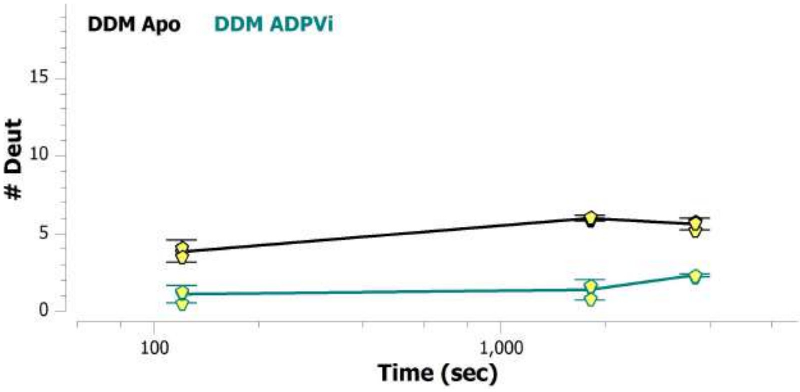


##### MSBA 227-235: TKRFDKVSN (#43)

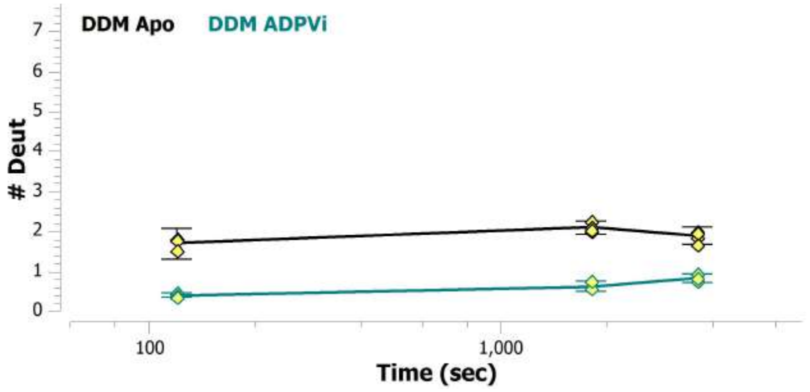

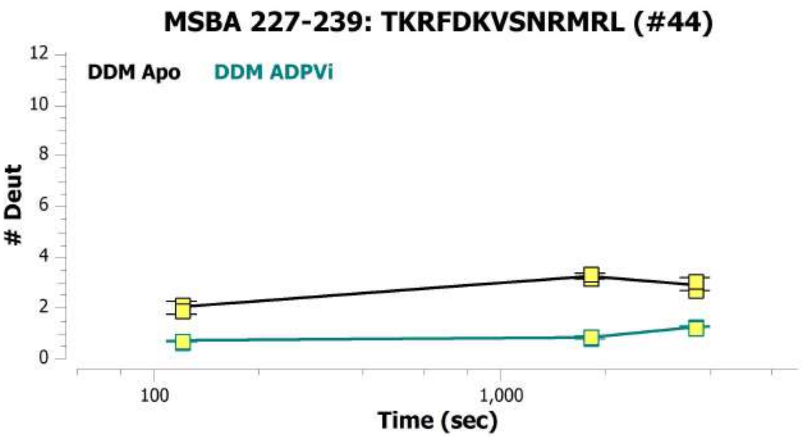
MSBA 227-239: TKRFDKVSNRMRL (#44)
DDM Apo
DDM ADPVi
### Deut
Time (sec)
100
1,000
0
2
4
6
8
10
12

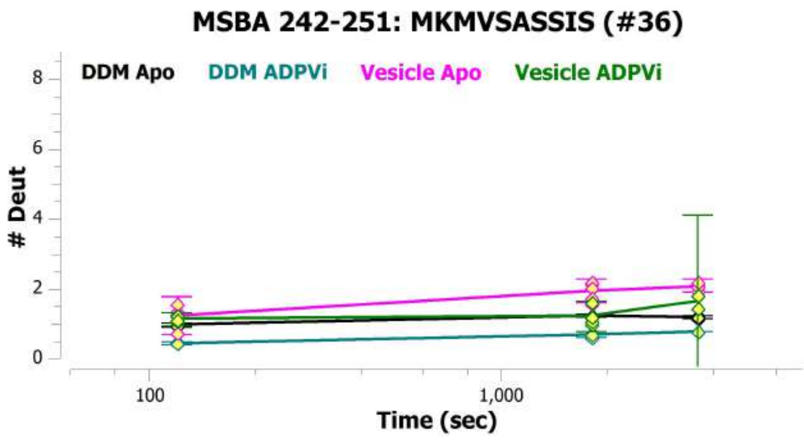
MSBA 242-251: MKMVSASSIS (#36)
DDM Apo
DDM ADPVi
Vesicle Apo
Vesicle ADPVi
### Deut
Time (sec)
100
1,000
0
2
4
6
8

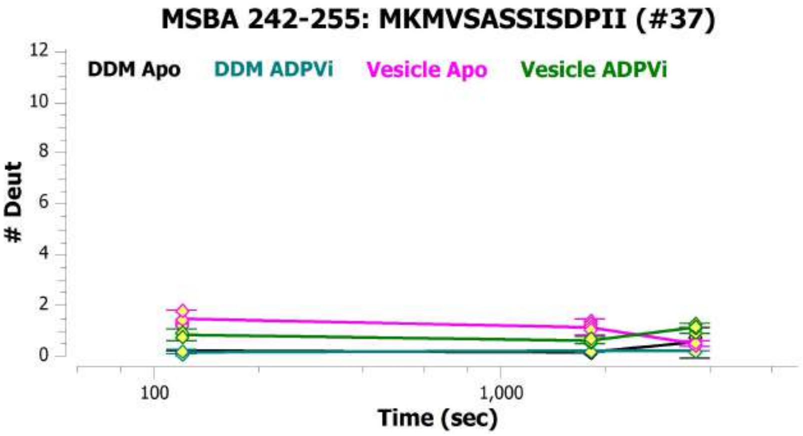
MSBA 242-255: MKMVSASSISDPII (#37)
DDM Apo
DDM ADPVi
Vesicle Apo
Vesicle ADPVi
### Deut
Time (sec)
100
1,000
0
2
4
6
8
10
12

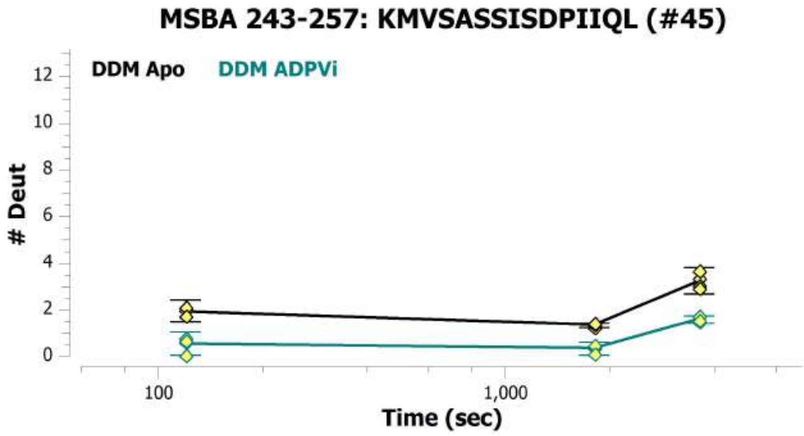
MSBA 243-257: KMVSASSISDPIIQL (#45)
DDM Apo
DDM ADPVi
### Deut
Time (sec)
100
1,000
0
2
4
6
8
10
12

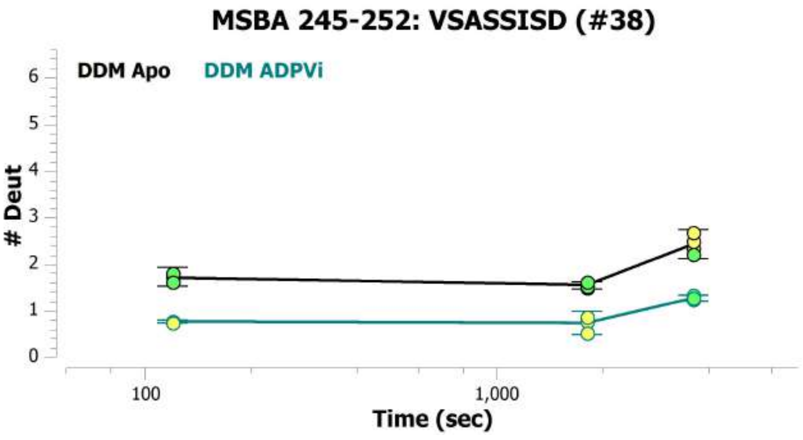
MSBA 245-252: VSASSISD (#38)
DDM Apo
DDM ADPVi
### Deut
Time (sec)
100
1,000
0
1
2
3
4
5
6

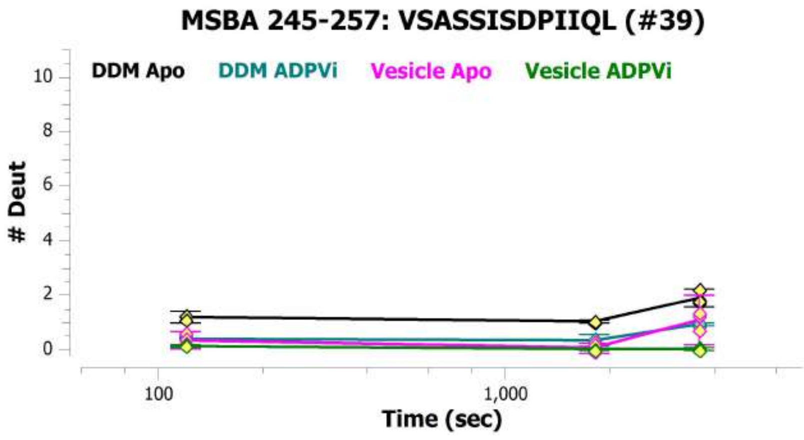
MSBA 245-257: VSASSISDPIIQL (#39)
DDM Apo
DDM ADPVi
Vesicle Apo
Vesicle ADPVi
### Deut
Time (sec)
100
1,000
0
2
4
6
8
10

MSBA 247-257: ASSISDPIIQL (#46)

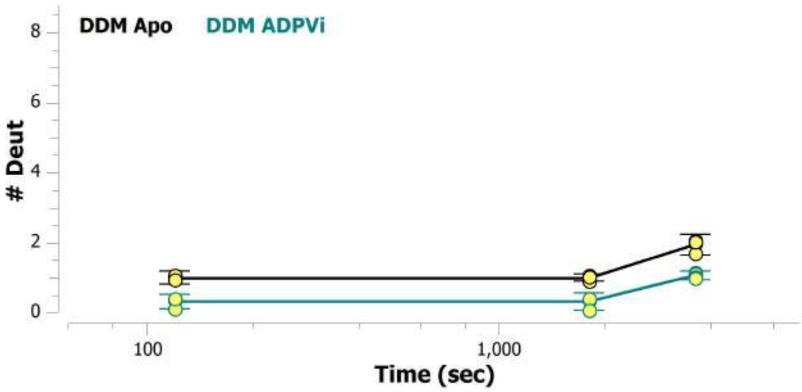


MSBA 248-257: SSISDPIIQL (#47)

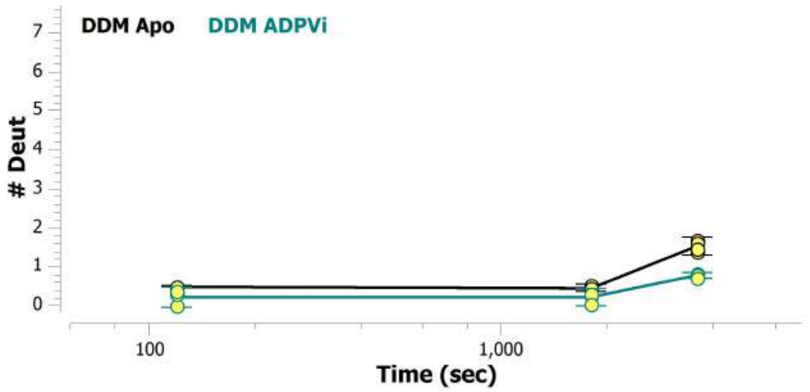


MSBA 250-257: ISDPIIQL (#48)

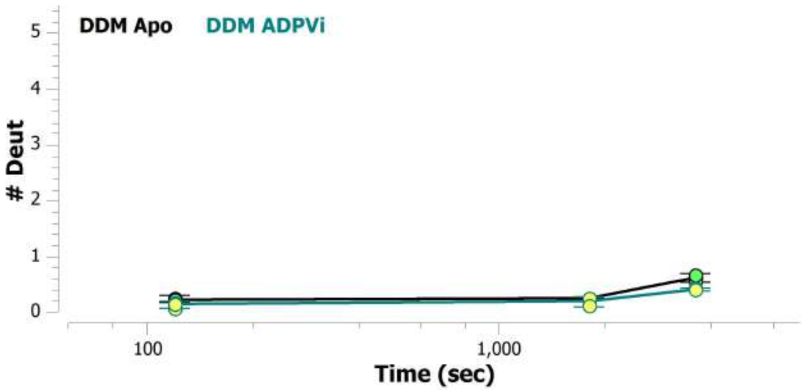


MSBA 258-263: IASLAL (#40)

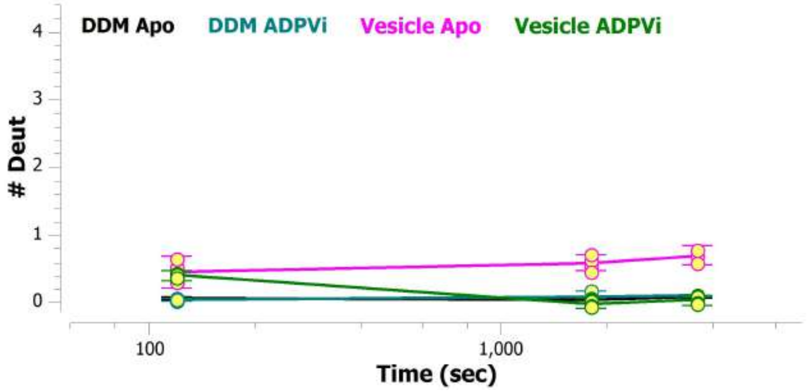


MSBA 258-264: IASLALA (#41)

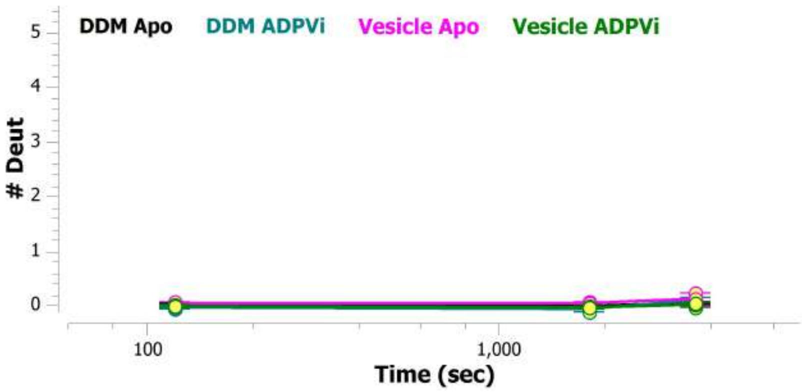


MSBA 258-265: IASLALAF (#42)

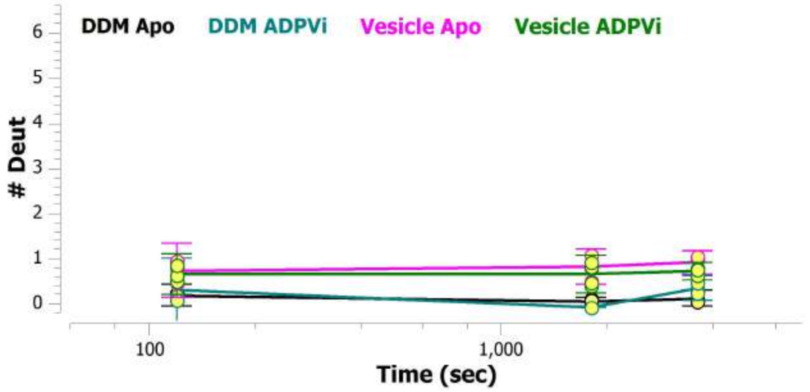

MSBA 258-277: IASLALAFVLYAASFPSVMD (#49)

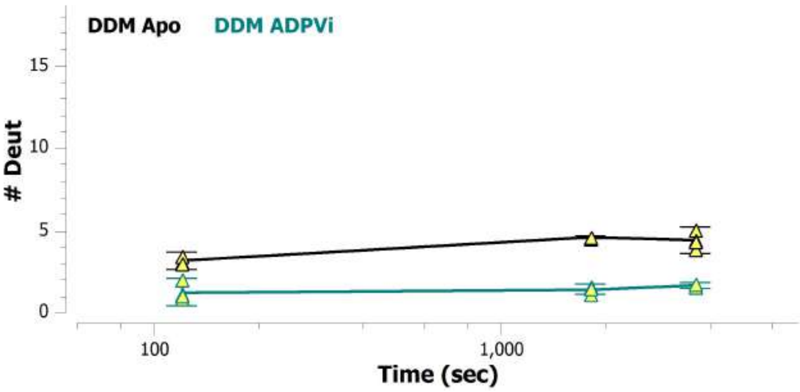


MSBA 266-272: VLYAASF (#43)

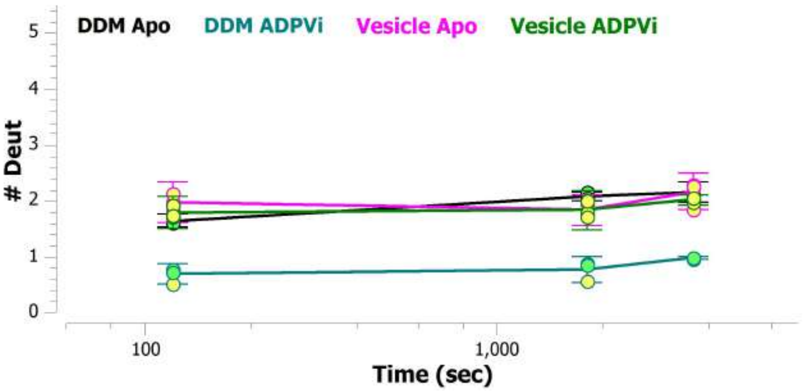


MSBA 268-276: YAASFPSVM (#50)

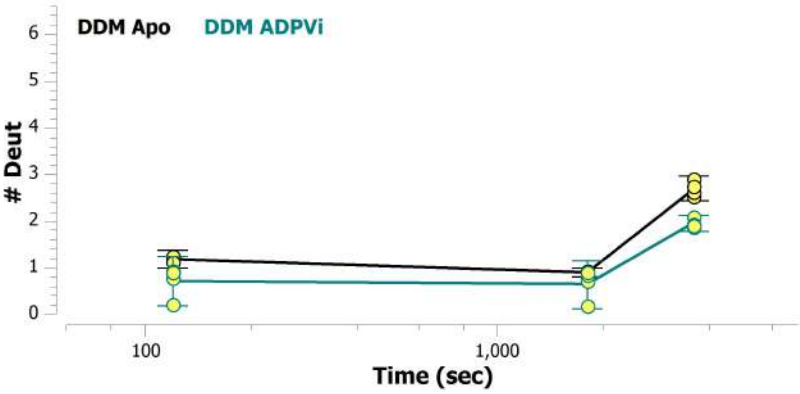


MSBA 269-276: AASFPSVM (#51)

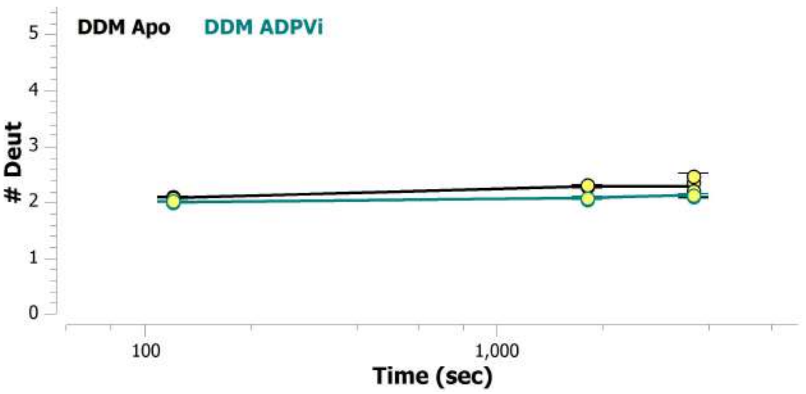


MSBA 270-276: ASFPSVM (#44)

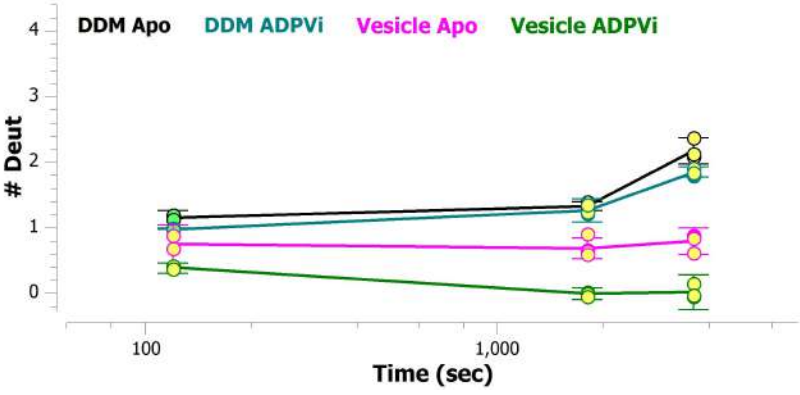


MSBA 277-282: DSLTAG (#45)

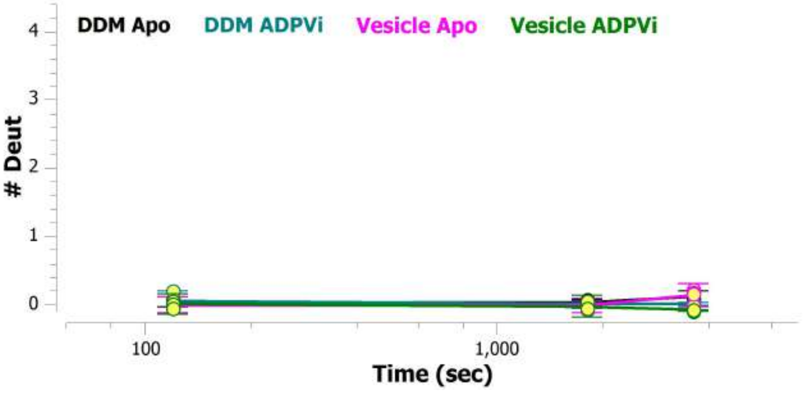

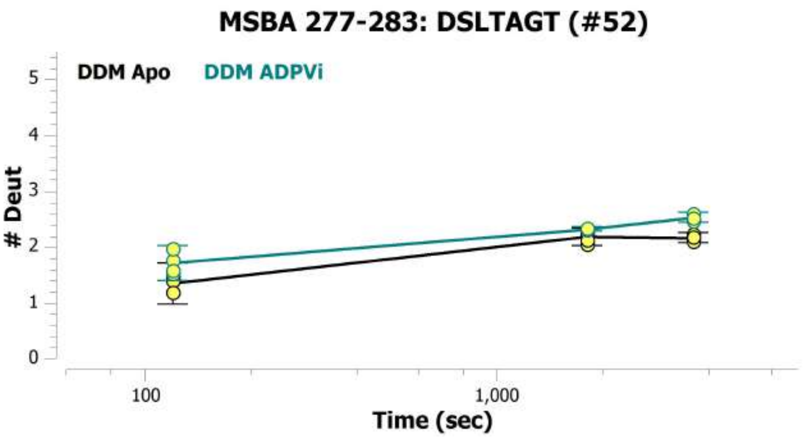

MSBA 277-283: DSLTAGT (#52)
DDM Apo
DDM ADPVi
### Deut
Time (sec)


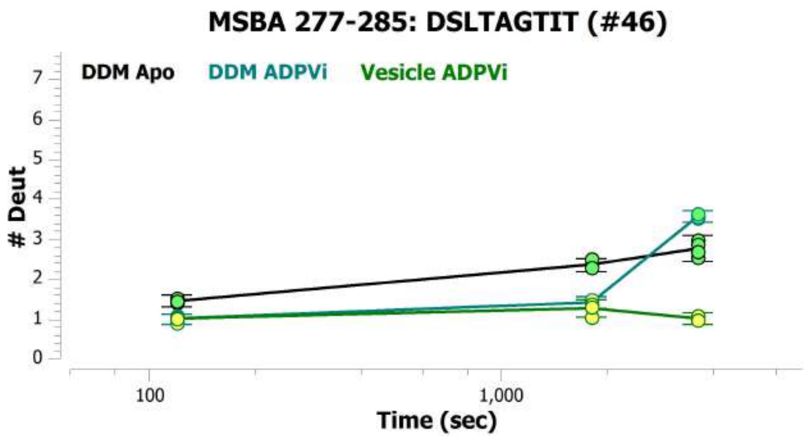

MSBA 277-285: DSLTAGTIT (#46)
DDM Apo
DDM ADPVi
Vesicle ADPVi
### Deut
Time (sec)


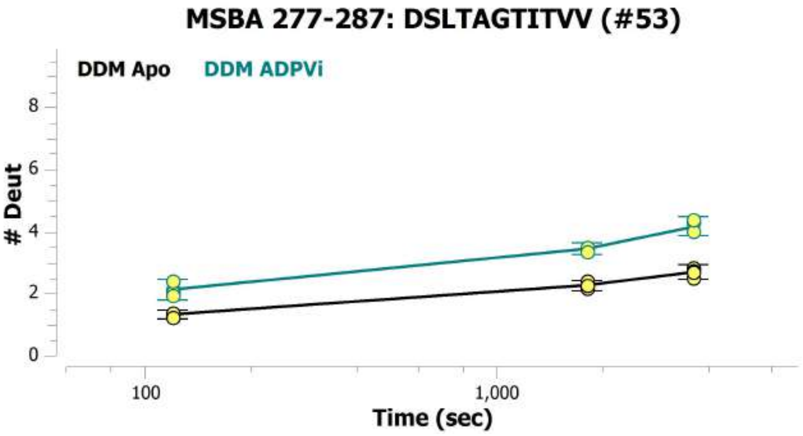

MSBA 277-287: DSLTAGTITVV (#53)
DDM Apo
DDM ADPVi
### Deut
Time (sec)


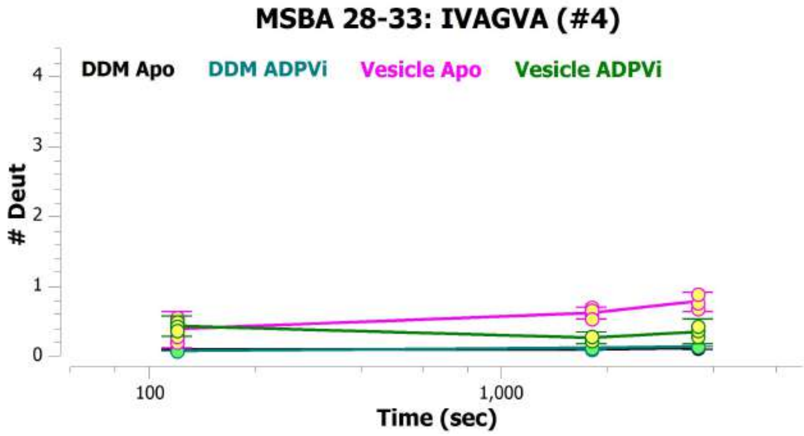

MSBA 28-33: IVAGVA (#4)
DDM Apo
DDM ADPVi
Vesicle Apo
Vesicle ADPVi
### Deut
Time (sec)


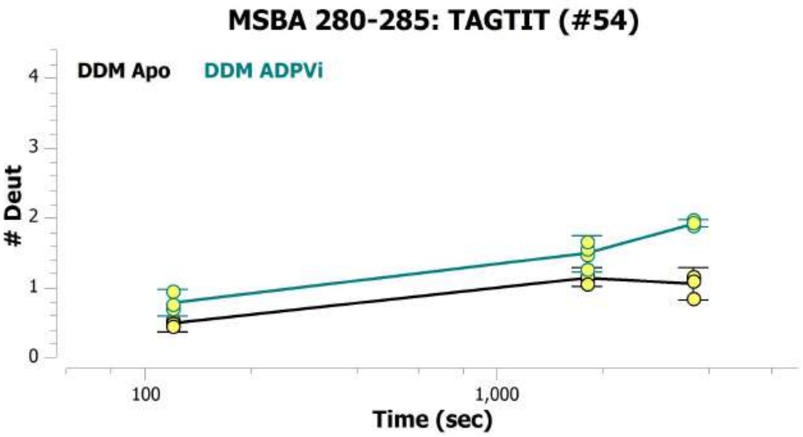

MSBA 280-285: TAGTIT (#54)
DDM Apo
DDM ADPVi
### Deut
Time (sec)


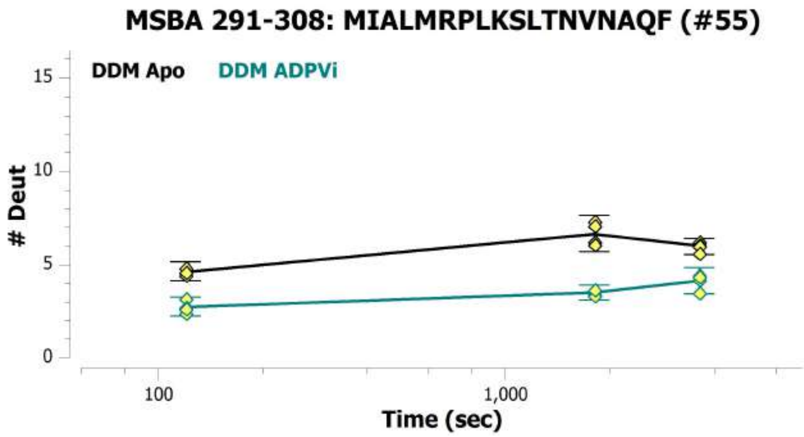

MSBA 291-308: MIALMRPLKSLTNVNAQF (#55)
DDM Apo
DDM ADPVi
### Deut
Time (sec)

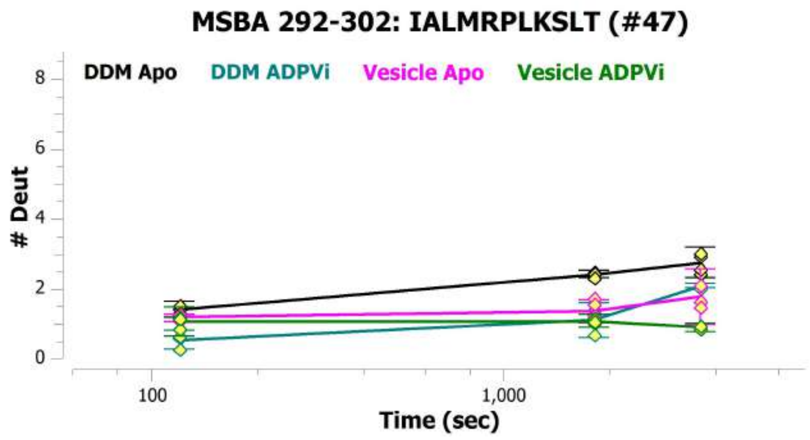
MSBA 292-302: IALMRPLKSLT (#47)
DDM Apo
DDM ADPVi
Vesicle Apo
Vesicle ADPVi
### Deut
Time (sec)
100
1,000

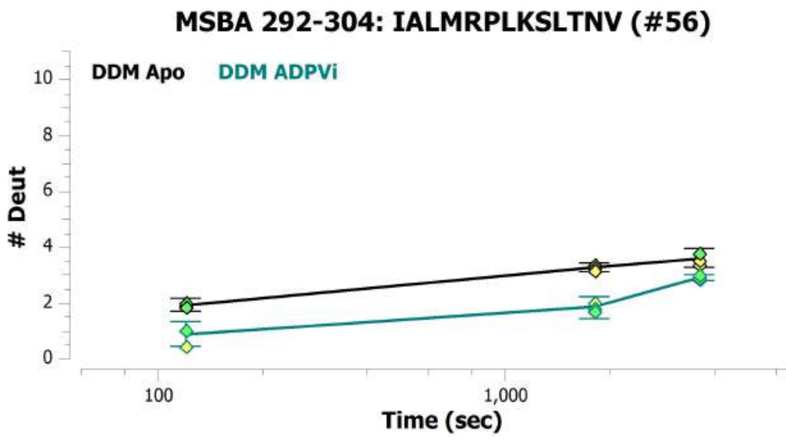
MSBA 292-304: IALMRPLKSLTNV (#56)
DDM Apo
DDM ADPVi
### Deut
Time (sec)
100
1,000

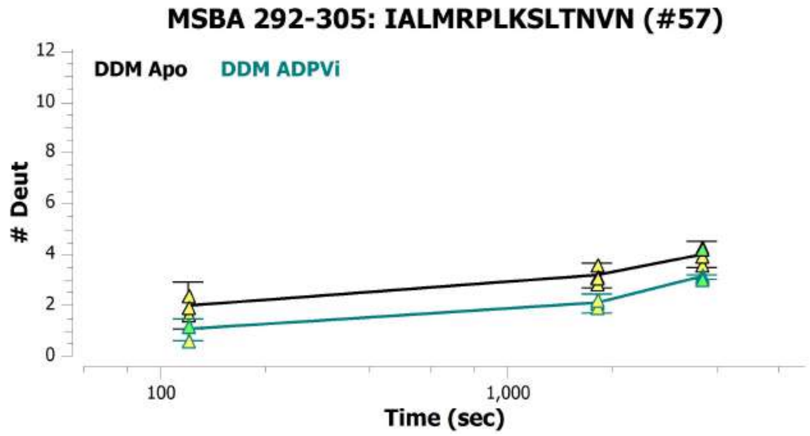
MSBA 292-305: IALMRPLKSLTNVN (#57)
DDM Apo
DDM ADPVi
### Deut
Time (sec)
100
1,000

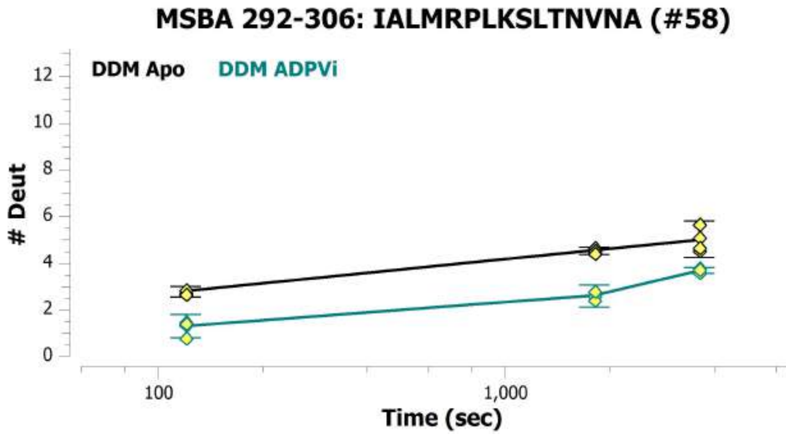
MSBA 292-306: IALMRPLKSLTNVNA (#58)
DDM Apo
DDM ADPVi
### Deut
Time (sec)
100
1,000

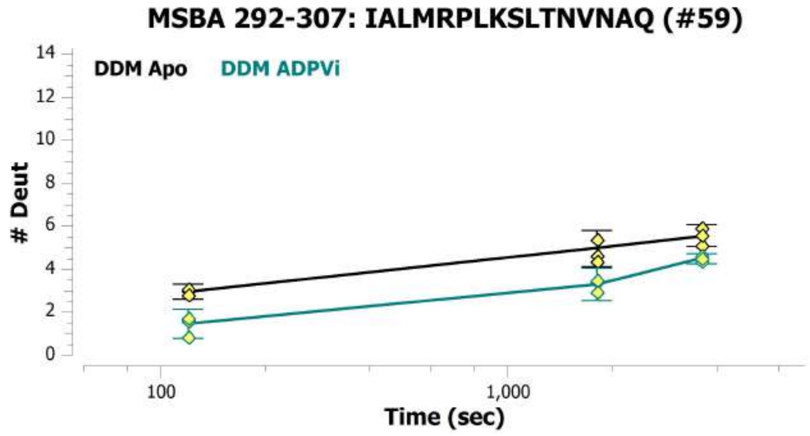
MSBA 292-307: IALMRPLKSLTNVNAQ (#59)
DDM Apo
DDM ADPVi
### Deut
Time (sec)
100
1,000

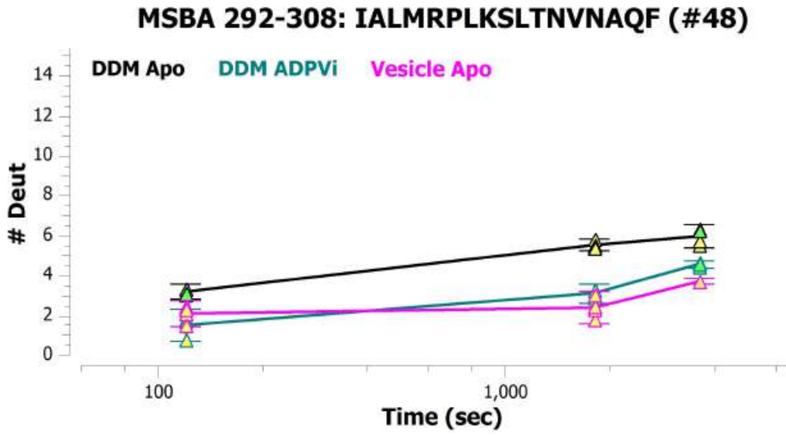
MSBA 292-308: IALMRPLKSLTNVNAQF (#48)
DDM Apo
DDM ADPVi
Vesicle Apo
### Deut
Time (sec)
100
1,000

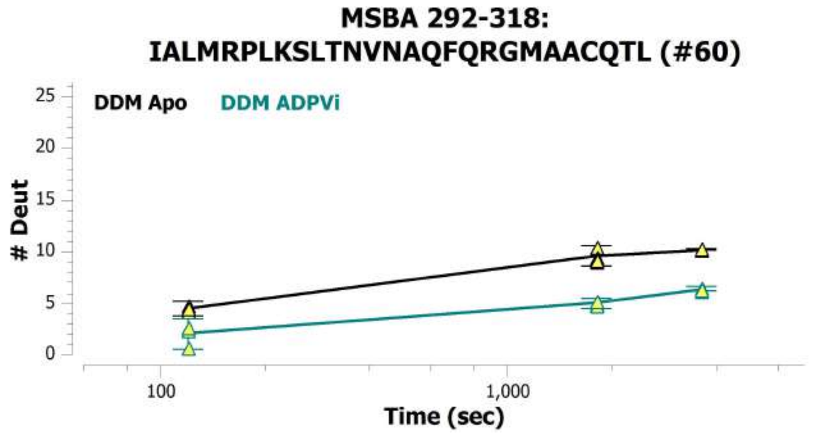
MSBA 292-318:
IALMRPLKSLTNVNAQFQRGMAACQTL (#60)
DDM Apo
DDM ADPVi
### Deut
Time (sec)
0
5
10
15
20
25
100
1,000

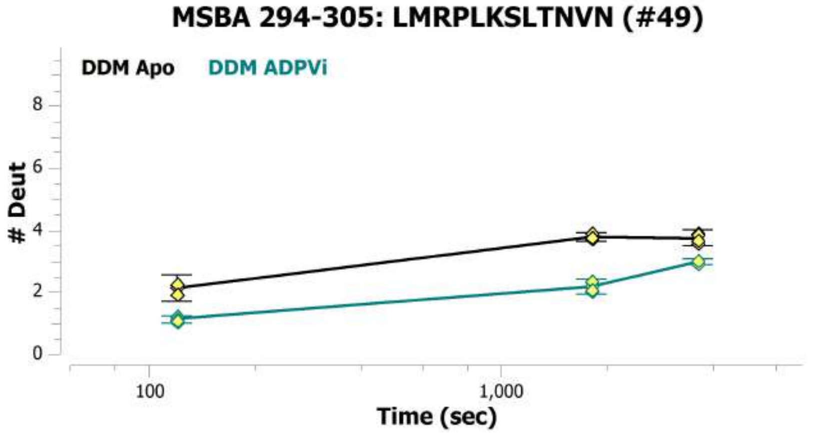
MSBA 294-305: LMRPLKSLTNVN (#49)
DDM Apo
DDM ADPVi
### Deut
Time (sec)
0
2
4
6
8
100
1,000

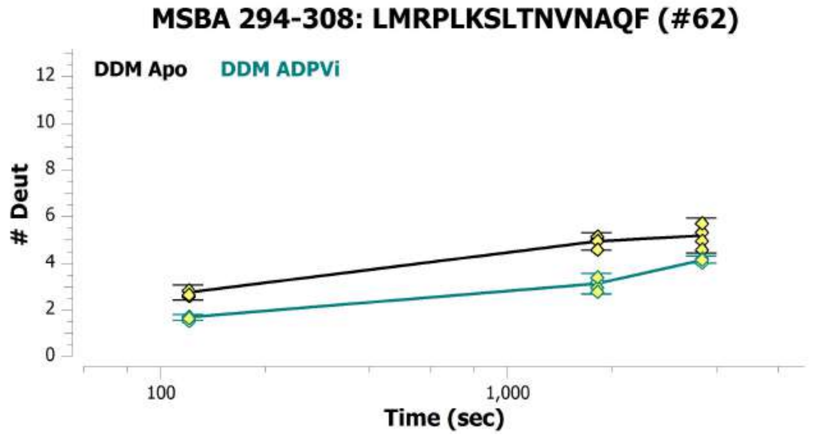
MSBA 294-308: LMRPLKSLTNVNAQF (#62)
DDM Apo
DDM ADPVi
### Deut
Time (sec)
0
2
4
6
8
10
12
100
1,000

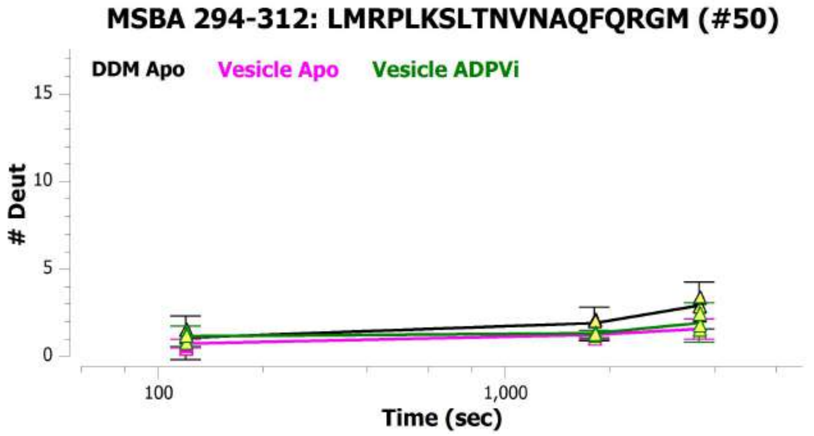
MSBA 294-312: LMRPLKSLTNVNAQFQRGM (#50)
DDM Apo
Vesicle Apo
Vesicle ADPVi
### Deut
Time (sec)
0
5
10
15
100
1,000

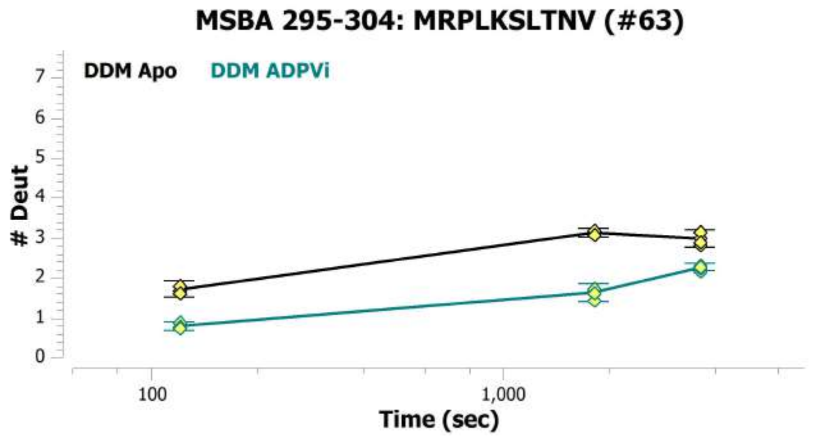
MSBA 295-304: MRPLKSLTNV (#63)
DDM Apo
DDM ADPVi
### Deut
Time (sec)
0
1
2
3
4
5
6
7
100
1,000

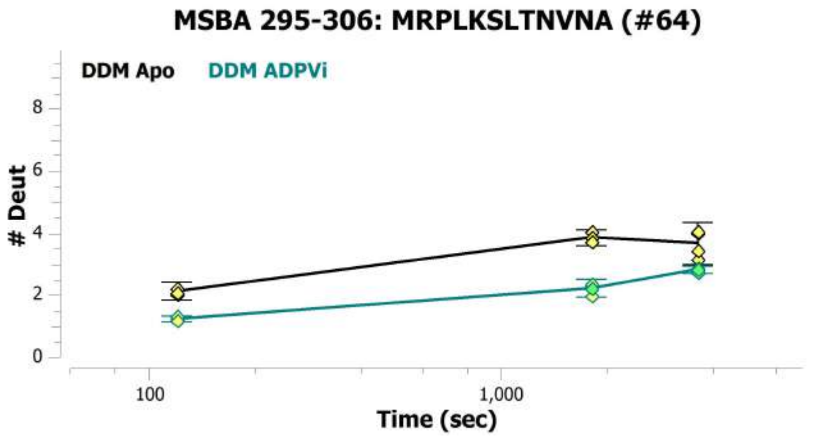
MSBA 295-306: MRPLKSLTNVNA (#64)
DDM Apo
DDM ADPVi
### Deut
Time (sec)
0
2
4
6
8
100
1,000

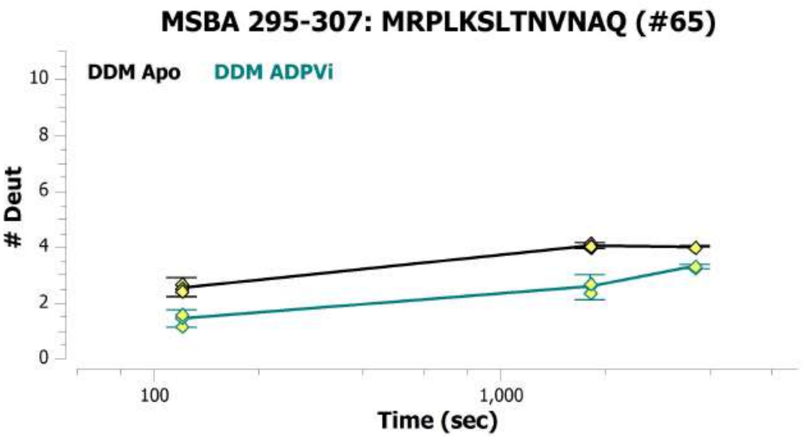
MSBA 295-307: MRPLKSLTNVNAQ (#65)
DDM Apo
DDM ADPVi
### Deut
Time (sec)

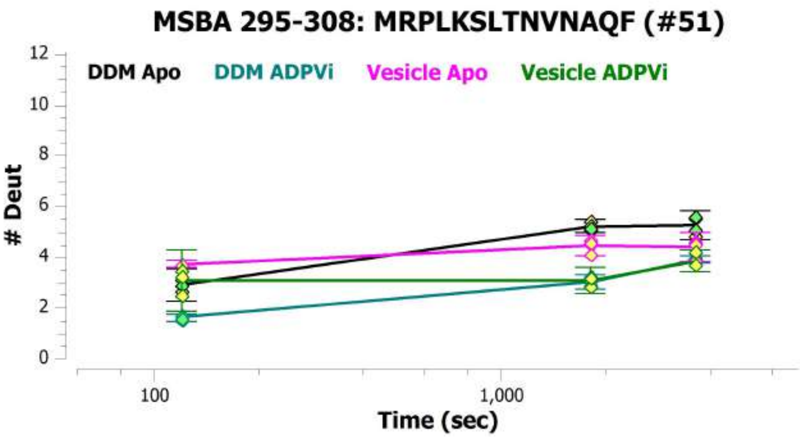
MSBA 295-308: MRPLKSLTNVNAQF (#51)
DDM Apo
DDM ADPVi
Vesicle Apo
Vesicle ADPVi
### Deut
Time (sec)

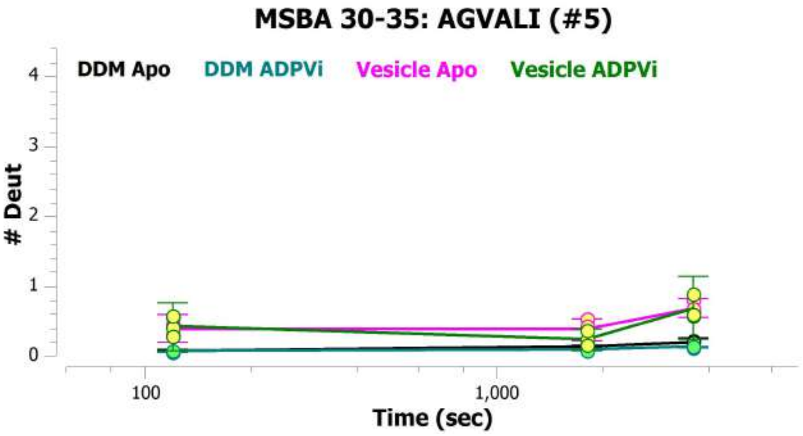
MSBA 30-35: AGVALI (#5)
DDM Apo
DDM ADPVi
Vesicle Apo
Vesicle ADPVi
### Deut
Time (sec)

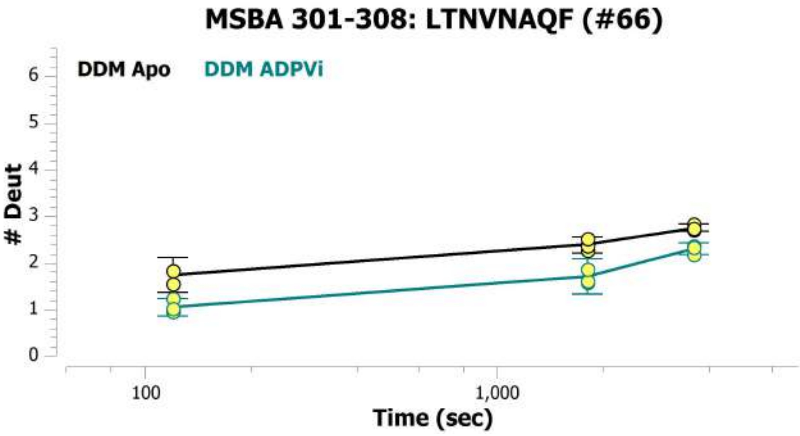
MSBA 301-308: LTNVNAQF (#66)
DDM Apo
DDM ADPVi
### Deut
Time (sec)

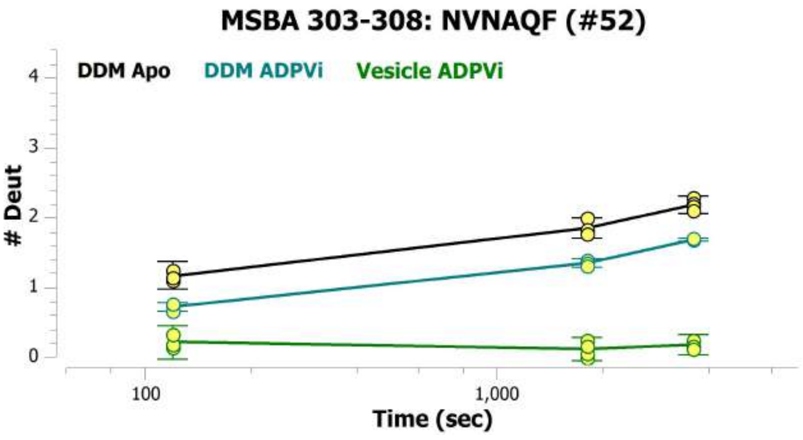
MSBA 303-308: NVNAQF (#52)
DDM Apo
DDM ADPVi
Vesicle ADPVi
### Deut
Time (sec)

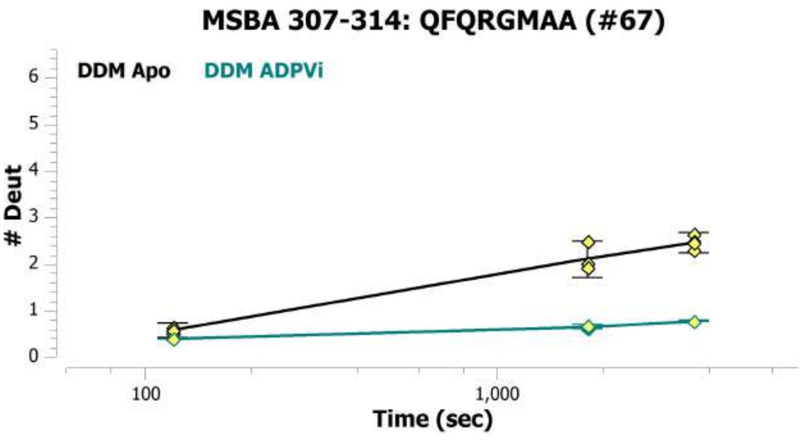
MSBA 307-314: QFQRGMAA (#67)
DDM Apo
DDM ADPVi
### Deut
Time (sec)

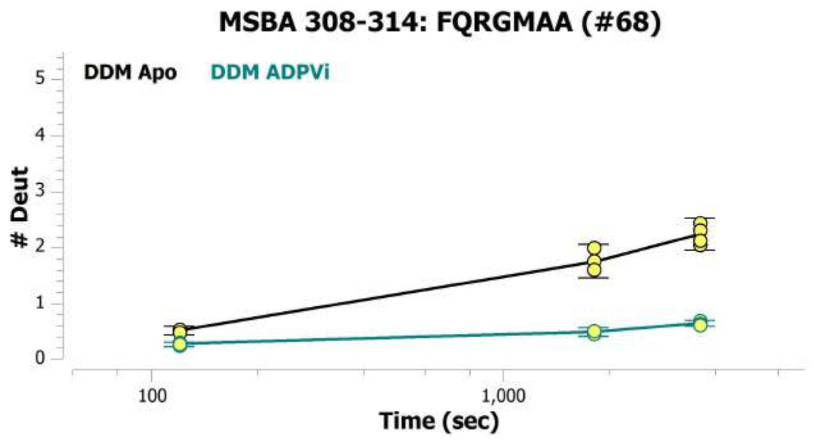
MSBA 308-314: FQRGMAA (#68)
DDM Apo
DDM ADPVi
### Deut
0
1
2
3
4
5
100
1,000
Time (sec)

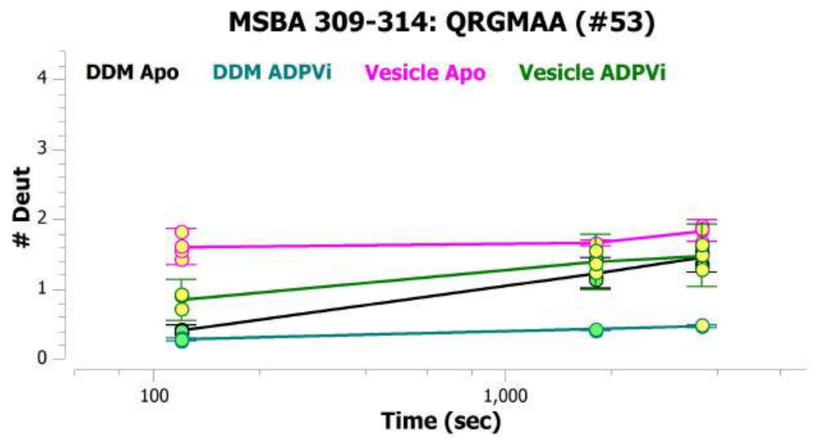
MSBA 309-314: QRGMAA (#53)
DDM Apo
DDM ADPVi
Vesicle Apo
Vesicle ADPVi
### Deut
0
1
2
3
4
100
1,000
Time (sec)

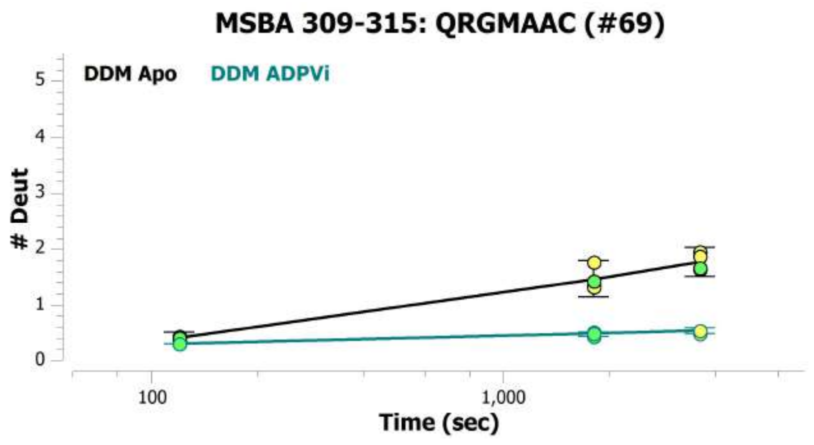
MSBA 309-315: QRGMAAC (#69)
DDM Apo
DDM ADPVi
### Deut
0
1
2
3
4
5
100
1,000
Time (sec)

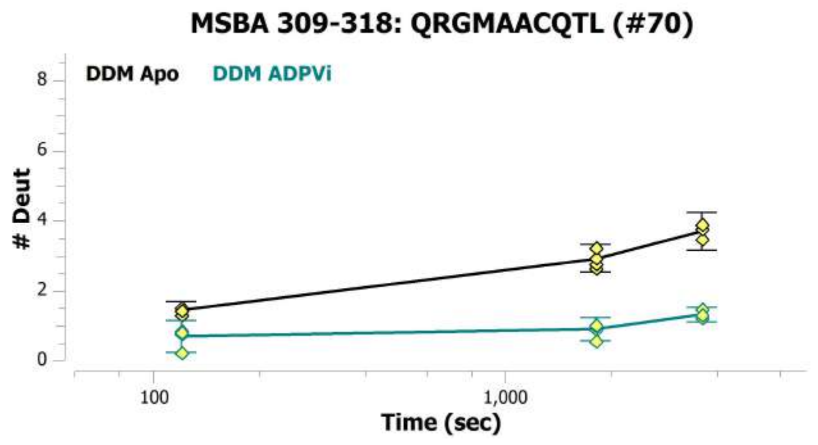
MSBA 309-318: QRGMAACQTL (#70)
DDM Apo
DDM ADPVi
### Deut
0
2
4
6
8
100
1,000
Time (sec)

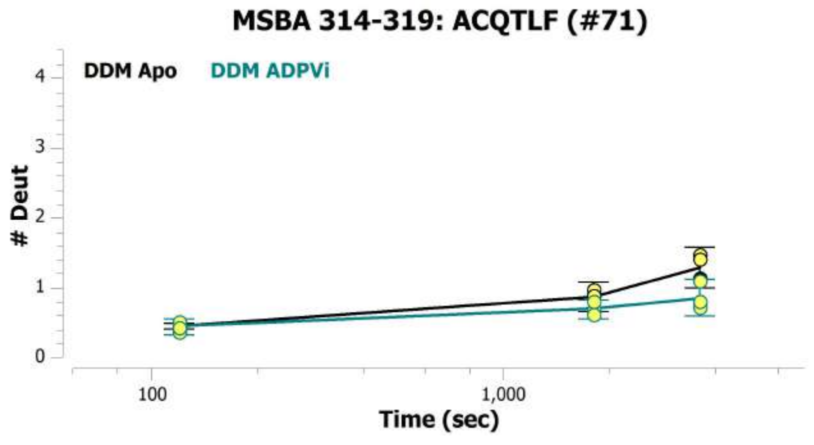
MSBA 314-319: ACQTLF (#71)
DDM Apo
DDM ADPVi
### Deut
0
1
2
3
4
100
1,000
Time (sec)

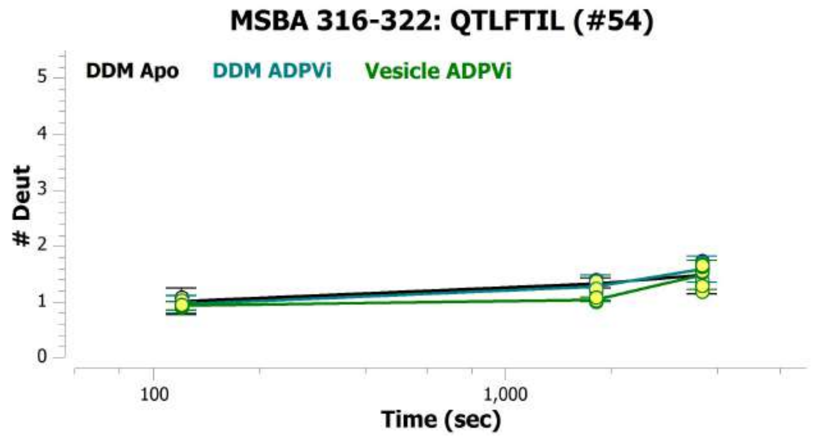
MSBA 316-322: QTLFTIL (#54)
DDM Apo
DDM ADPVi
Vesicle ADPVi
### Deut
0
1
2
3
4
5
100
1,000
Time (sec)

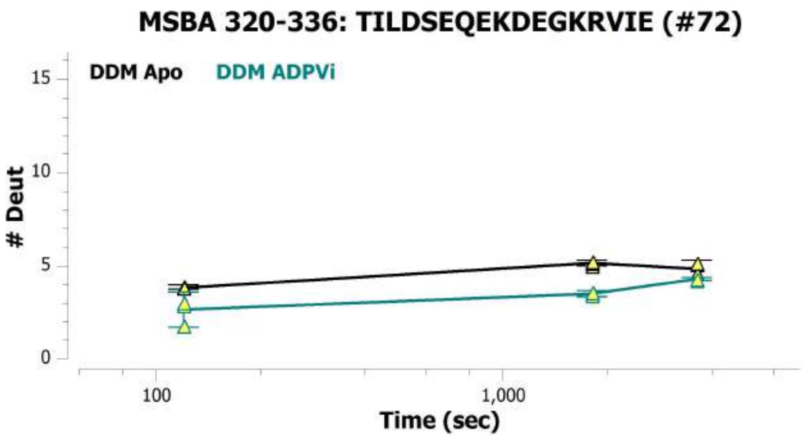

MSBA 320-336: TILDSEQEKDEGKRVIE (#72)
DDM Apo
DDM ADPVi
### Deut
Time (sec)


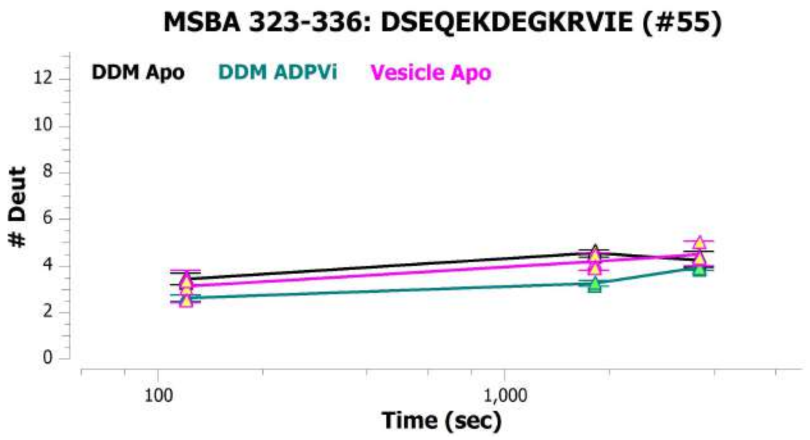

MSBA 323-336: DSEQEKDEGKRVIE (#55)
DDM Apo
DDM ADPVi
Vesicle Apo
### Deut
Time (sec)


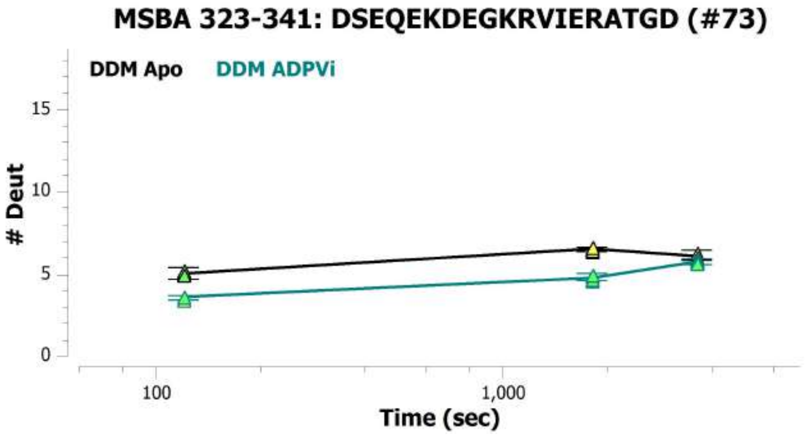

MSBA 323-341: DSEQEKDEGKRVIERATGD (#73)
DDM Apo
DDM ADPVi
### Deut
Time (sec)


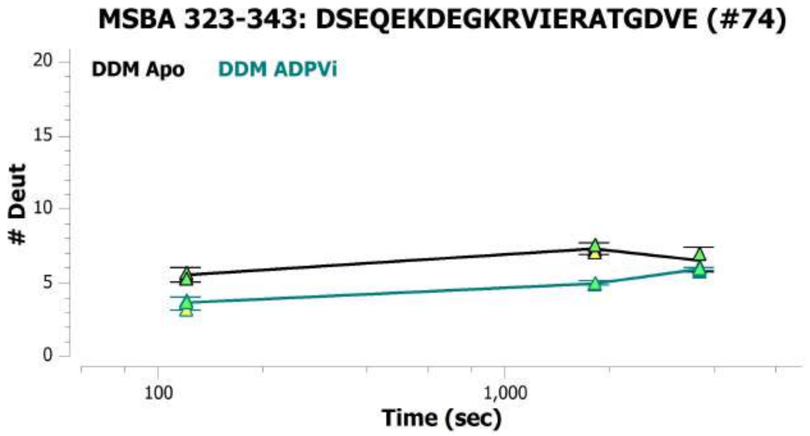

MSBA 323-343: DSEQEKDEGKRVIERATGDVE (#74)
DDM Apo
DDM ADPVi
### Deut
Time (sec)


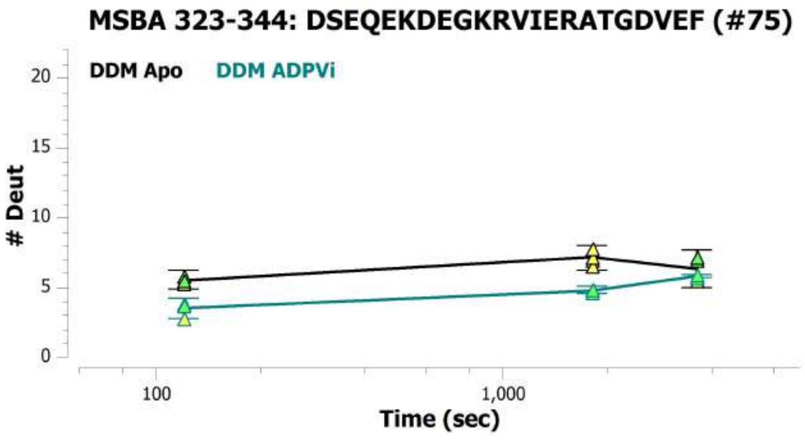

MSBA 323-344: DSEQEKDEGKRVIERATGDVEF (#75)
DDM Apo
DDM ADPVi
### Deut
Time (sec)


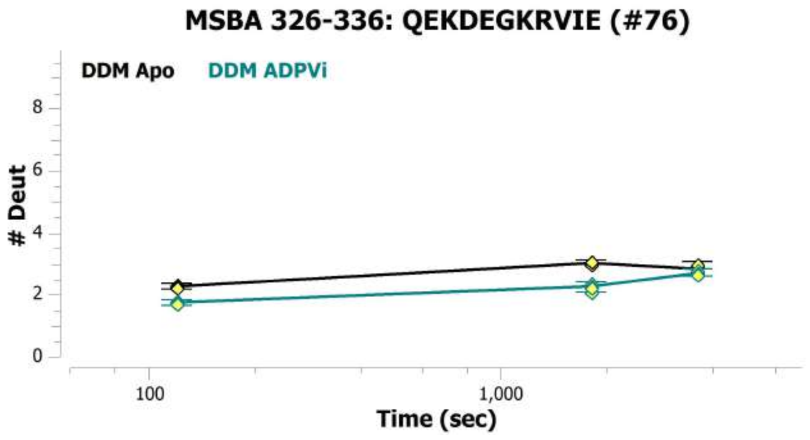

MSBA 326-336: QEKDEGKRVIE (#76)
DDM Apo
DDM ADPVi
### Deut
Time (sec)

MSBA 327-343: EKDEGKRVIERATGDVE (#77)

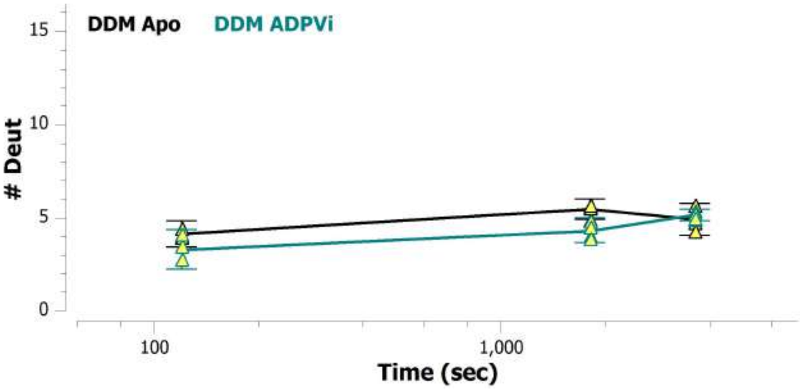


MSBA 327-344: EKDEGKRVIERATGDVEF (#78)

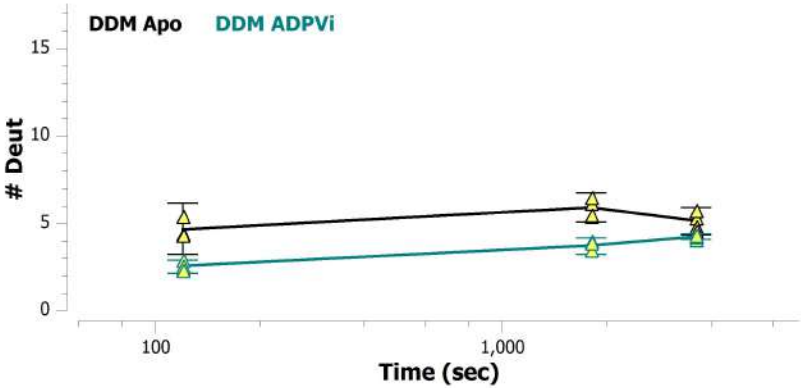


MSBA 337-344: RATGDVEF (#56)

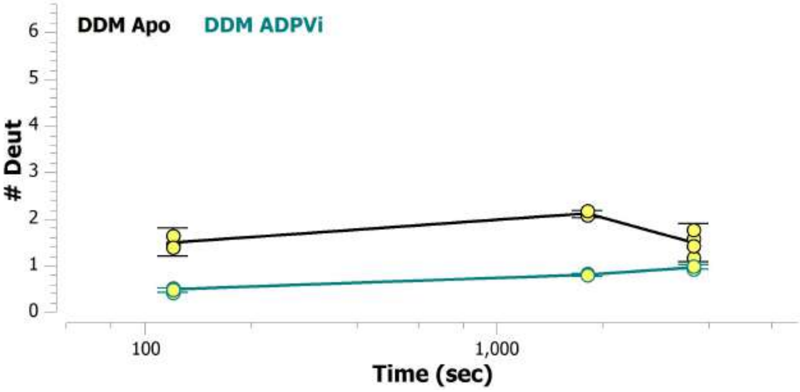


MSBA 34-44: LILNAASDTFM (#2)

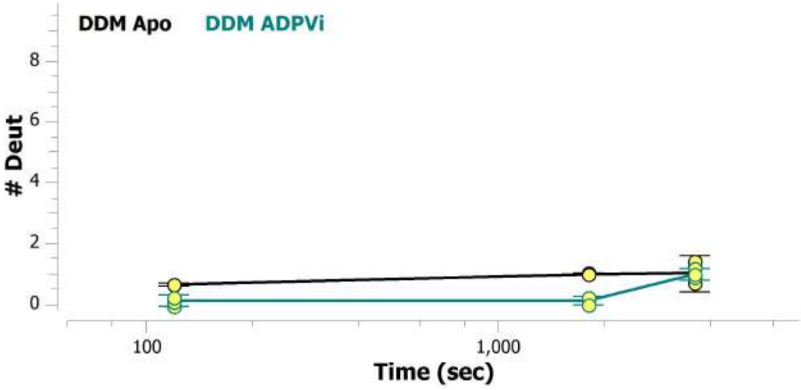


MSBA 344-349: FRNVTF (#79)

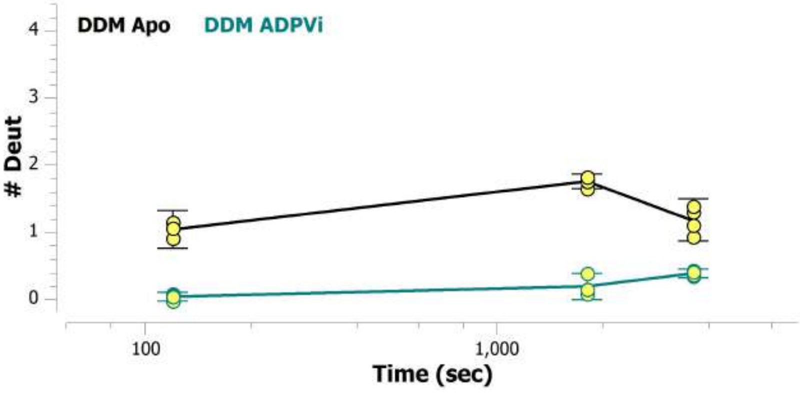


MSBA 344-359: FRNVTFTYPGRDVPAL (#57)

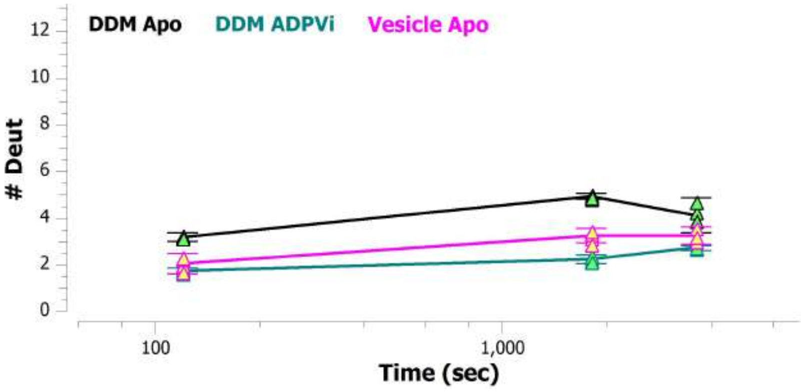

MSBA 344-361: FRNVTFTYPGRDVPALRN (#58)

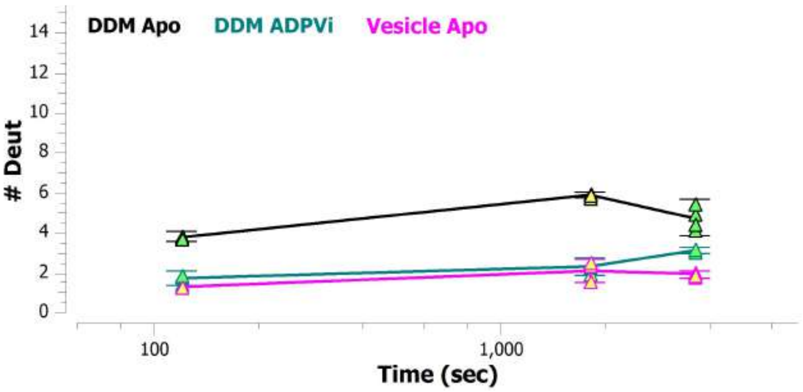


MSBA 344-363: FRNVTFTYPGRDVPALRNIN (#59)

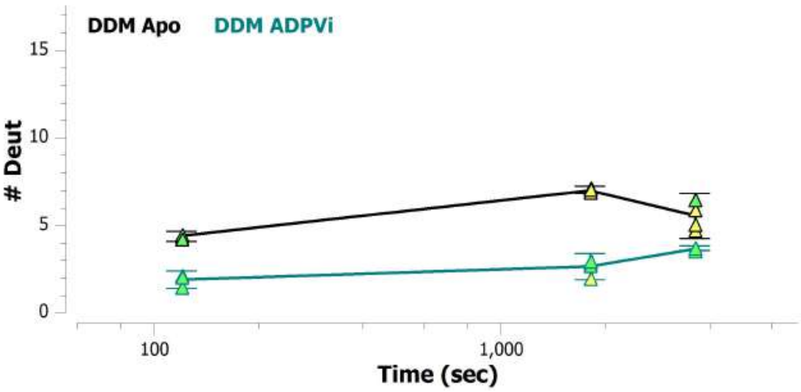


MSBA 345-359: RNVTFTYPGRDVPAL (#60)

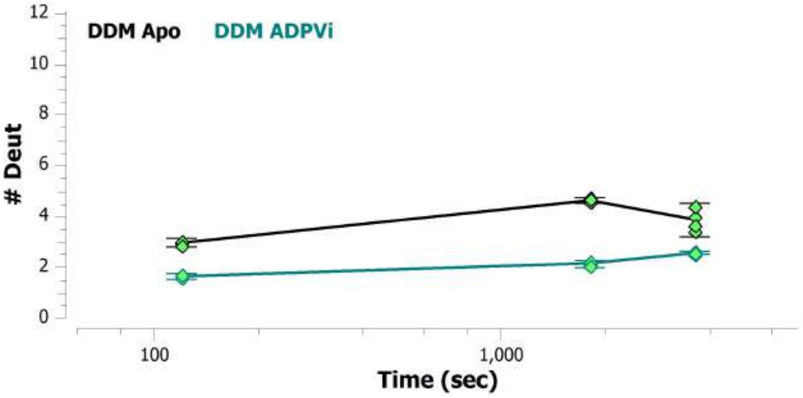


MSBA 345-361: RNVTFTYPGRDVPALRN (#80)

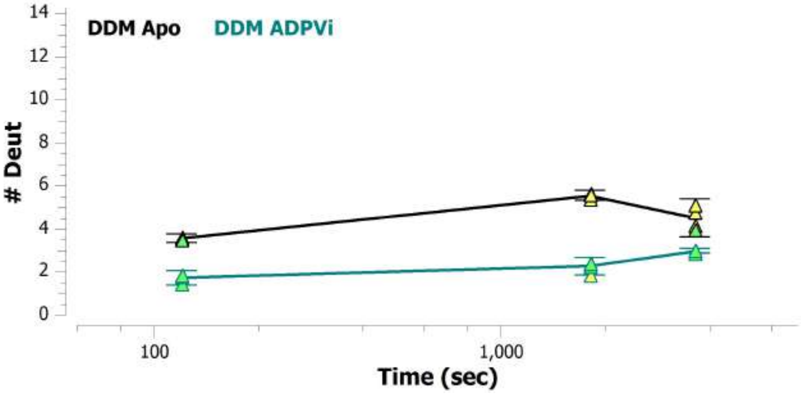


MSBA 345-363: RNVTFTYPGRDVPALRNIN (#61)

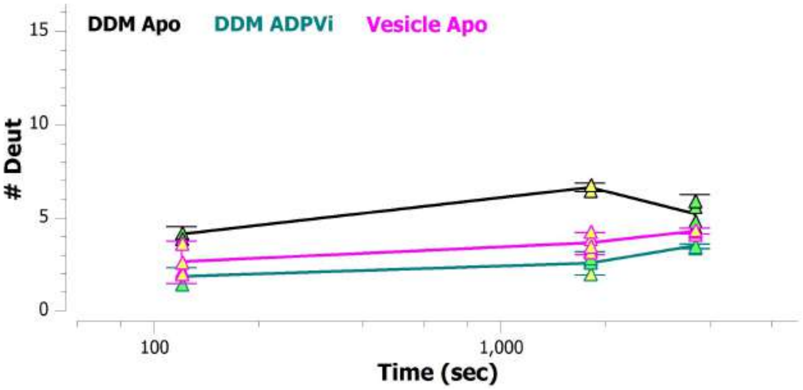


MSBA 345-374:
RNVTFTYPGRDVPALRNINLKIPAGKTVAL (#81)

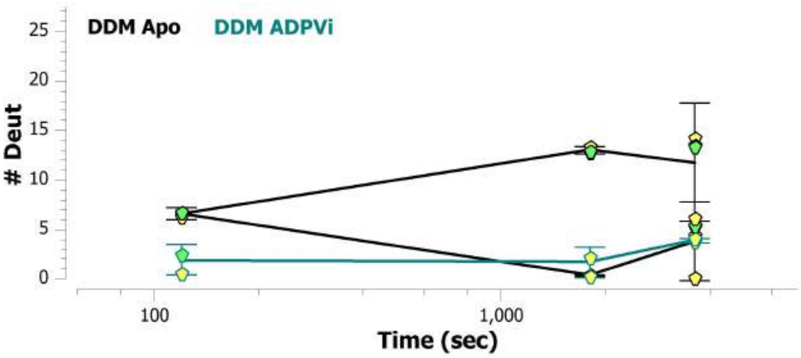

MSBA 347-361: VTFTYPGRDVPALRN (#62)

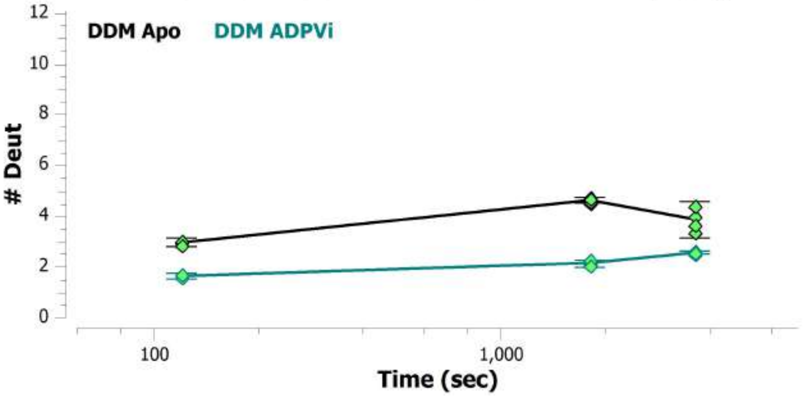


MSBA 349-359: FTYPGRDVPAL (#63)

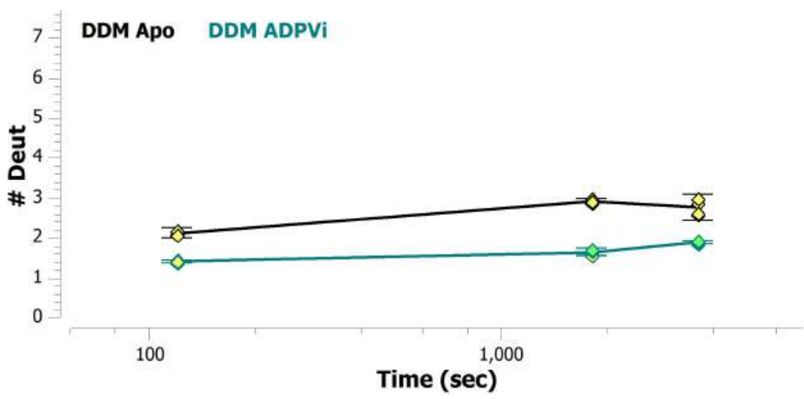


MSBA 349-363: FTYPGRDVPALRNIN (#82)

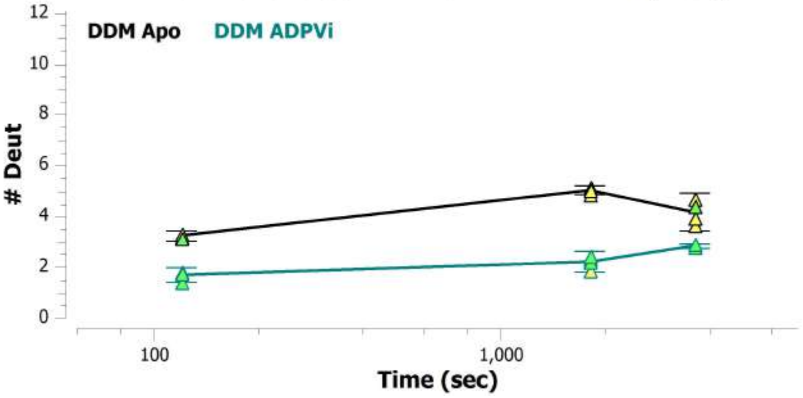


MSBA 35-44: ILNAASDTFM (#3)

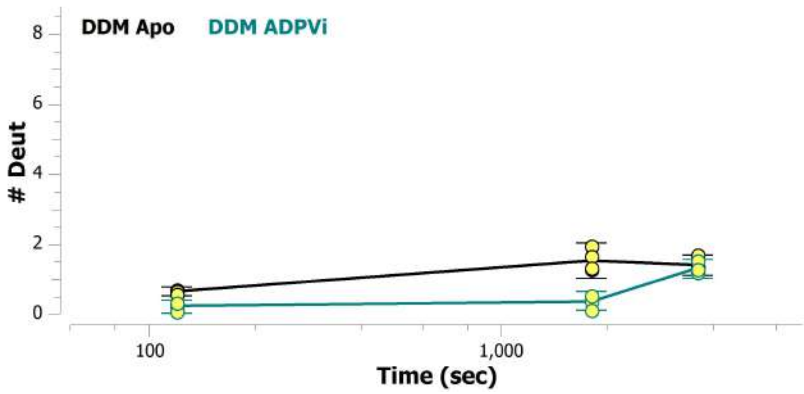


MSBA 350-363: TYPGRDVPALRNIN (#83)

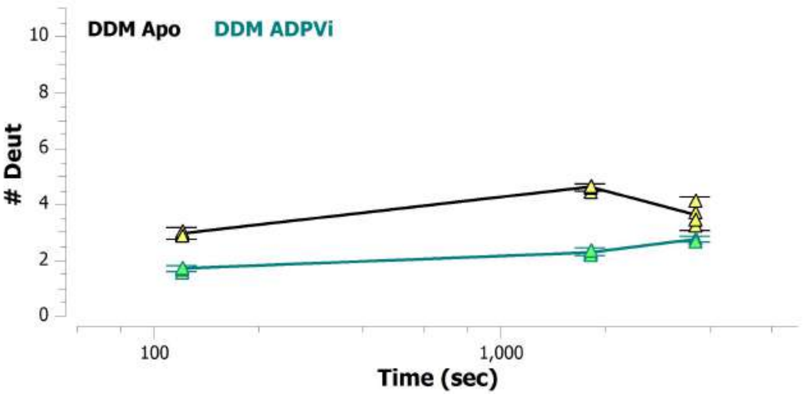


MSBA 351-363: YPGRDVPALRNIN (#84)

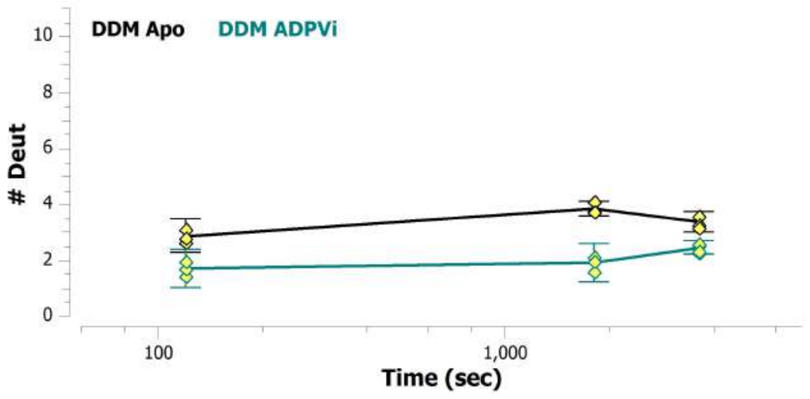

MSBA 352-363: PGRDVPALRNIN (#85)

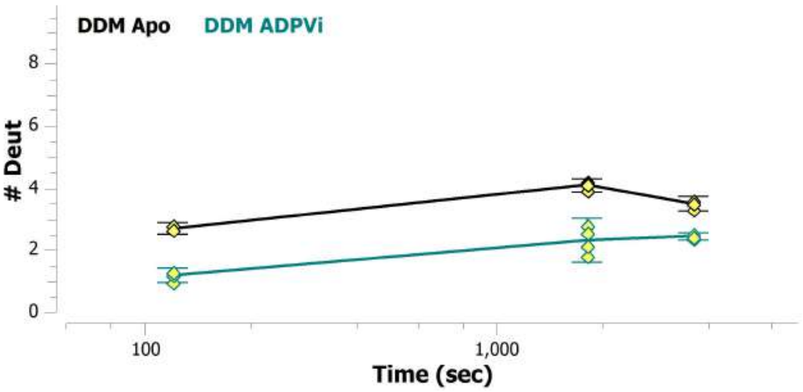


MSBA 360-374: RNINLKIPAGKTVAL (#86)

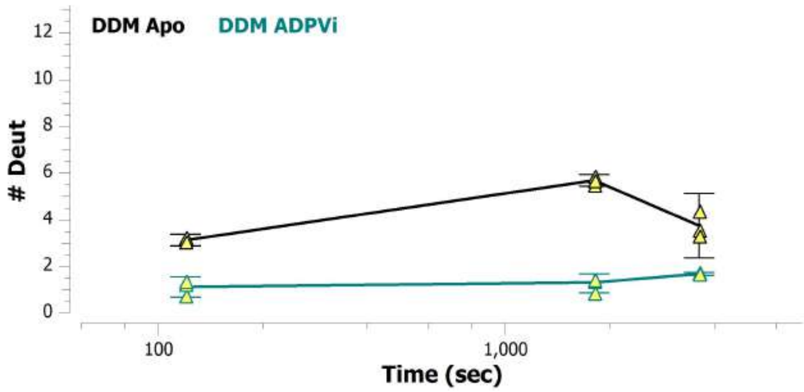


MSBA 362-374: INLKIPAGKTVAL (#64)

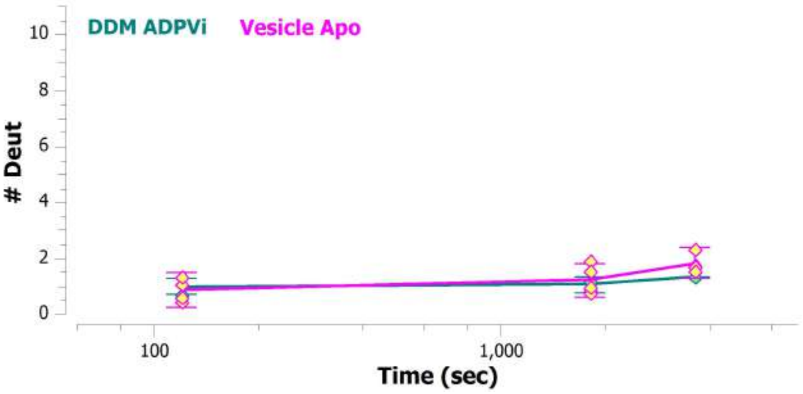


MSBA 363-374: NLKIPAGKTVAL (#87)

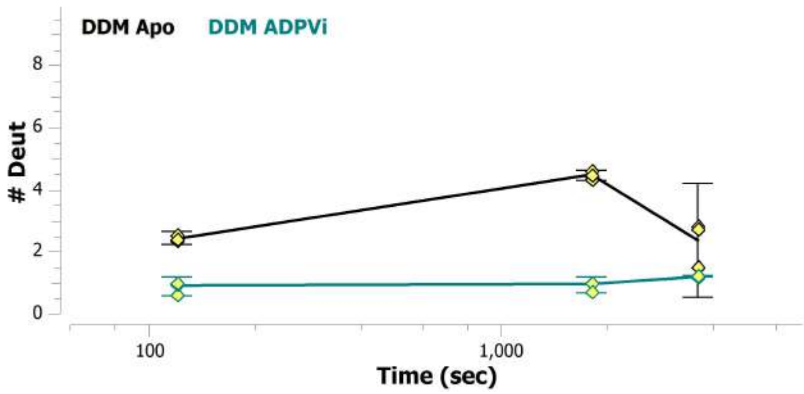


MSBA 364-371: LKIPAGKT (#88)

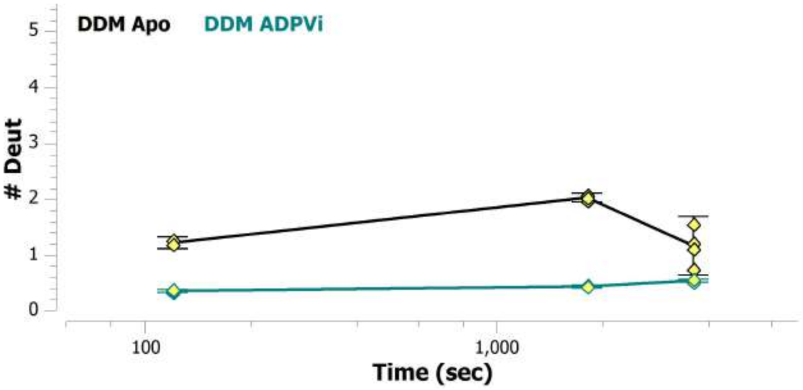


MSBA 364-373: LKIPAGKTVA (#89)

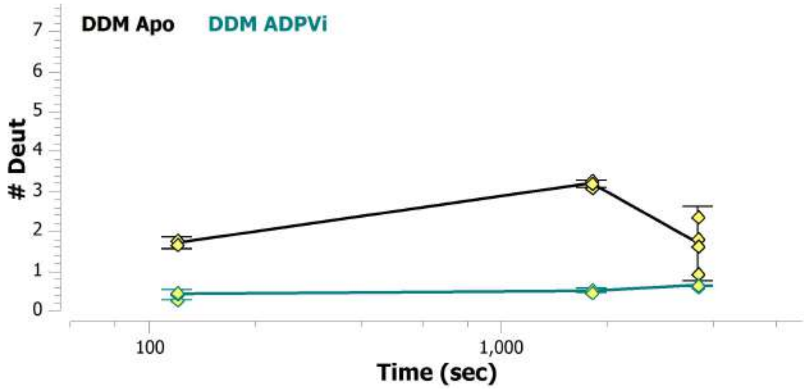

MSBA 364-374: LKIPAGKTVAL (#65)

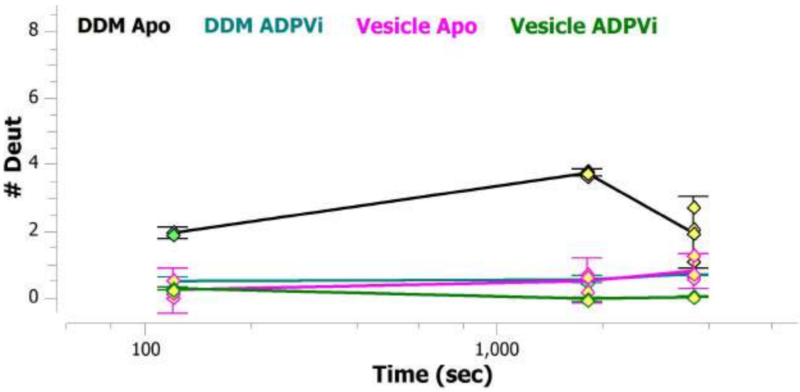


MSBA 365-374: KIPAGKTVAL (#90)

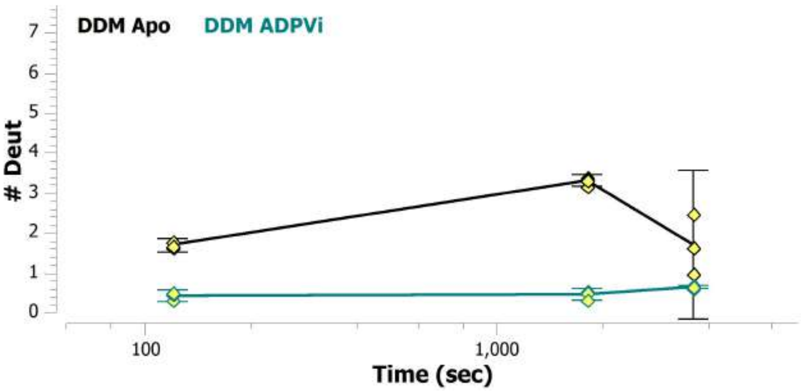


MSBA 369-374: GKTVAL (#91)

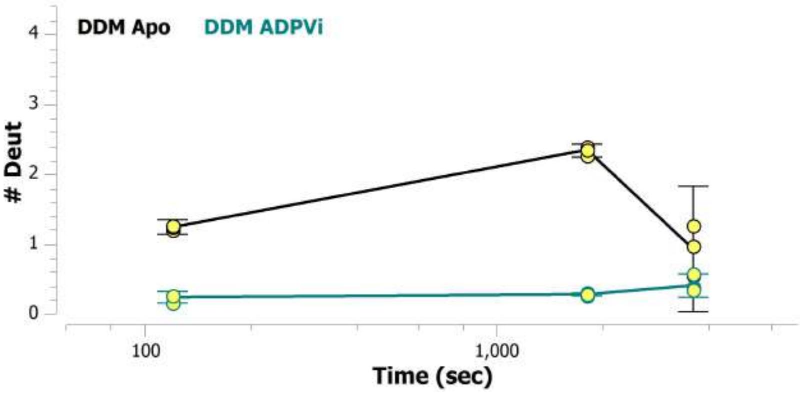


MSBA 37-43: NAASDTF (#4)

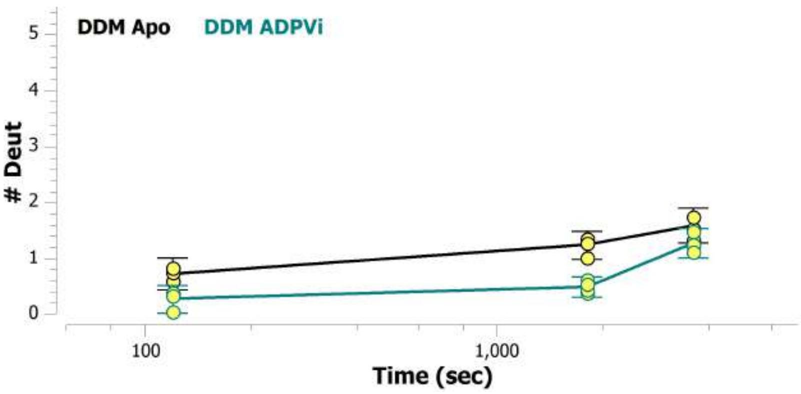


MSBA 37-44: NAASDTFM (#5)

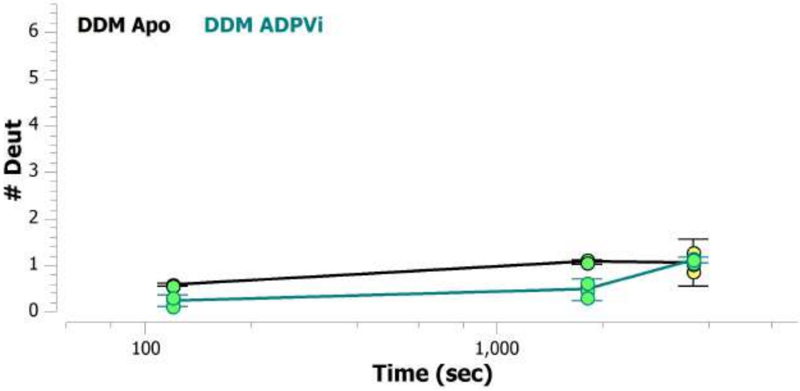


MSBA 371-382: TVALVGRSGSGK (#66)

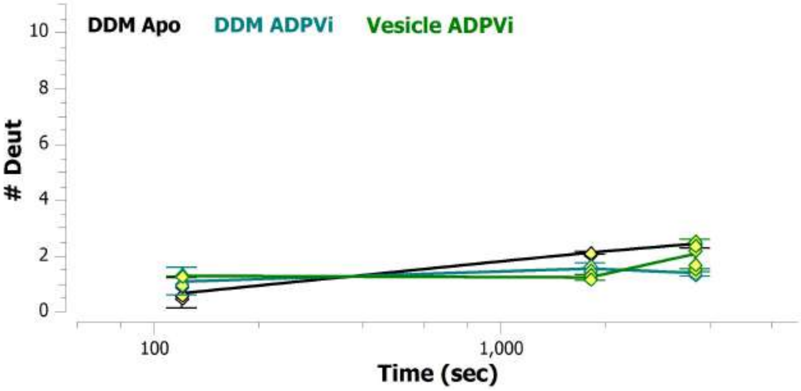

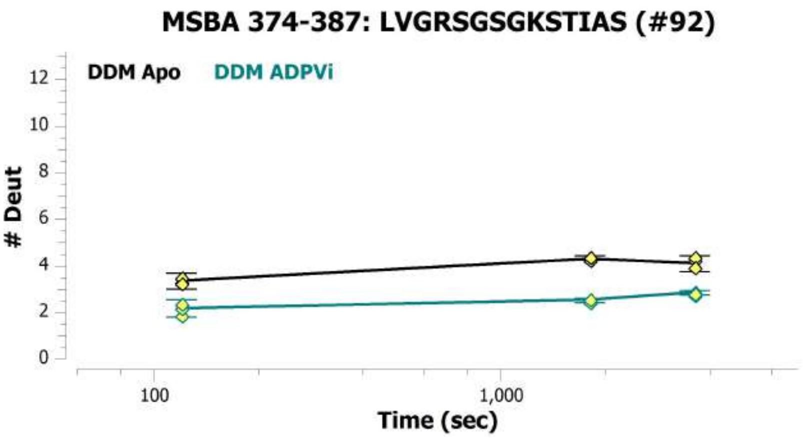
MSBA 374-387: LVGRSGSGKSTIAS (#92)
DDM Apo
DDM ADPVi
### Deut
0
2
4
6
8
10
12
100
1,000
Time (sec)

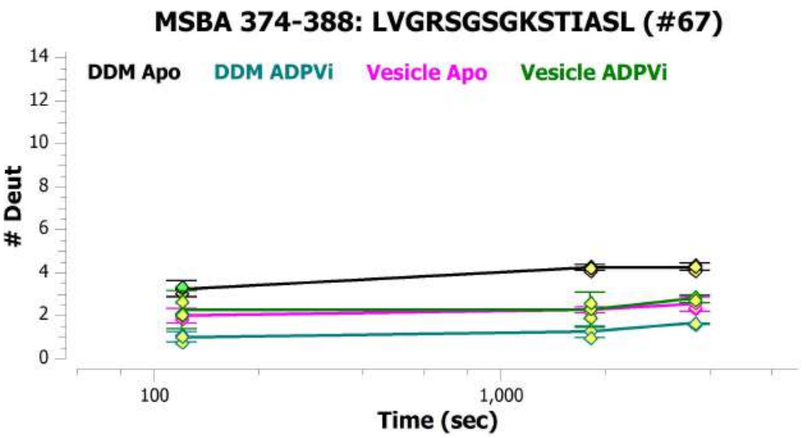
MSBA 374-388: LVGRSGSGKSTIASL (#67)
DDM Apo
DDM ADPVi
Vesicle Apo
Vesicle ADPVi
### Deut
0
2
4
6
8
10
12
14
100
1,000
Time (sec)

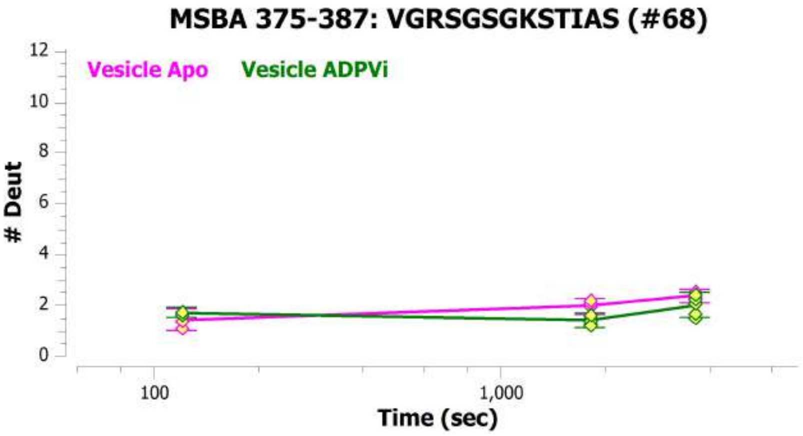
MSBA 375-387: VGRSGSGKSTIAS (#68)
Vesicle Apo
Vesicle ADPVi
### Deut
0
2
4
6
8
10
12
100
1,000
Time (sec)

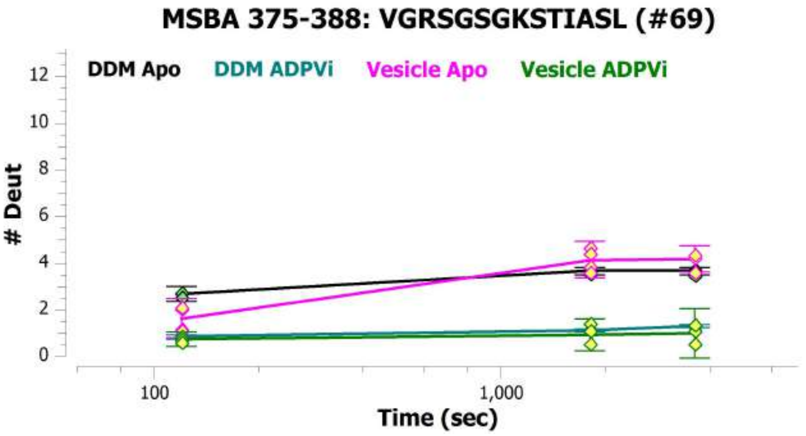
MSBA 375-388: VGRSGSGKSTIASL (#69)
DDM Apo
DDM ADPVi
Vesicle Apo
Vesicle ADPVi
### Deut
0
2
4
6
8
10
12
100
1,000
Time (sec)

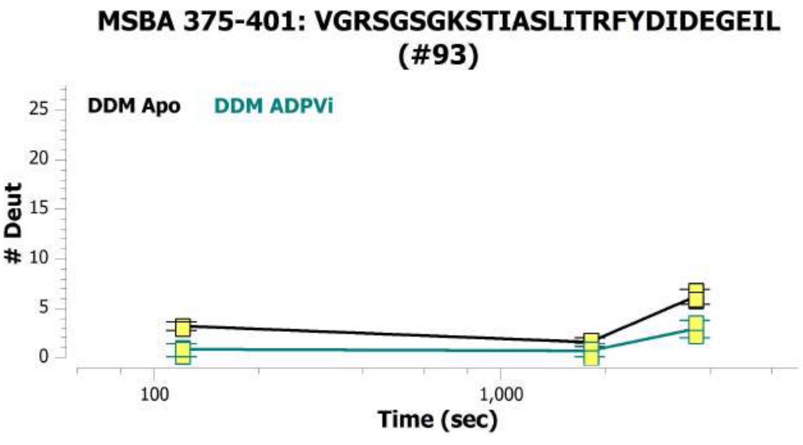
MSBA 375-401: VGRSGSGKSTIASLITRFYDIDEGEIL
(#93)
DDM Apo
DDM ADPVi
### Deut
0
5
10
15
20
25
100
1,000
Time (sec)

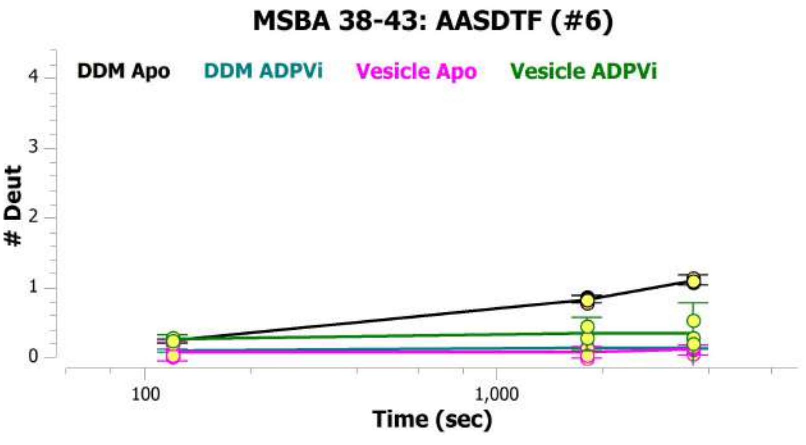
MSBA 38-43: AASDTF (#6)
DDM Apo
DDM ADPVi
Vesicle Apo
Vesicle ADPVi
### Deut
0
1
2
3
4
100
1,000
Time (sec)

**MSBA 389-401: ITRFYDIDEGEIL (#94)**

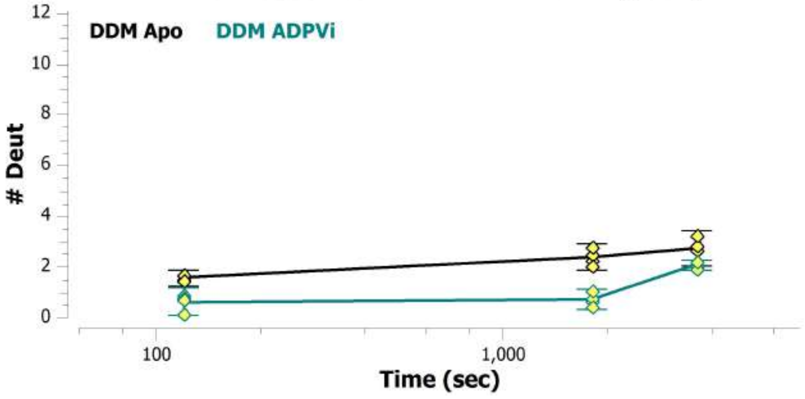


**MSBA 39-44: ASDTFM (#7)**

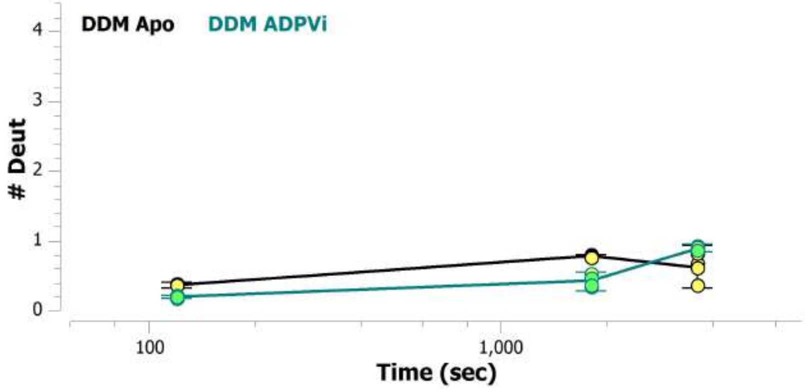


**MSBA 393-398: YDIDEG (#95)**

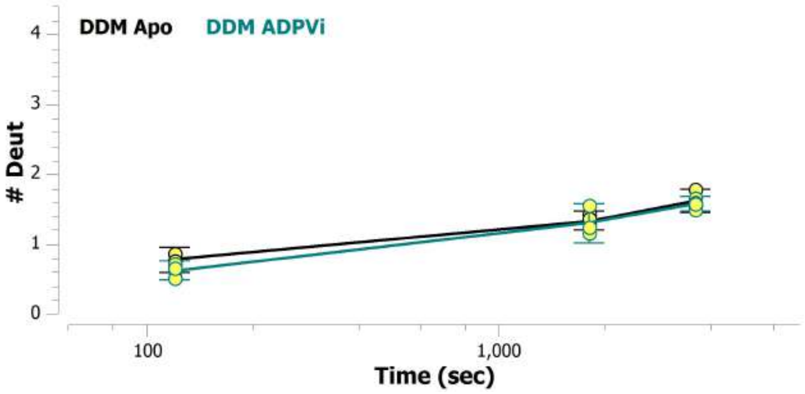


**MSBA 393-399: YDIDEGE (#96)**

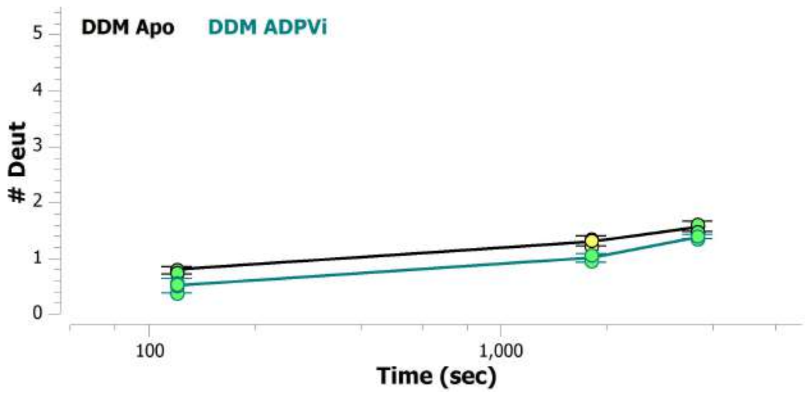


**MSBA 393-401: YDIDEGEIL (#70)**

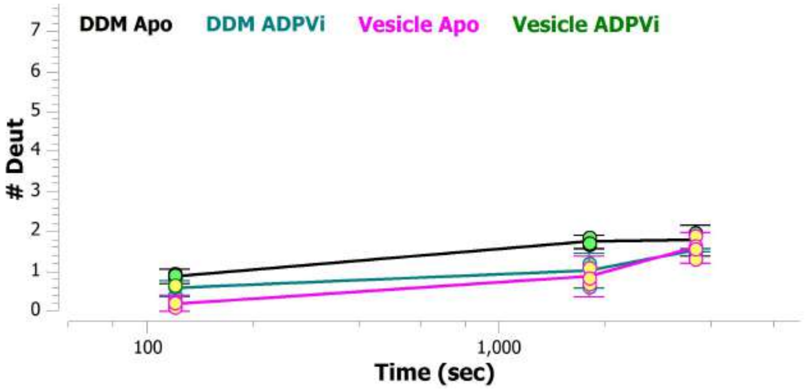


**MSBA 394-414: DIDEGEILMDGHDLREYTLAS (#71)**

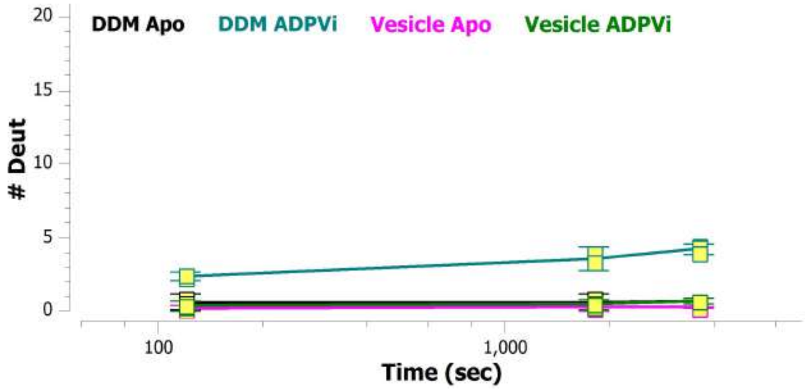

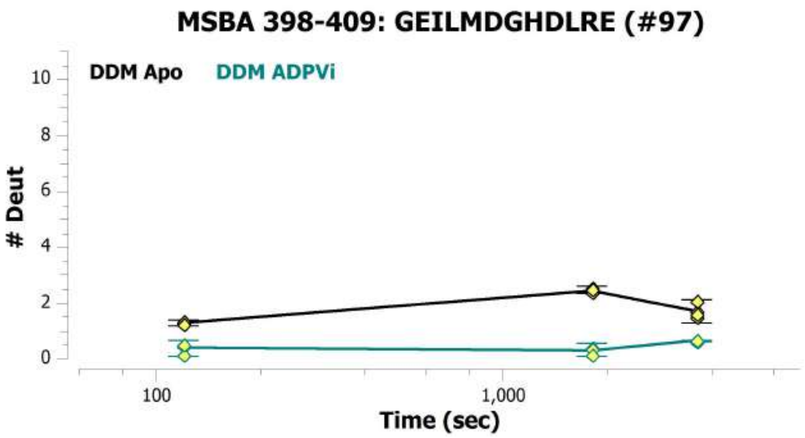
MSBA 398-409: GEILMDGHDLRE (#97)
DDM Apo
DDM ADPVi
### Deut
Time (sec)
0
2
4
6
8
10
100
1,000

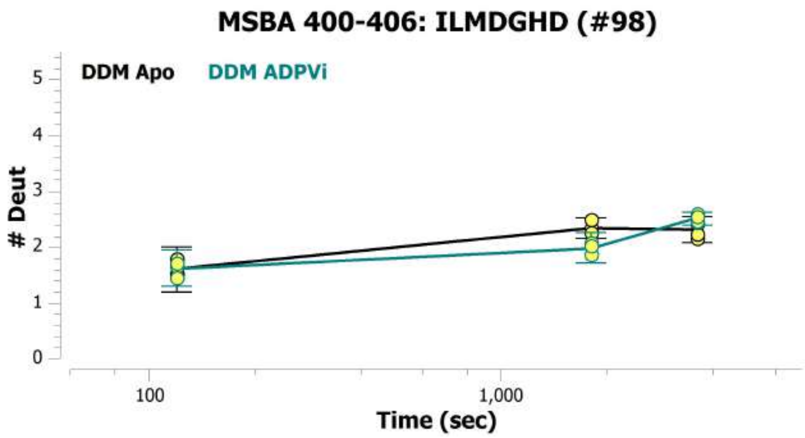
MSBA 400-406: ILMDGHD (#98)
DDM Apo
DDM ADPVi
### Deut
Time (sec)
0
1
2
3
4
5
100
1,000

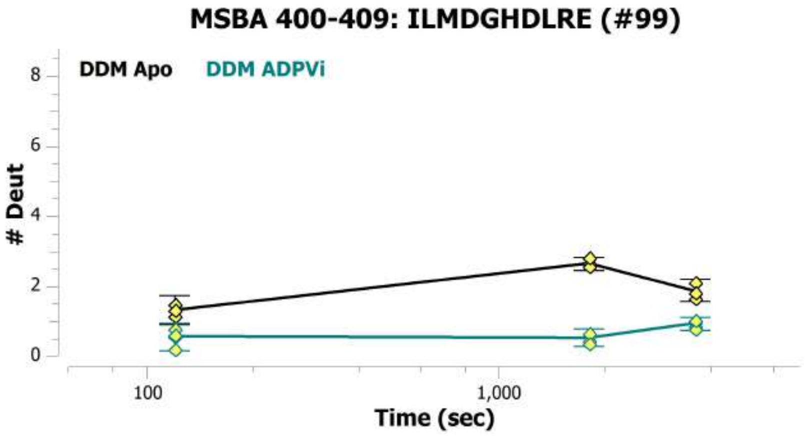
MSBA 400-409: ILMDGHDLRE (#99)
DDM Apo
DDM ADPVi
### Deut
Time (sec)
0
2
4
6
8
100
1,000

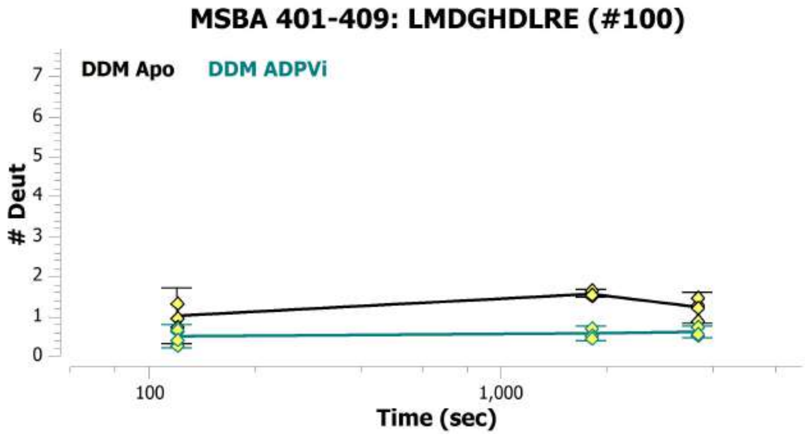
MSBA 401-409: LMDGHDLRE (#100)
DDM Apo
DDM ADPVi
### Deut
Time (sec)
0
1
2
3
4
5
6
7
100
1,000

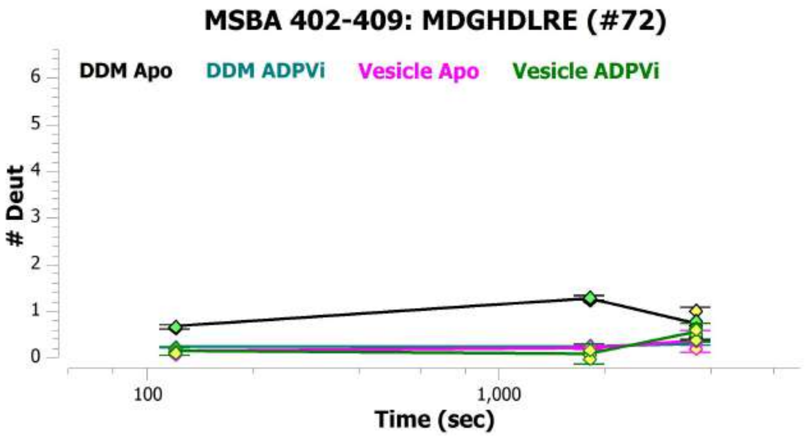
MSBA 402-409: MDGHDLRE (#72)
DDM Apo
DDM ADPVi
Vesicle Apo
Vesicle ADPVi
### Deut
Time (sec)
0
1
2
3
4
5
6
100
1,000

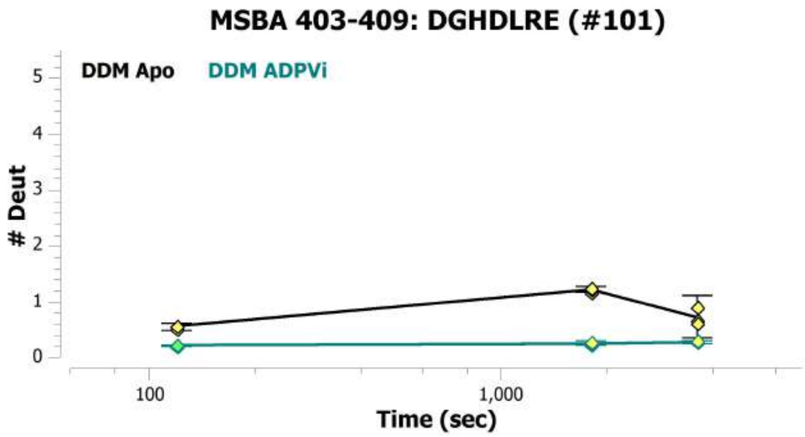
MSBA 403-409: DGHDLRE (#101)
DDM Apo
DDM ADPVi
### Deut
Time (sec)
0
1
2
3
4
5
100
1,000

**MSBA 410-421: YTLASLRNQVAL (#73)**

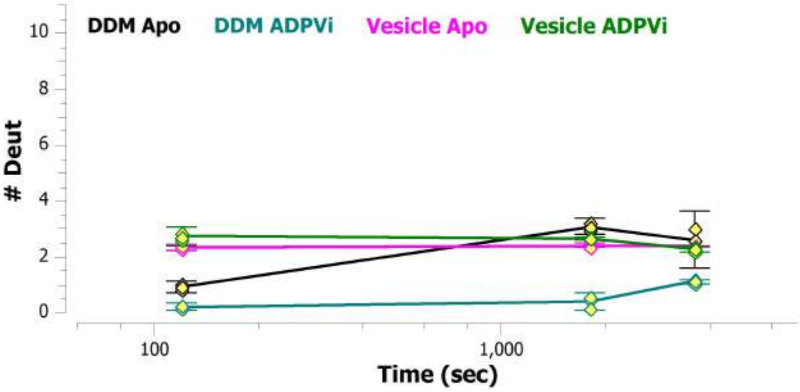


**MSBA 413-420: ASLRNQVA (#102)**

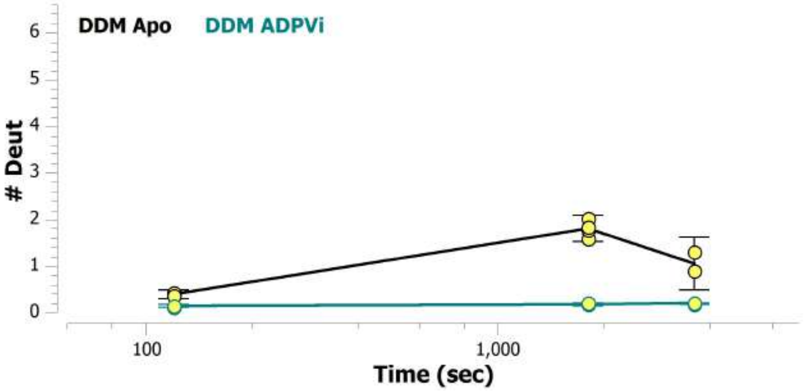


**MSBA 413-421: ASLRNQVAL (#74)**

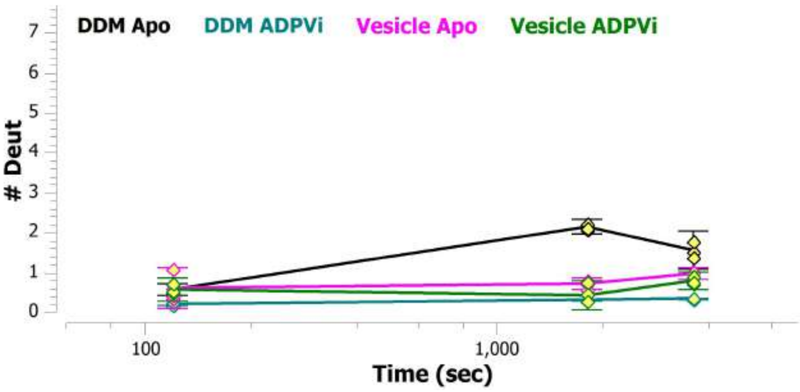


**MSBA 416-421: RNQVAL (#75)**

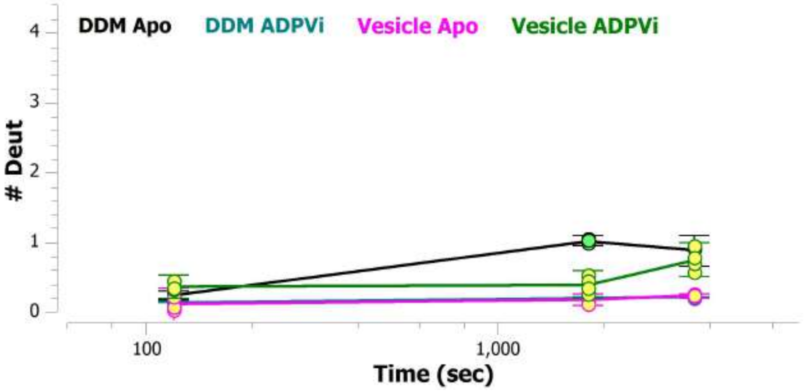


**MSBA 421-428: LVSQNVHL (#103)**

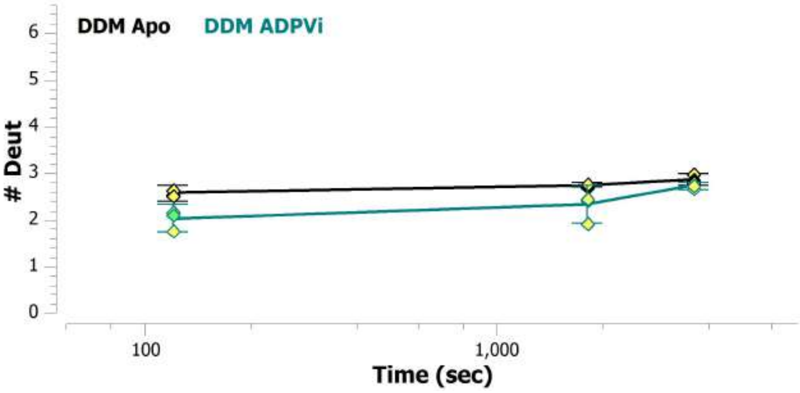


**MSBA 422-428: VSQNVHL (#76)**

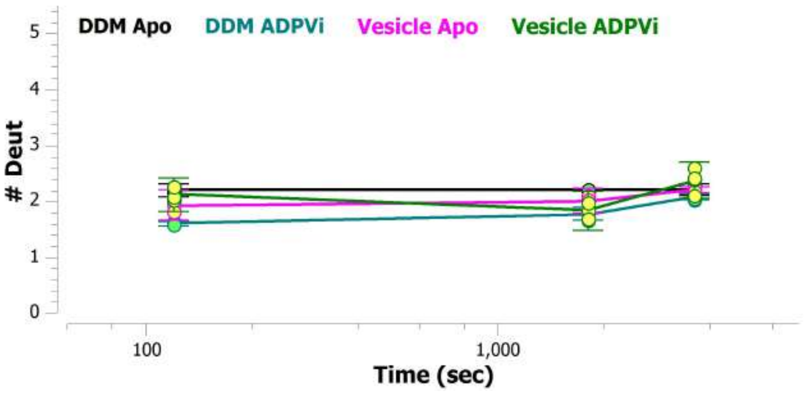

MSBA 422-436: VSQNVHLFNDTVANN (#104)

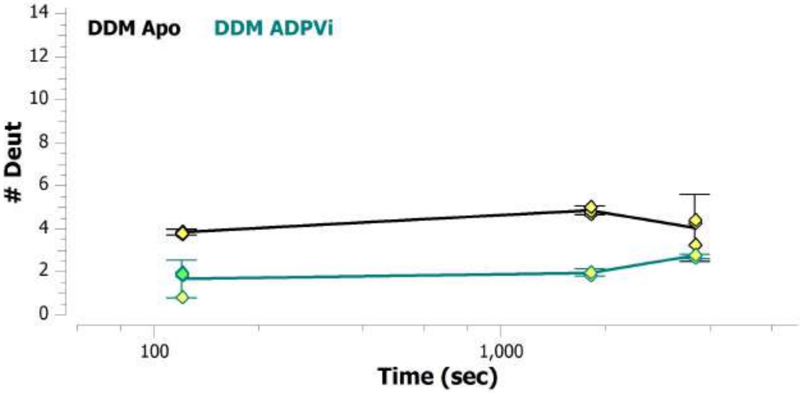


MSBA 422-438: VSQNVHLFNDTVANNIA (#105)

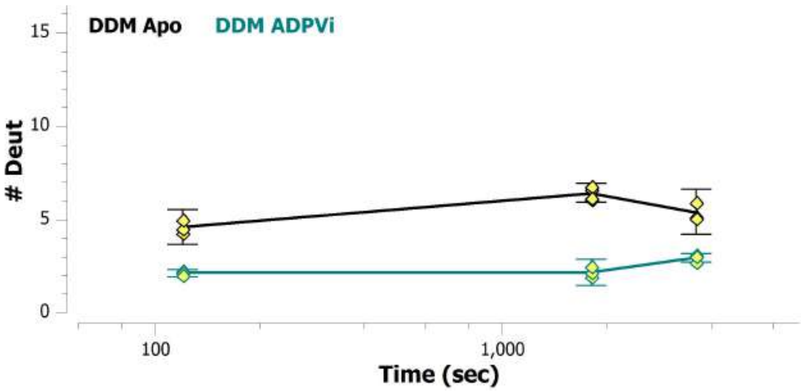


MSBA 423-428: SQNVHL (#77)

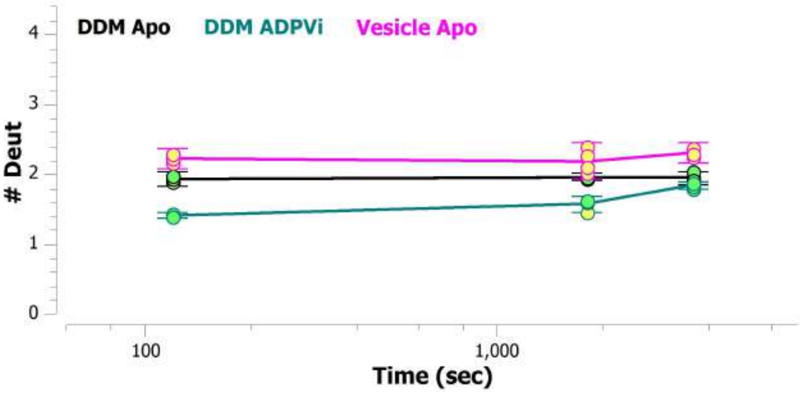


MSBA 429-436: FNDTVANN (#78)

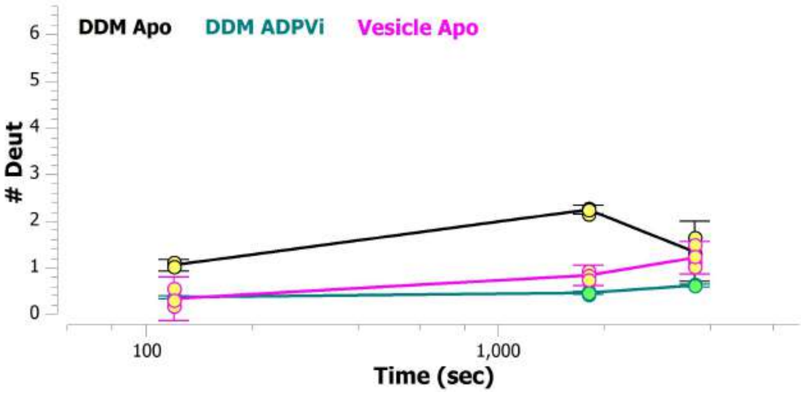


MSBA 429-438: FNDTVANNIA (#106)

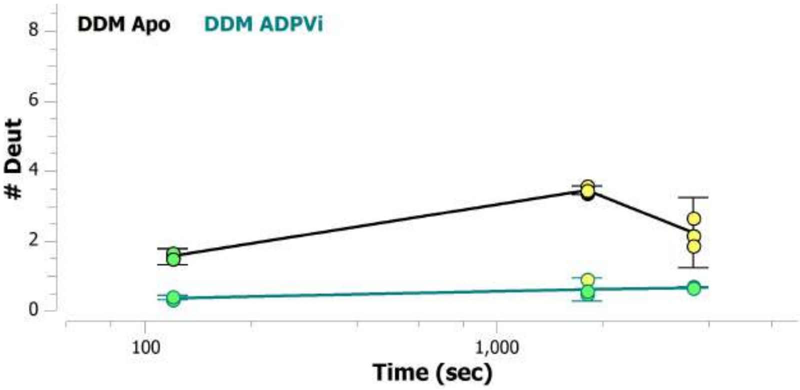


MSBA 429-439: FNDTVANNIAY (#107)

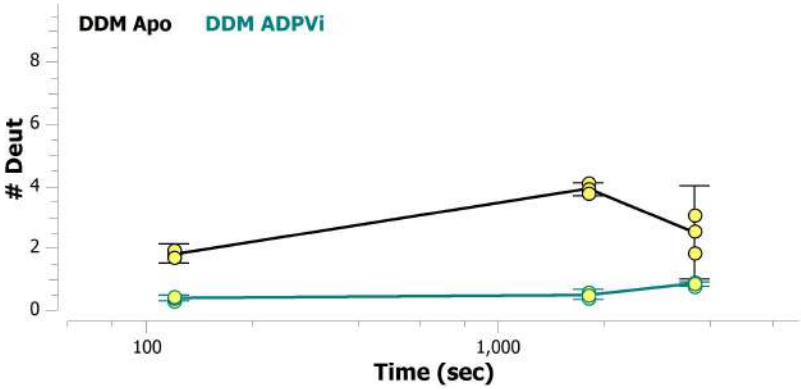

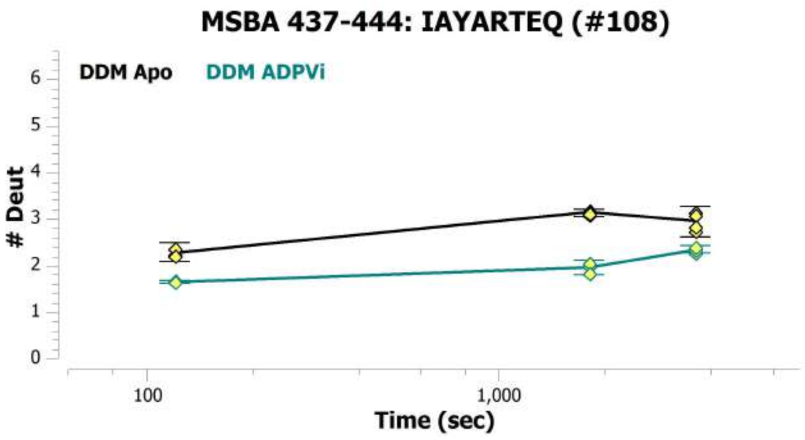
MSBA 437-444: IAYARTEQ (#108)
DDM Apo
DDM ADPVi
### Deut
0
1
2
3
4
5
6
100
1,000
Time (sec)

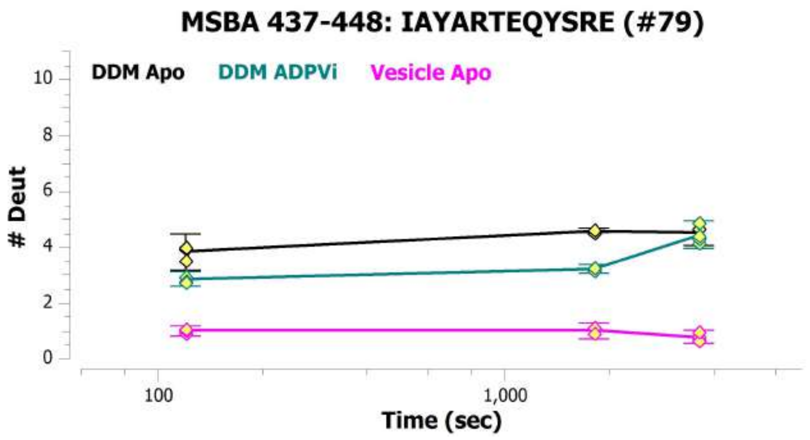
MSBA 437-448: IAYARTEQYSRE (#79)
DDM Apo
DDM ADPVi
Vesicle Apo
### Deut
0
2
4
6
8
10
100
1,000
Time (sec)

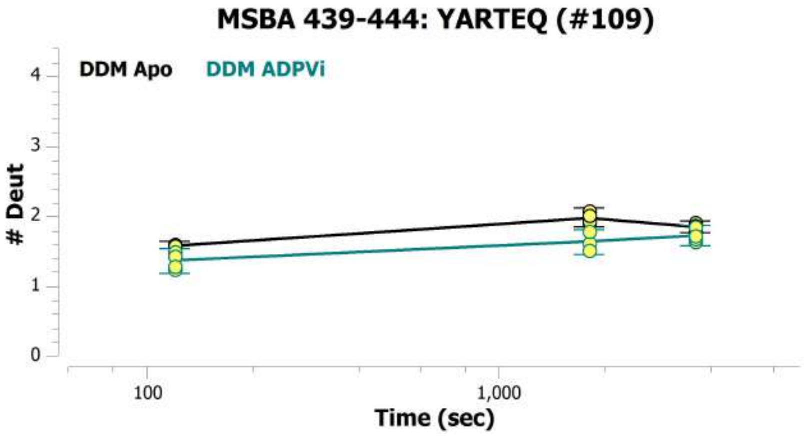
MSBA 439-444: YARTEQ (#109)
DDM Apo
DDM ADPVi
### Deut
0
1
2
3
4
100
1,000
Time (sec)

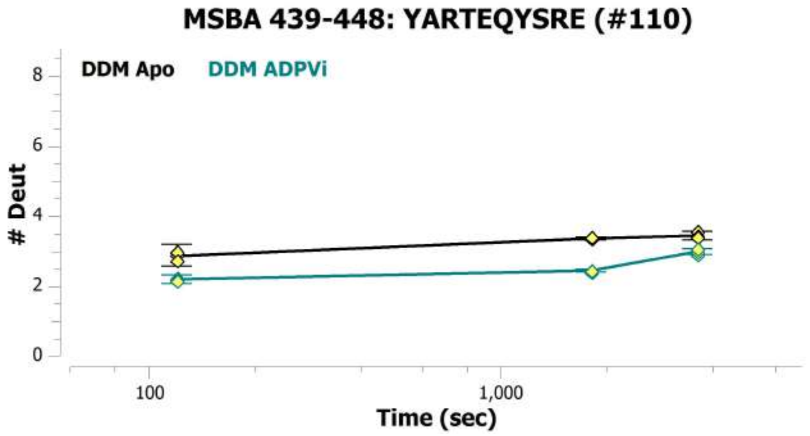
MSBA 439-448: YARTEQYSRE (#110)
DDM Apo
DDM ADPVi
### Deut
0
2
4
6
8
100
1,000
Time (sec)

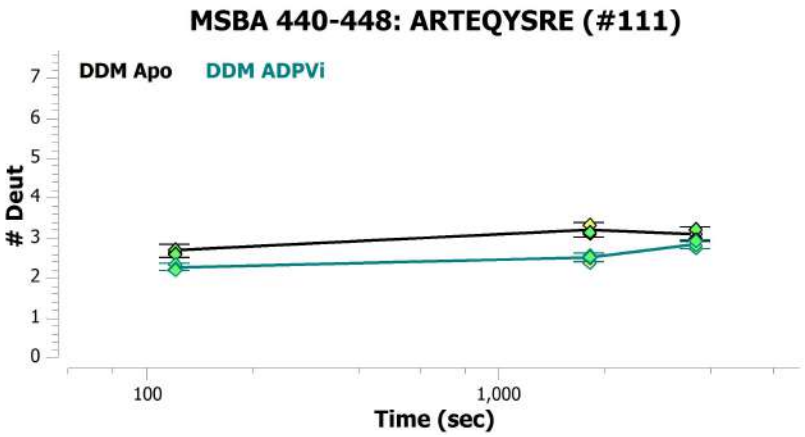
MSBA 440-448: ARTEQYSRE (#111)
DDM Apo
DDM ADPVi
### Deut
0
1
2
3
4
5
6
7
100
1,000
Time (sec)

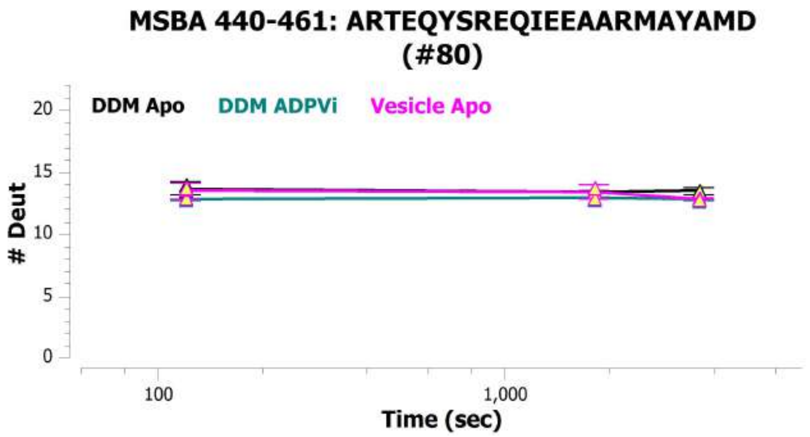
MSBA 440-461: ARTEQYSREQIEEAARMAYAMD (#80)
DDM Apo
DDM ADPVi
Vesicle Apo
### Deut
0
5
10
15
20
100
1,000
Time (sec)

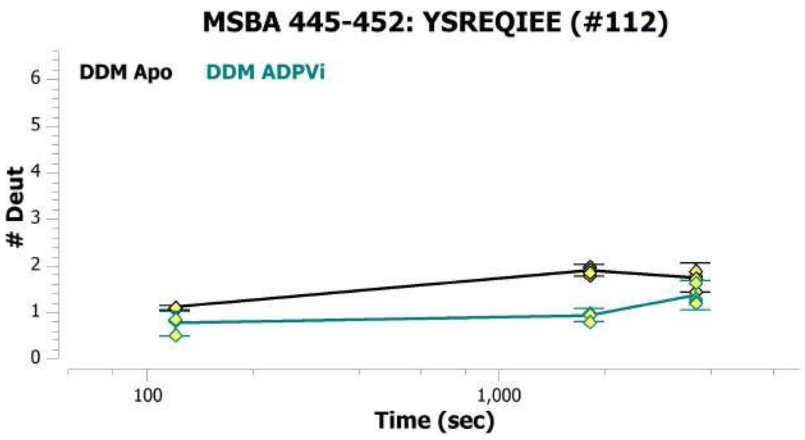
MSBA 445-452: YSREQIEE (#112)
DDM Apo
DDM ADPVi
### Deut
Time (sec)

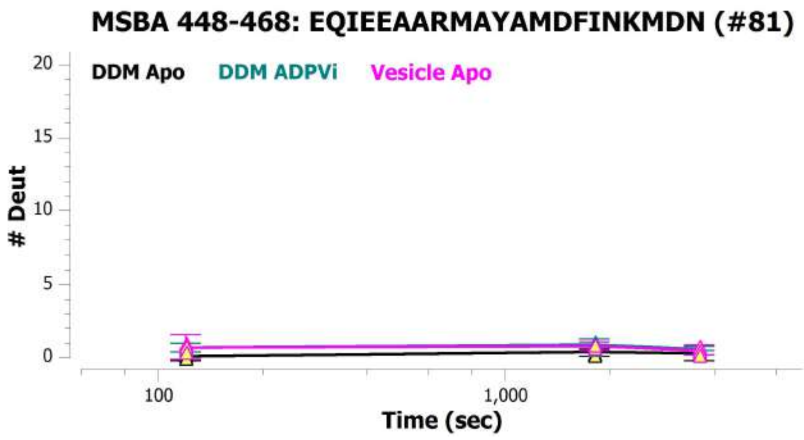
MSBA 448-468: EQIEEAARMAYAMDFINKMDN (#81)
DDM Apo
DDM ADPVi
Vesicle Apo
### Deut
Time (sec)

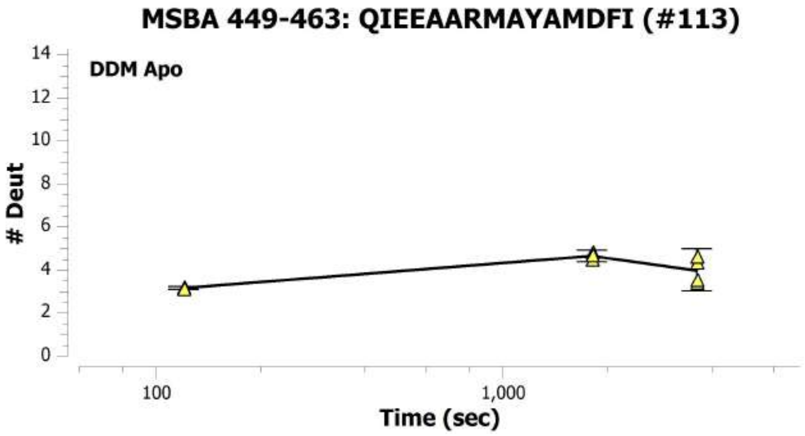
MSBA 449-463: QIEEAARMAYAMDFI (#113)
DDM Apo
### Deut
Time (sec)

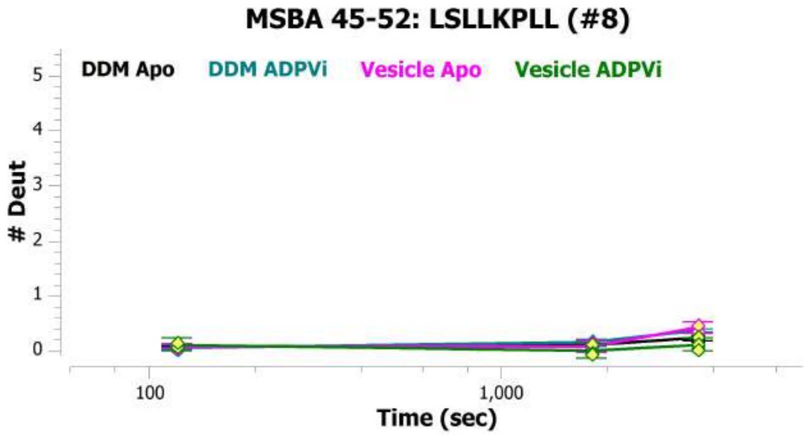
MSBA 45-52: LSLLKPLL (#8)
DDM Apo
DDM ADPVi
Vesicle Apo
Vesicle ADPVi
### Deut
Time (sec)

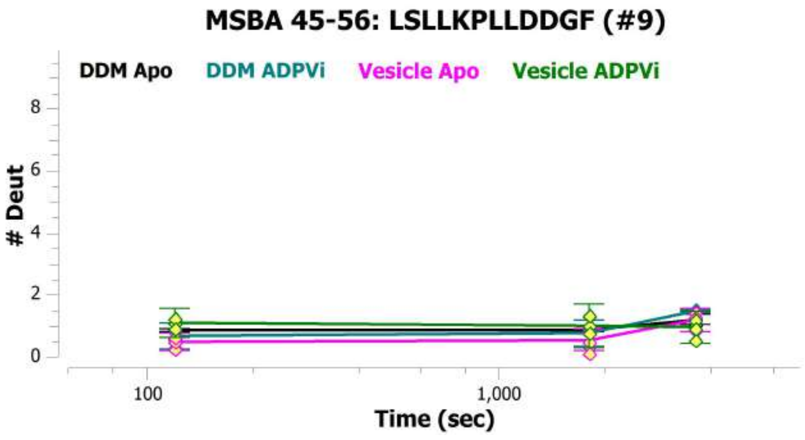
MSBA 45-56: LSLLKPLLDDGF (#9)
DDM Apo
DDM ADPVi
Vesicle Apo
Vesicle ADPVi
### Deut
Time (sec)

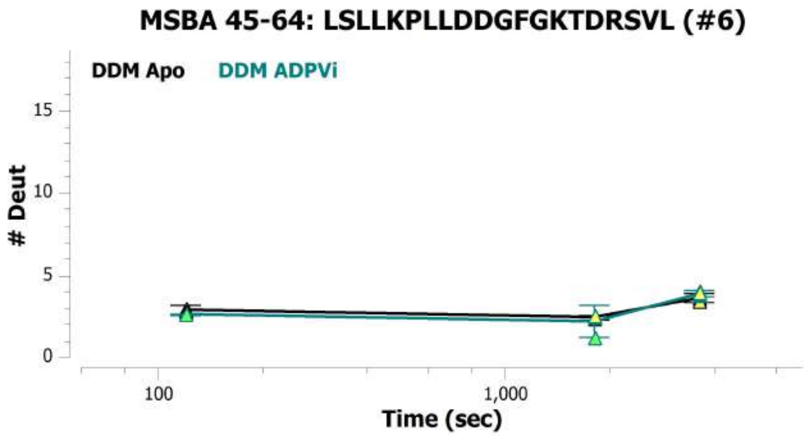
MSBA 45-64: LSLLKPLLDDGFGKTDRSVL (#6)
DDM Apo
DDM ADPVi
### Deut
Time (sec)

**MSBA 452-462: EAARMAYAMDF (#82)**

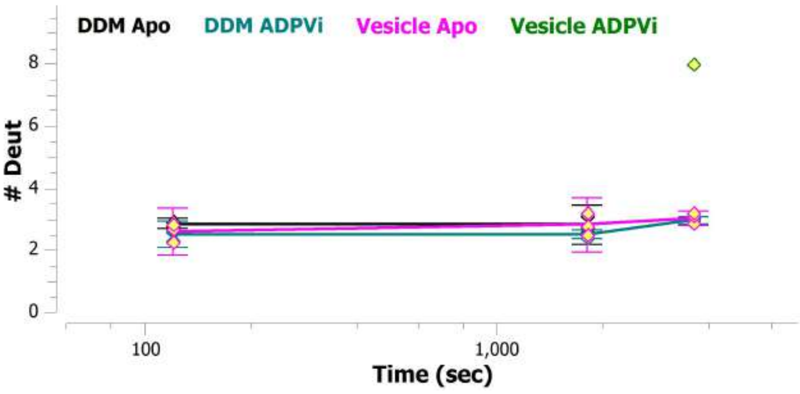


**MSBA 453-461: AARMAYAMD (#114)**

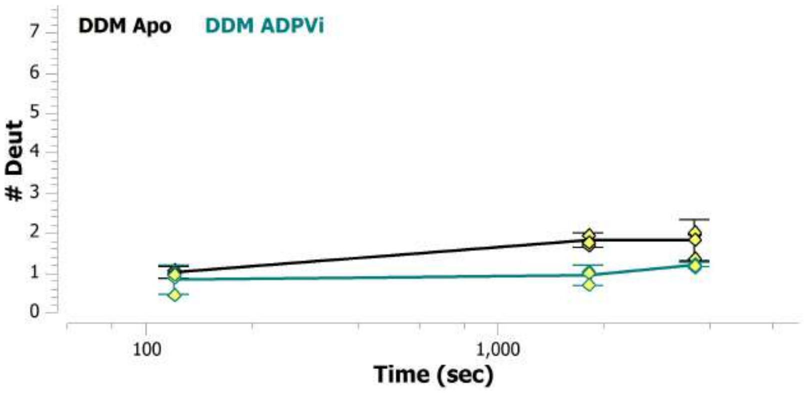


**MSBA 453-462: AARMAYAMDF (#115)**

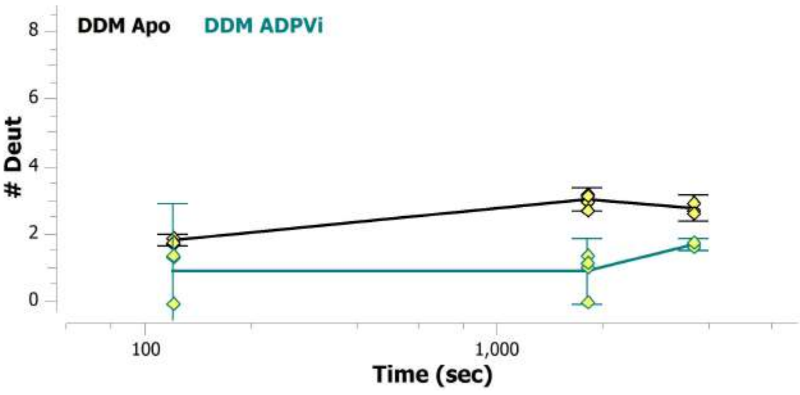


**MSBA 454-461: ARMAYAMD (#116)**

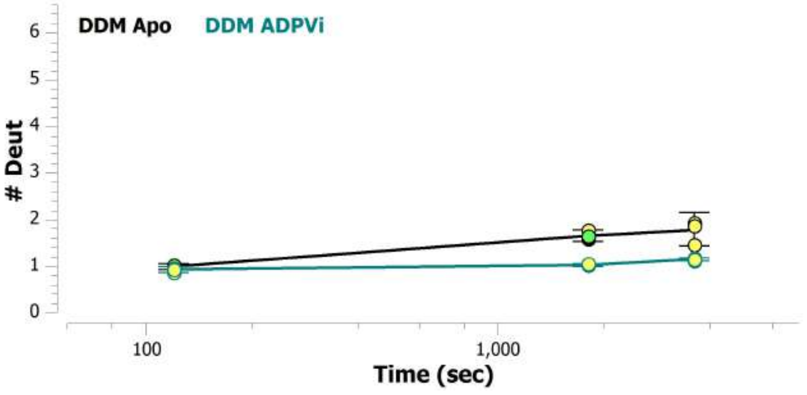


**MSBA 457-462: AYAMDF (#83)**

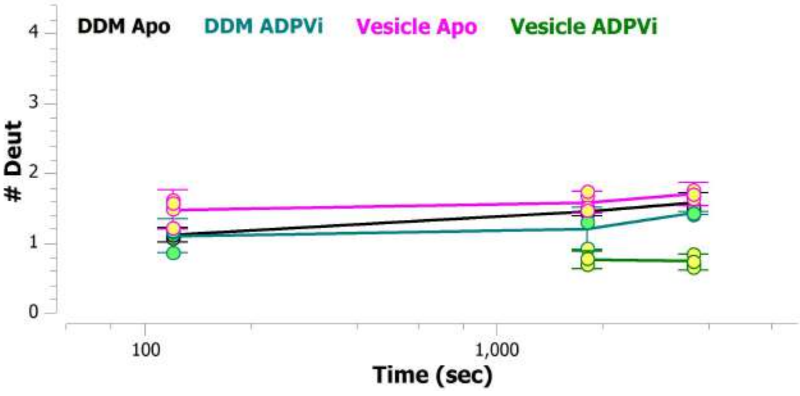


**MSBA 461-471: DFINKMDNGLD (#117)**

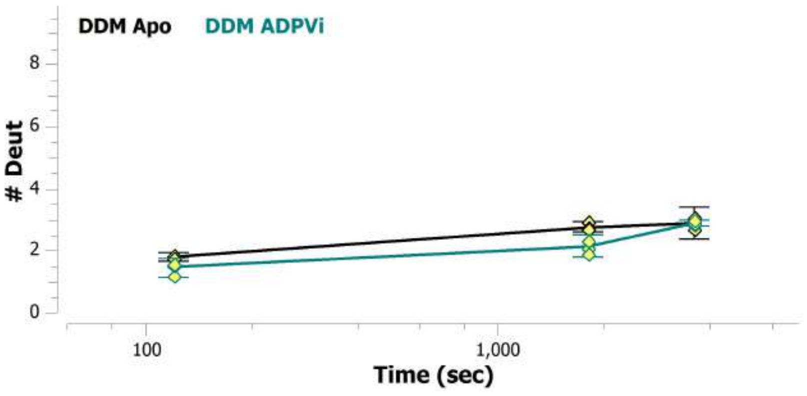

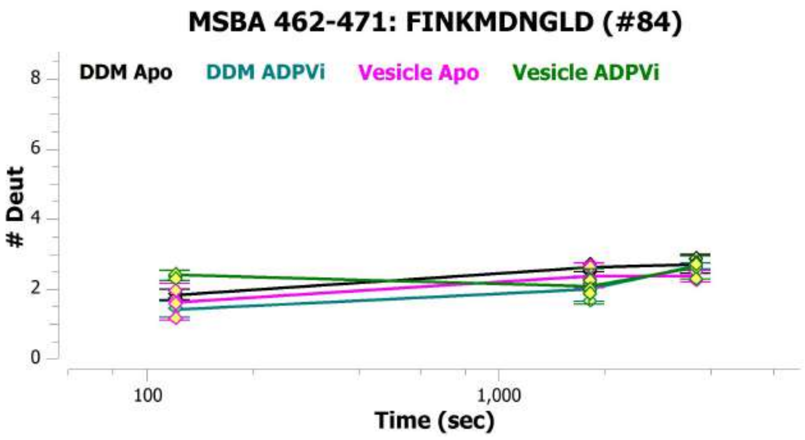

MSBA 462-471: FINKMDNGLD (#84)
DDM Apo
DDM ADPVi
Vesicle Apo
Vesicle ADPVi
### Deut
Time (sec)


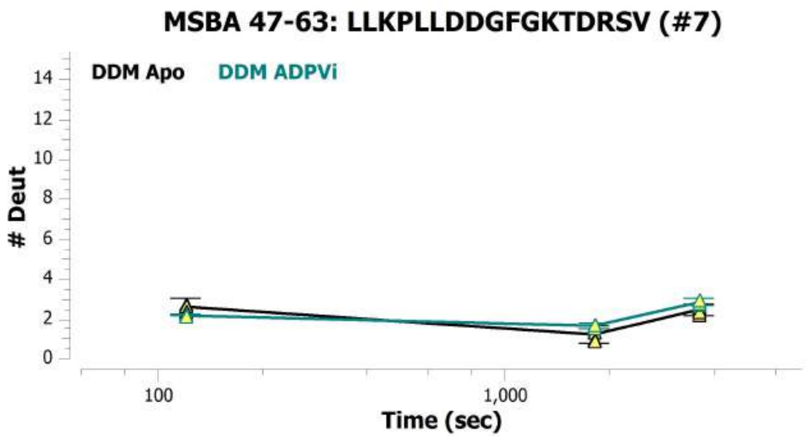

MSBA 47-63: LLKPLLDDGFGKTDRSV (#7)
DDM Apo
DDM ADPVi
### Deut
Time (sec)


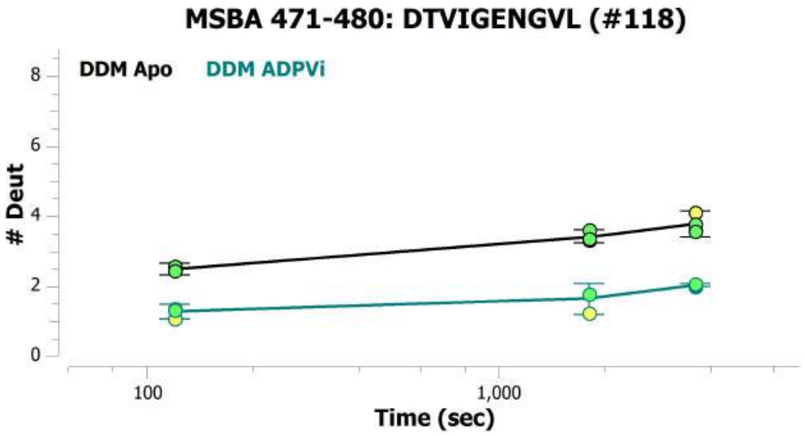

MSBA 471-480: DTVIGENGVL (#118)
DDM Apo
DDM ADPVi
### Deut
Time (sec)


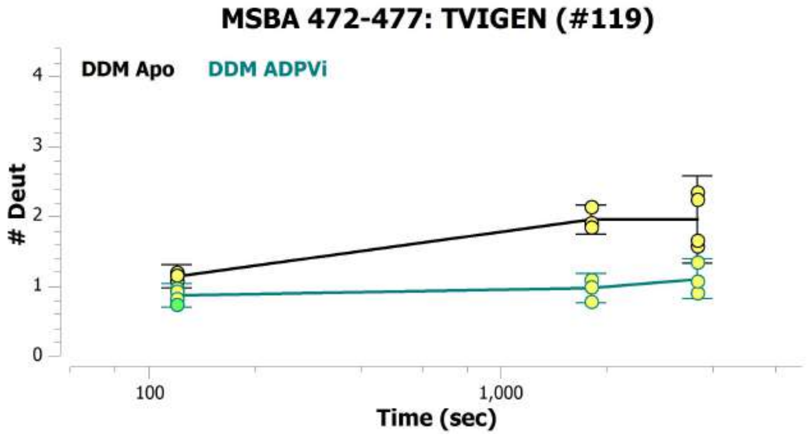

MSBA 472-477: TVIGEN (#119)
DDM Apo
DDM ADPVi
### Deut
Time (sec)


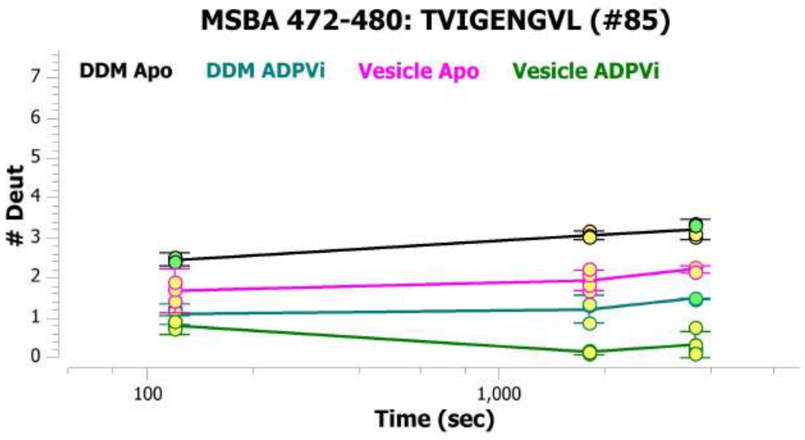

MSBA 472-480: TVIGENGVL (#85)
DDM Apo
DDM ADPVi
Vesicle Apo
Vesicle ADPVi
### Deut
Time (sec)


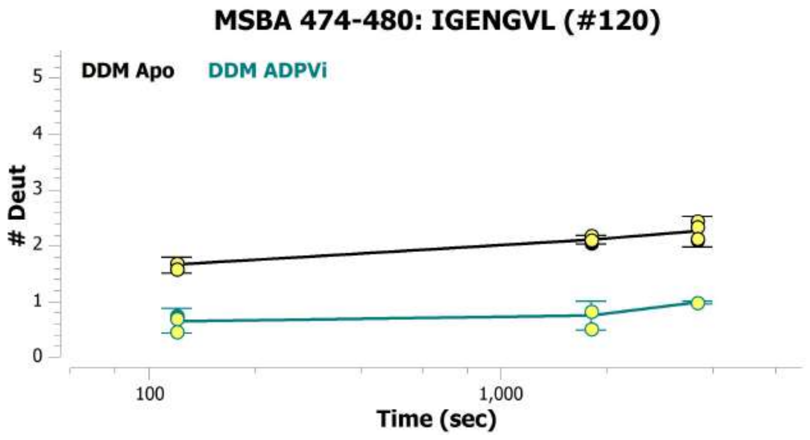

MSBA 474-480: IGENGVL (#120)
DDM Apo
DDM ADPVi
### Deut
Time (sec)

##### MSBA 475-480: GENGVL (#121)

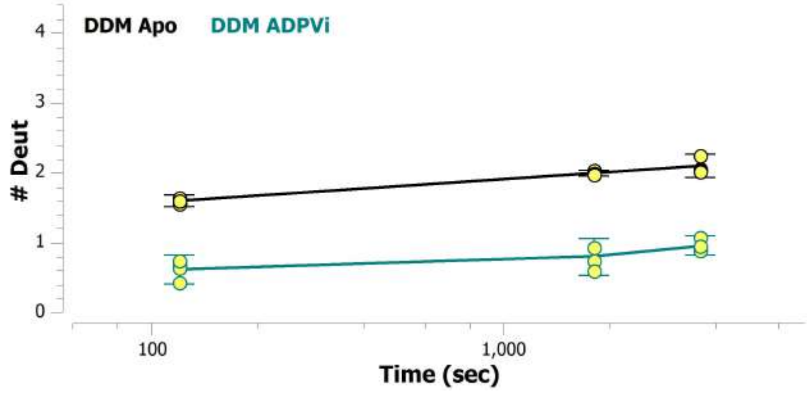


##### MSBA 481-491: LSGGQRQRIAI (#86)

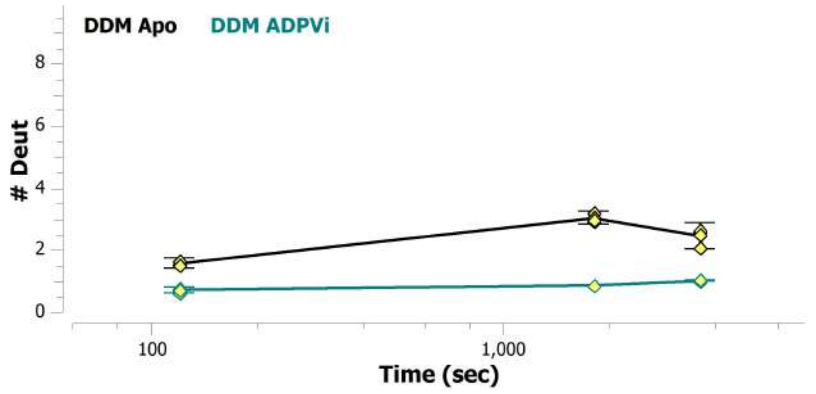


##### MSBA 481-492: LSGGQRQRIAIA (#87)

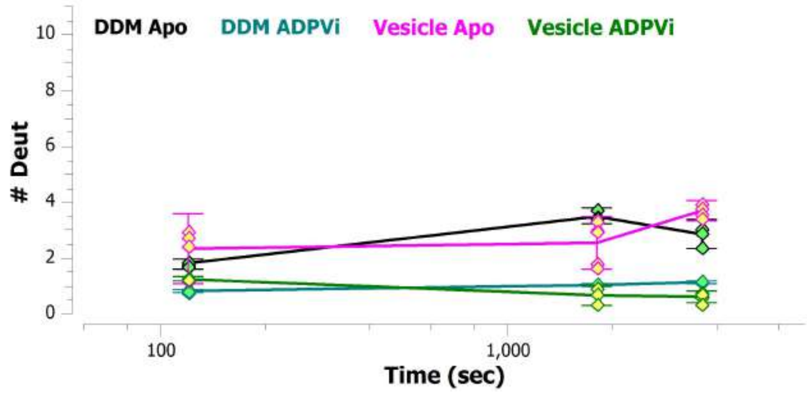


##### MSBA 481-496: LSGGQRQRIAIARALL (#122)

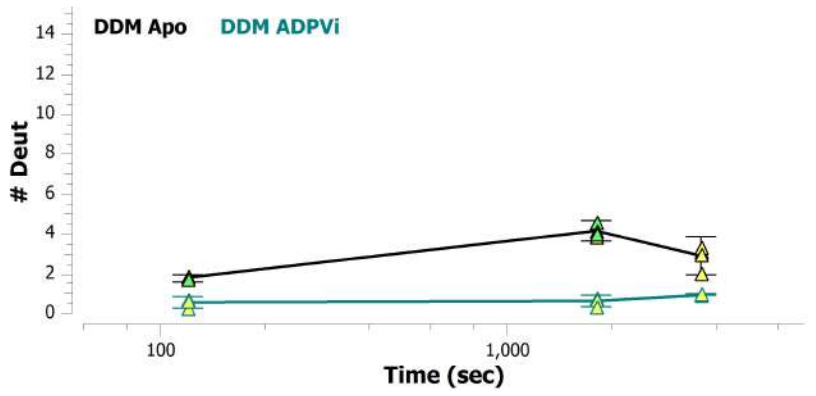


##### MSBA 481-502: LSGGQRQRIAIARALLRDSPIL (#88)

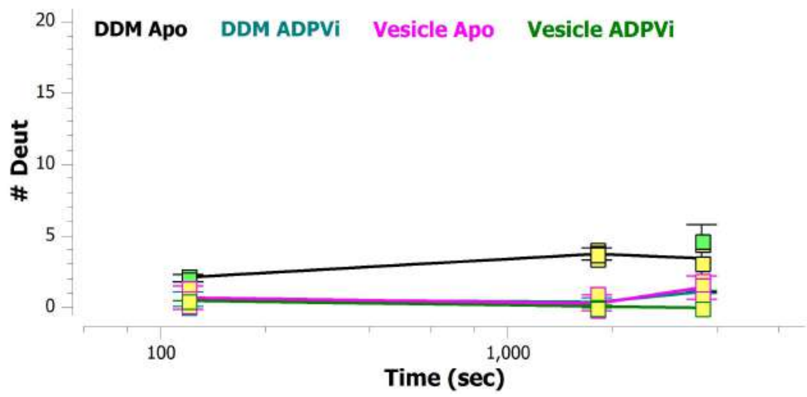


##### MSBA 482-492: SGGQRQRIAIA (#89)

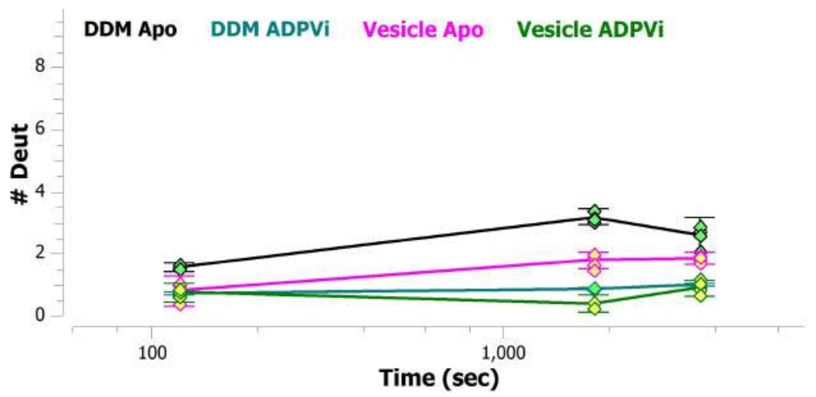

MSBA 482-494: SGGQRQRIAIARA (#123)

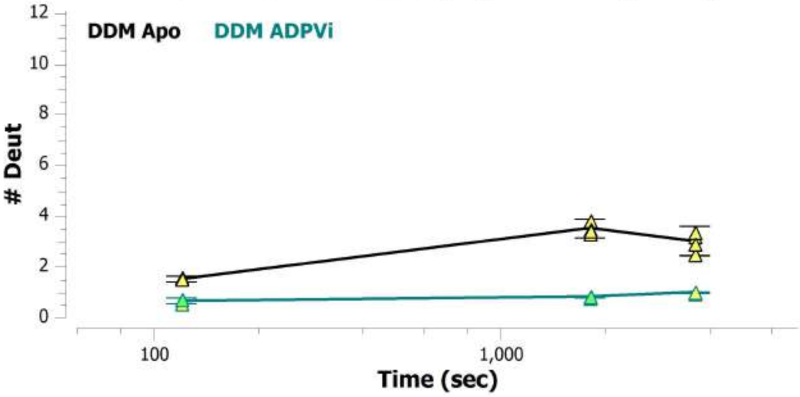


MSBA 482-495: SGGQRQRIAIARAL (#124)

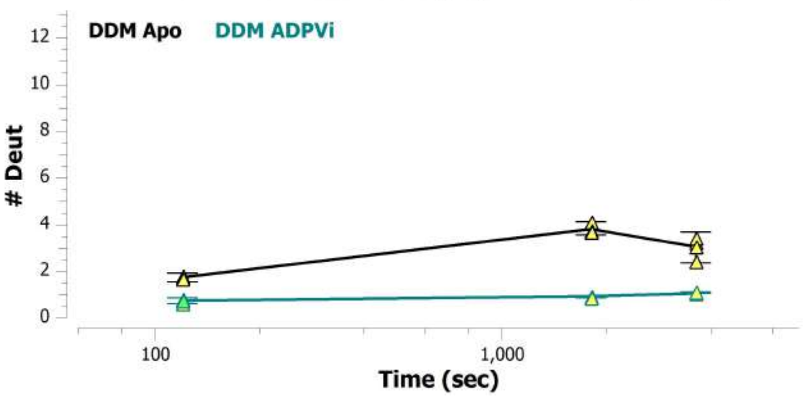


MSBA 482-496: SGGQRQRIAIARALL (#125)

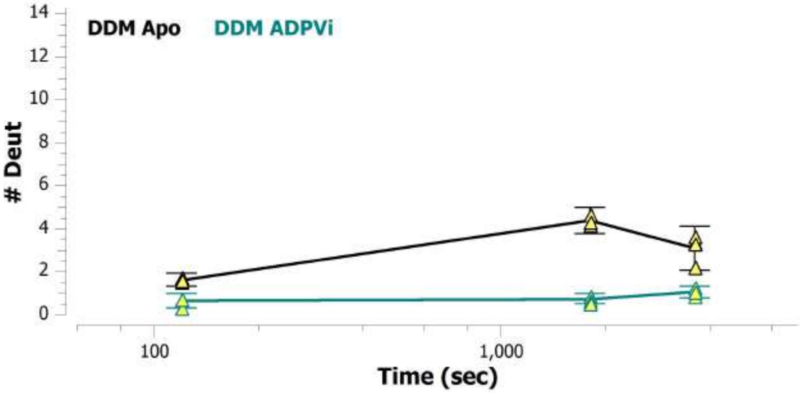


MSBA 483-492: GGQRQRIAIA (#90)

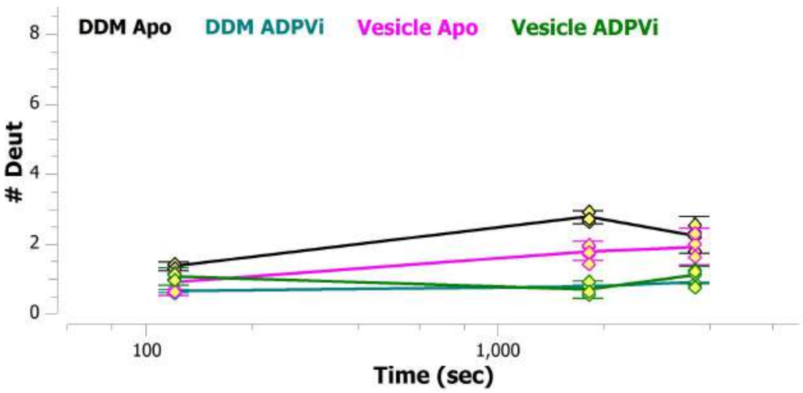


MSBA 483-494: GGQRQRIAIARA (#126)

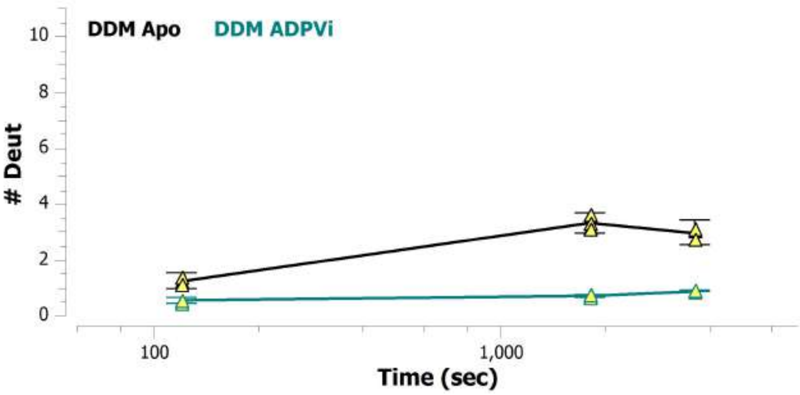


MSBA 483-496: GGQRQRIAIARALL (#127)

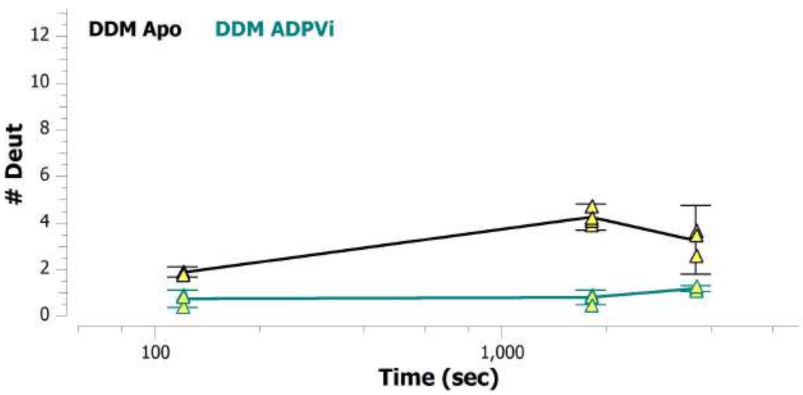

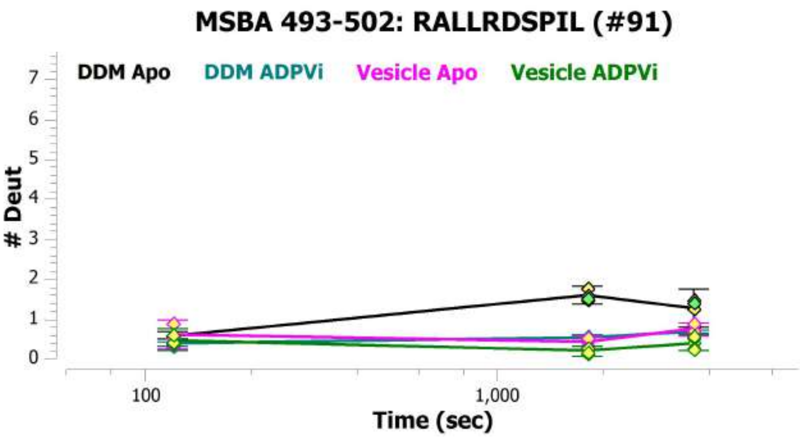
MSBA 493-502: RALLRDSPIL (#91)
DDM Apo
DDM ADPVi
Vesicle Apo
Vesicle ADPVi
### Deut
0
1
2
3
4
5
6
7
100
1,000
Time (sec)

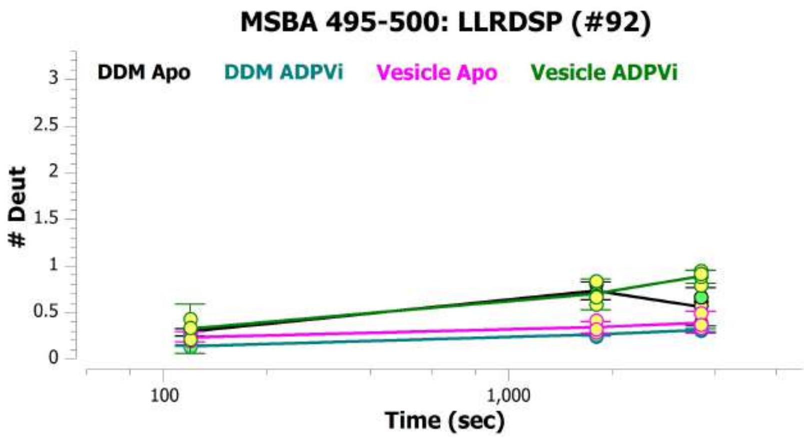
MSBA 495-500: LLRDSP (#92)
DDM Apo
DDM ADPVi
Vesicle Apo
Vesicle ADPVi
### Deut
0
0.5
1
1.5
2
2.5
3
100
1,000
Time (sec)

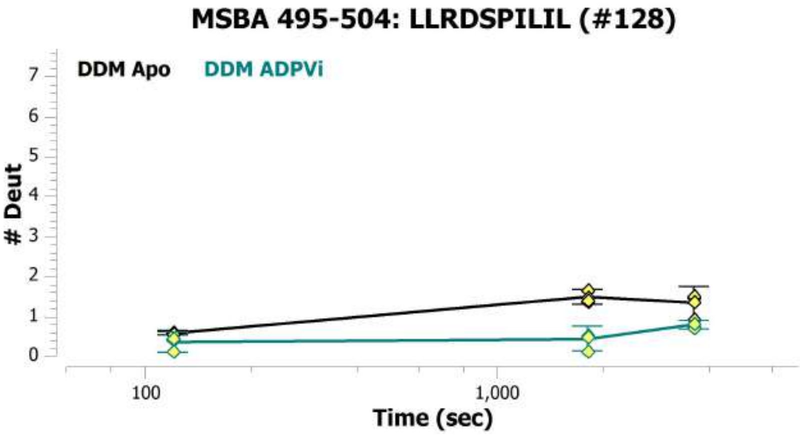
MSBA 495-504: LLRDSPILIL (#128)
DDM Apo
DDM ADPVi
### Deut
0
1
2
3
4
5
6
7
100
1,000
Time (sec)

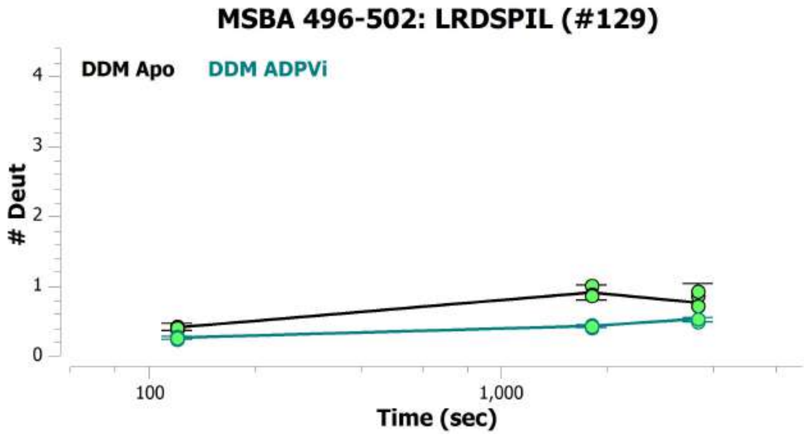
MSBA 496-502: LRDSPIL (#129)
DDM Apo
DDM ADPVi
### Deut
0
1
2
3
4
100
1,000
Time (sec)

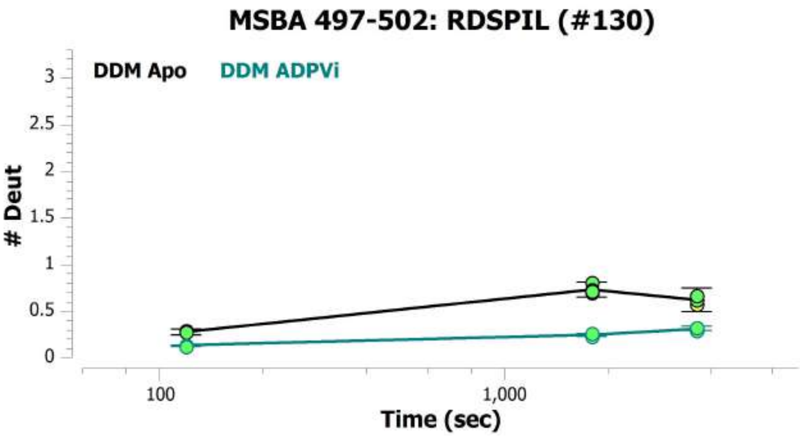
MSBA 497-502: RDSPIL (#130)
DDM Apo
DDM ADPVi
### Deut
0
0.5
1
1.5
2
2.5
3
100
1,000
Time (sec)

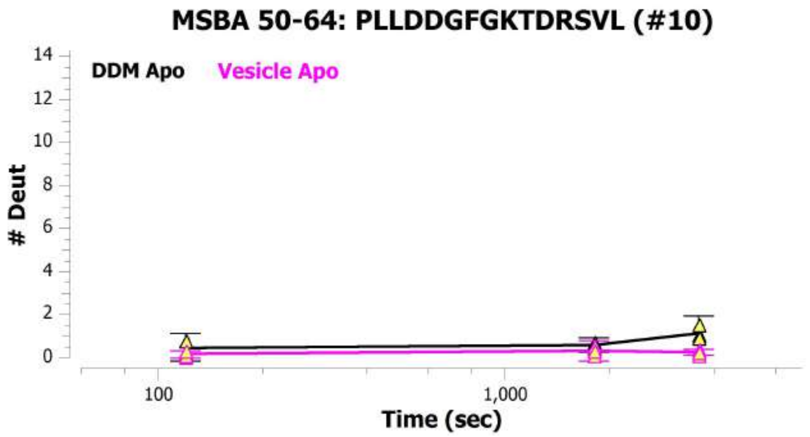
MSBA 50-64: PLLDDGFGKTDRSVL (#10)
DDM Apo
Vesicle Apo
### Deut
0
2
4
6
8
10
12
14
100
1,000
Time (sec)

MSBA 503-508: ILDEAT (#131)

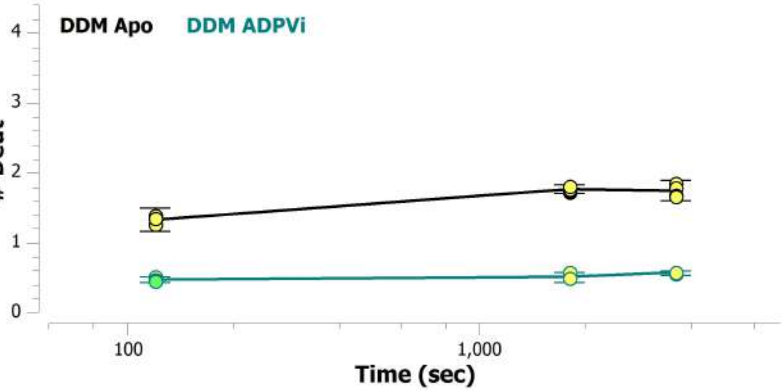


MSBA 503-511: ILDEATSAL (#93)

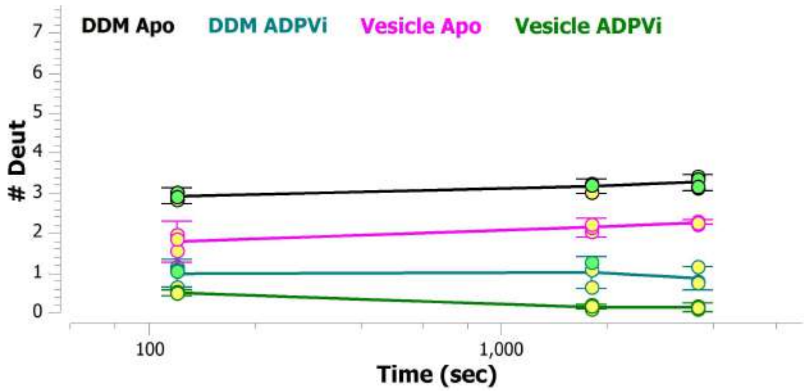


MSBA 505-511: DEATSAL (#132)

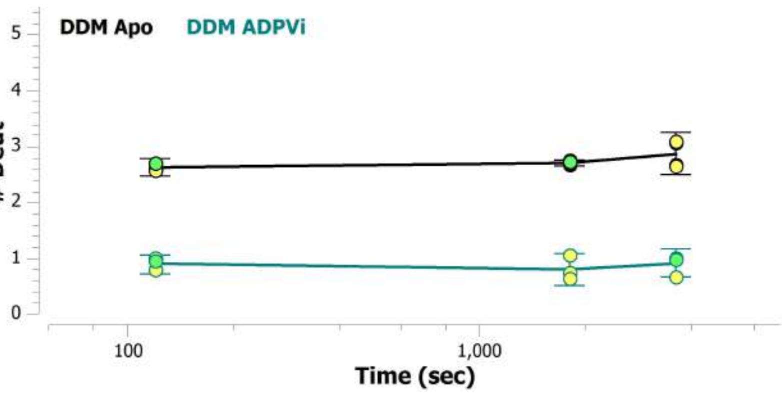


MSBA 506-511: EATSAL (#133)

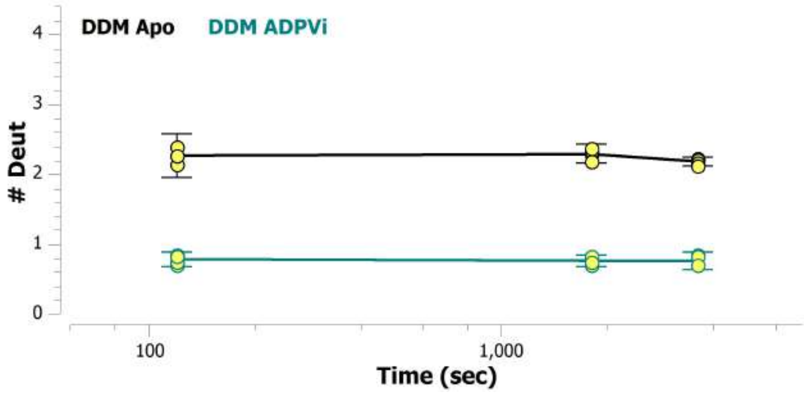


MSBA 51-56: LLDDGF (#11)

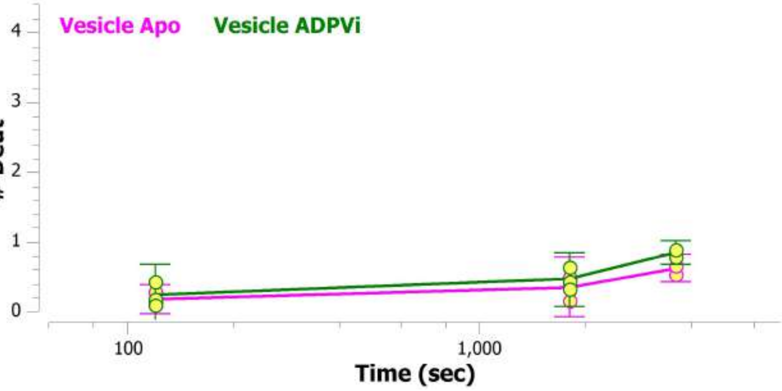


MSBA 511-516: LDTESE (#94)

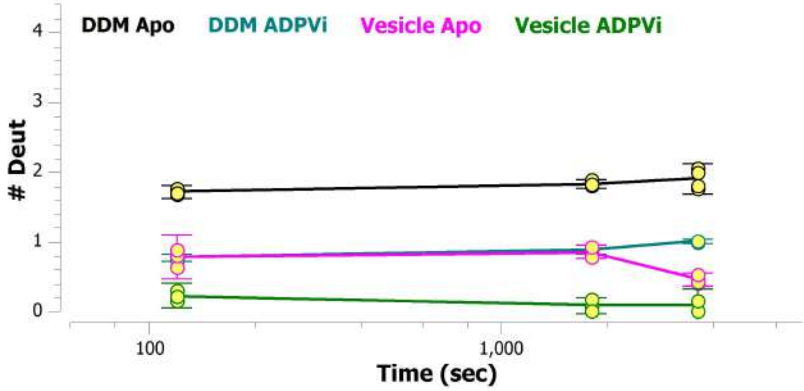

MSBA 512-520: DTESERAIQ (#134)

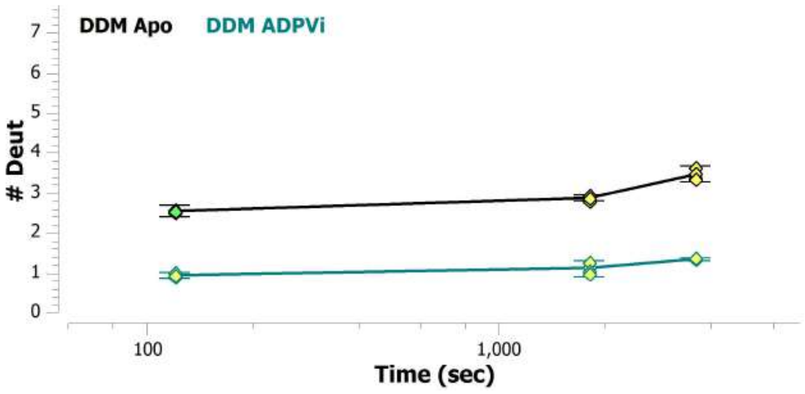


MSBA 512-522: DTESERAIQAA (#135)

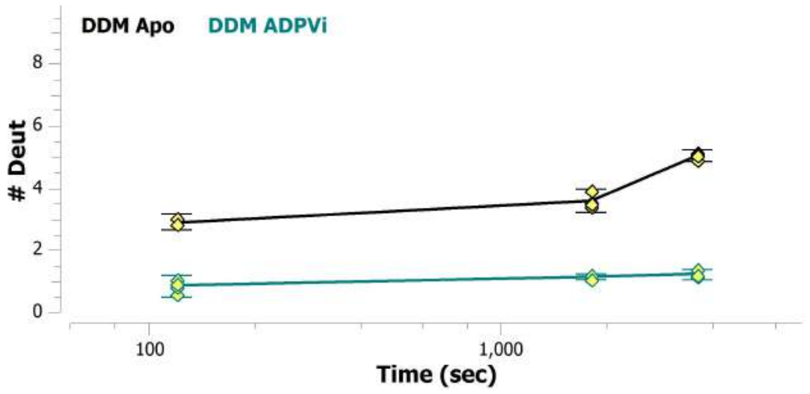


MSBA 512-523: DTESERAIQAAL (#136)

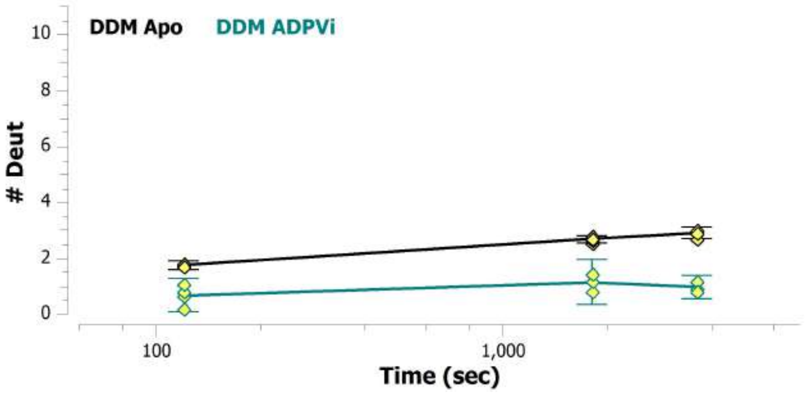


MSBA 517-523: RAIQAAL (#137)

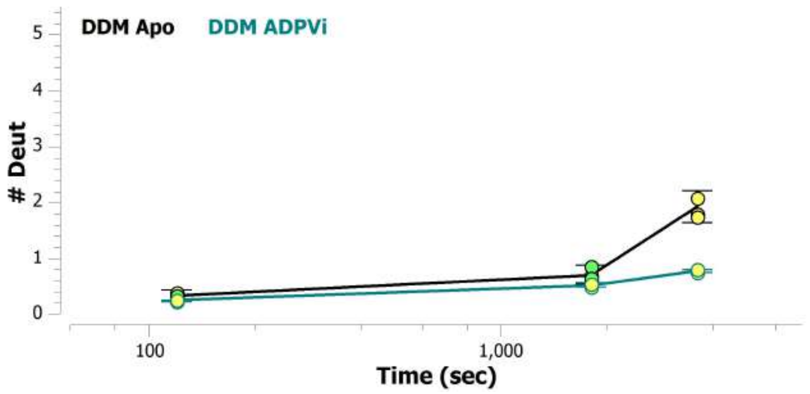


MSBA 521-533: AALDELQKNRTSL (#138)

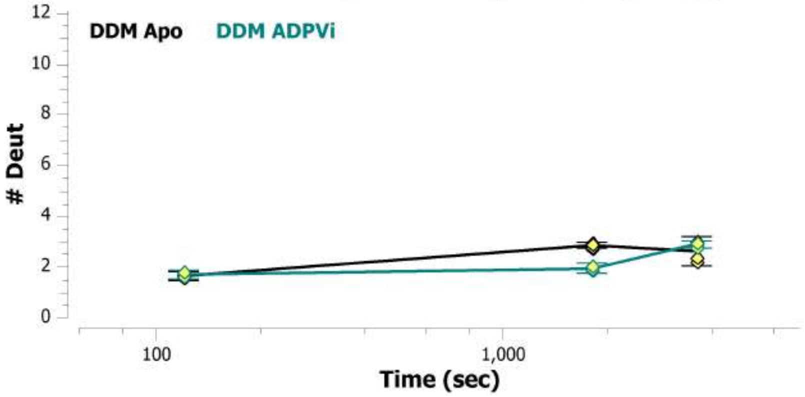


MSBA 523-533: LDELQKNRTSL (#139)

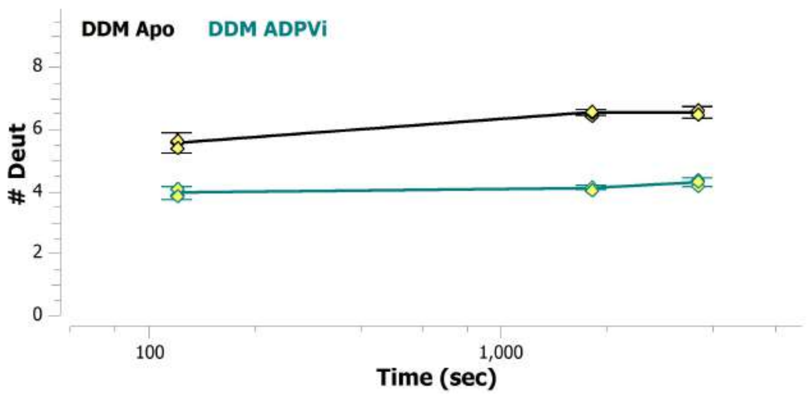

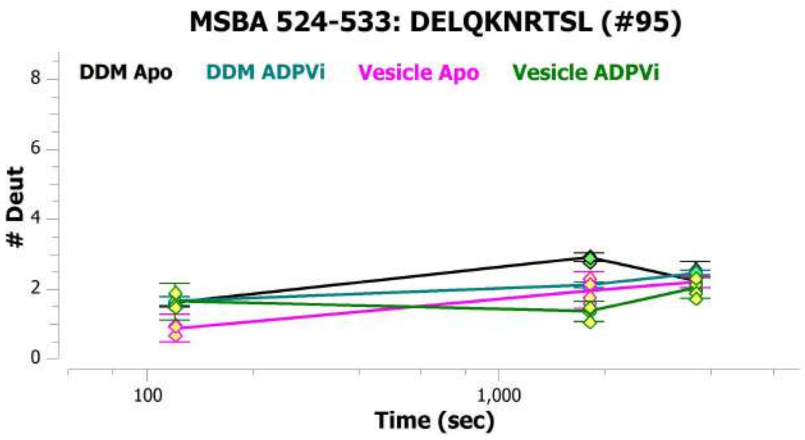
MSBA 524-533: DELQKNRTSL (#95)
DDM Apo
DDM ADPVi
Vesicle Apo
Vesicle ADPVi
### Deut
Time (sec)

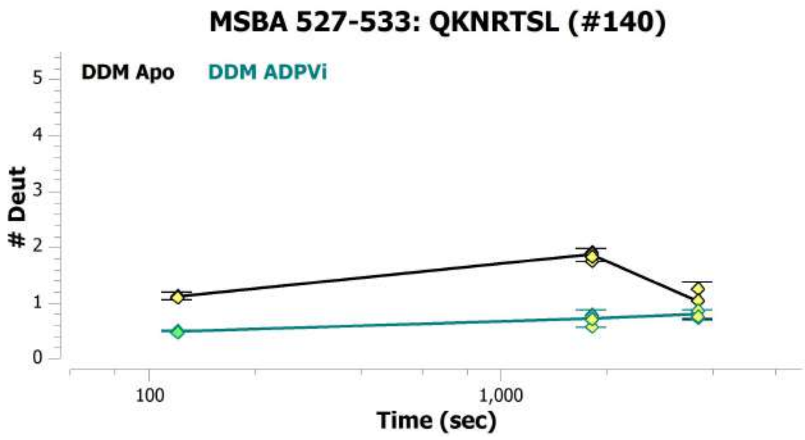
MSBA 527-533: QKNRTSL (#140)
DDM Apo
DDM ADPVi
### Deut
Time (sec)

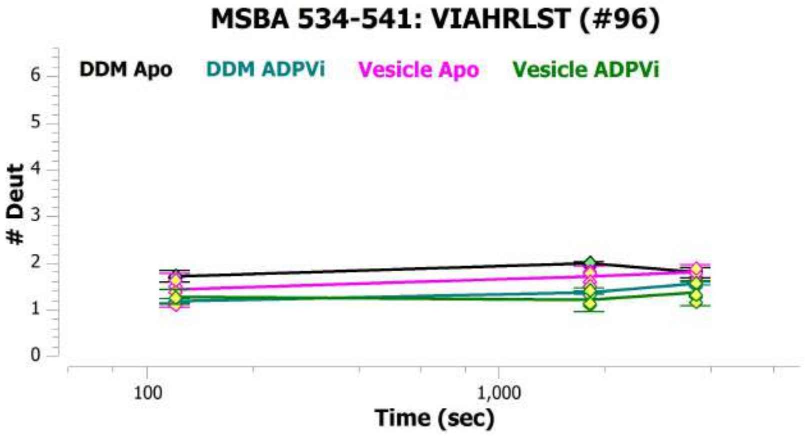
MSBA 534-541: VIAHRLST (#96)
DDM Apo
DDM ADPVi
Vesicle Apo
Vesicle ADPVi
### Deut
Time (sec)

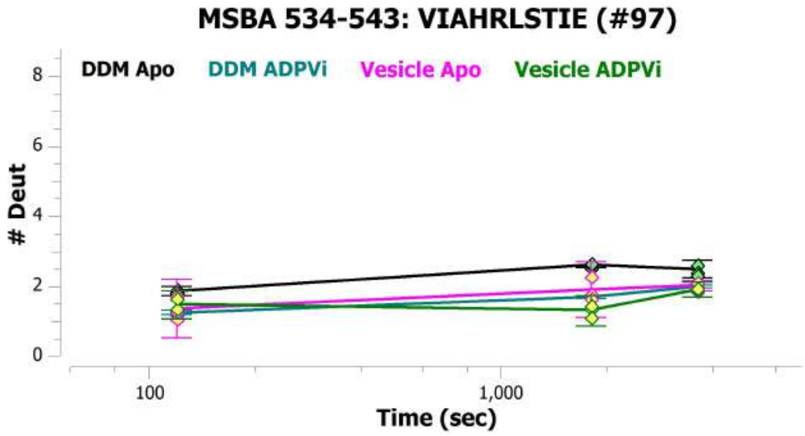
MSBA 534-543: VIAHRLSTIE (#97)
DDM Apo
DDM ADPVi
Vesicle Apo
Vesicle ADPVi
### Deut
Time (sec)

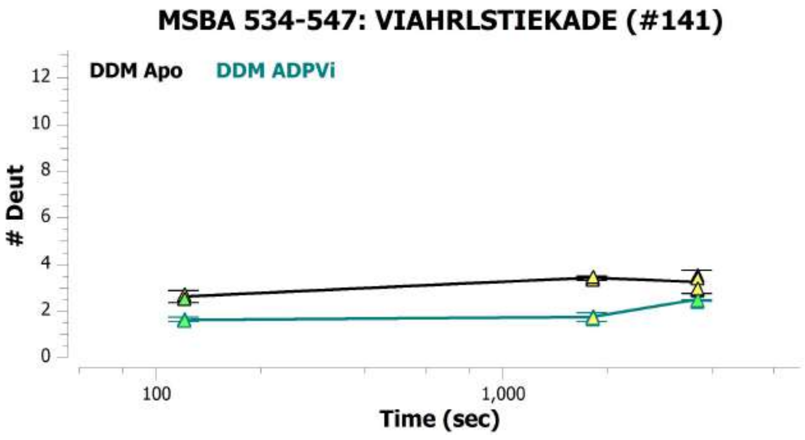
MSBA 534-547: VIAHRLSTIEKADE (#141)
DDM Apo
DDM ADPVi
### Deut
Time (sec)

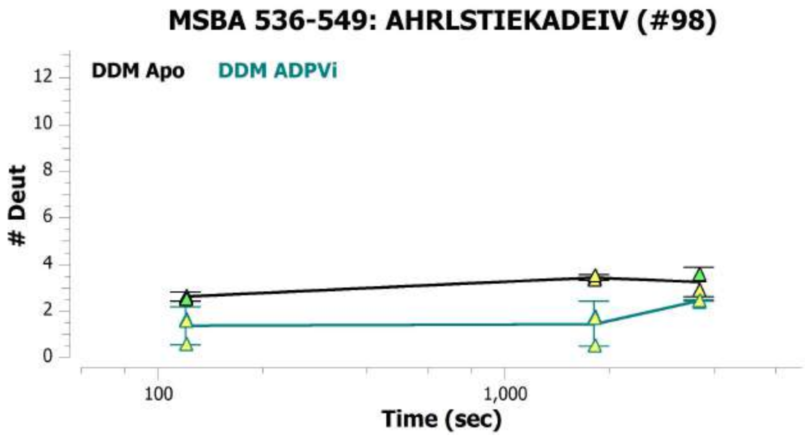
MSBA 536-549: AHRLSTIEKADEIV (#98)
DDM Apo
DDM ADPVi
### Deut
Time (sec)

MSBA 536-554: AHRLSTIEKADEIVVVEDG (#142)

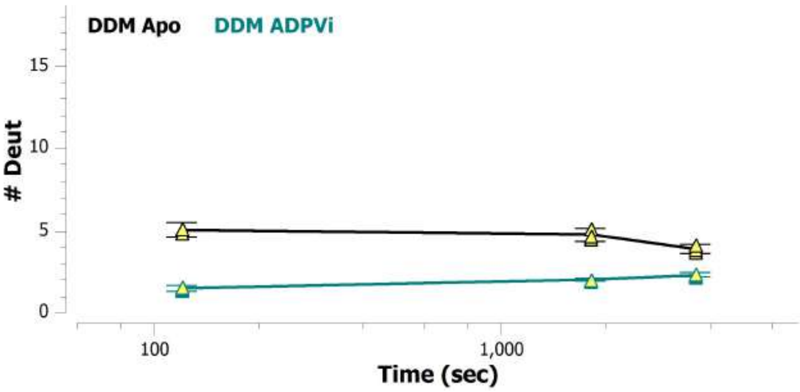


MSBA 542-547: IEKADE (#99)

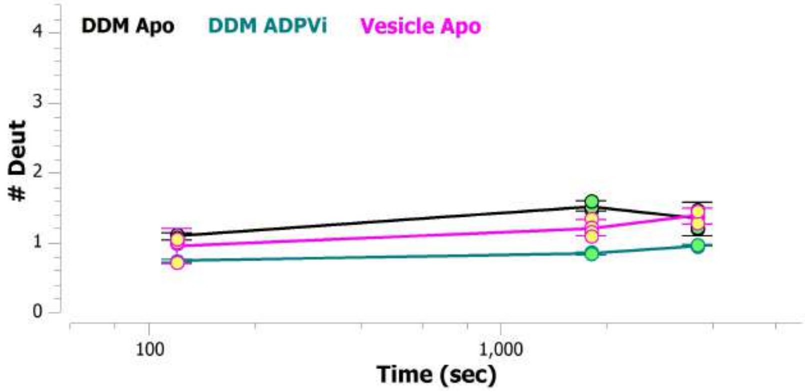


MSBA 548-553: IVVVED (#143)

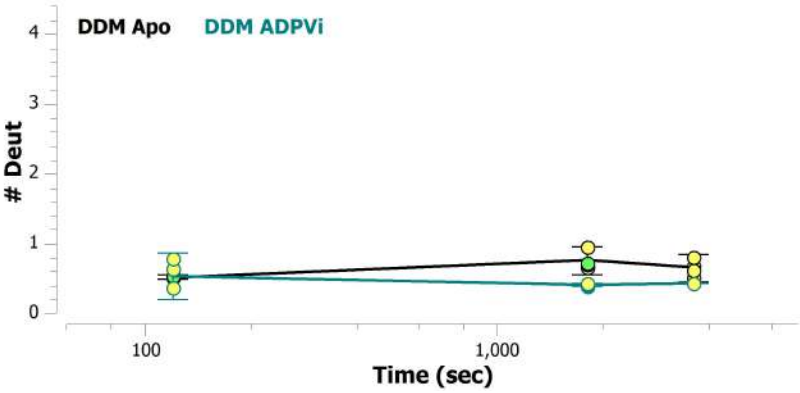


MSBA 548-554: IVVVEDG (#144)

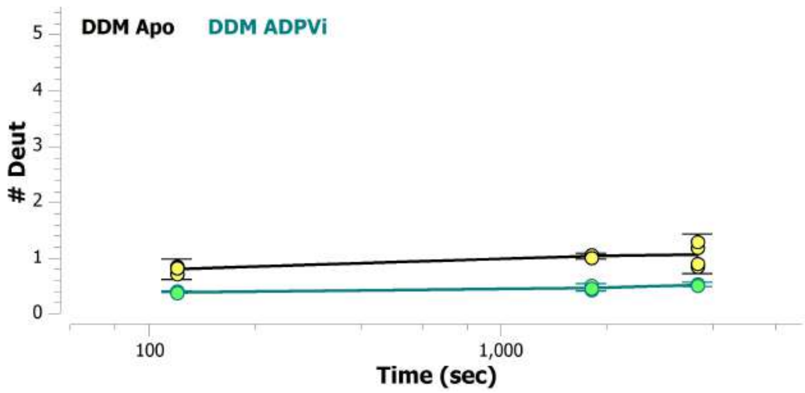


MSBA 548-555: IVVVEDGV (#100)

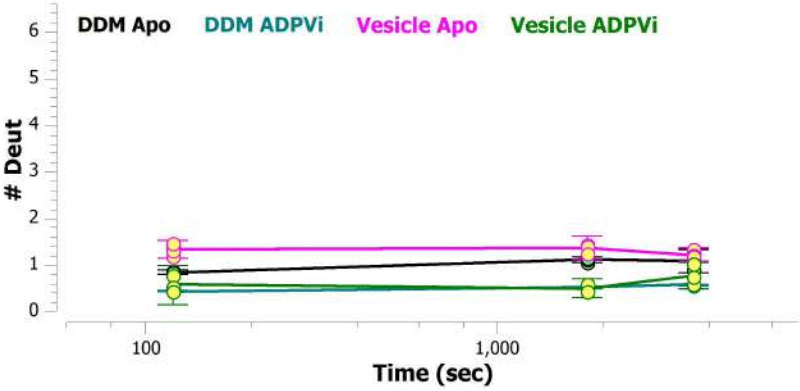


MSBA 548-556: IVVVEDGVI (#145)

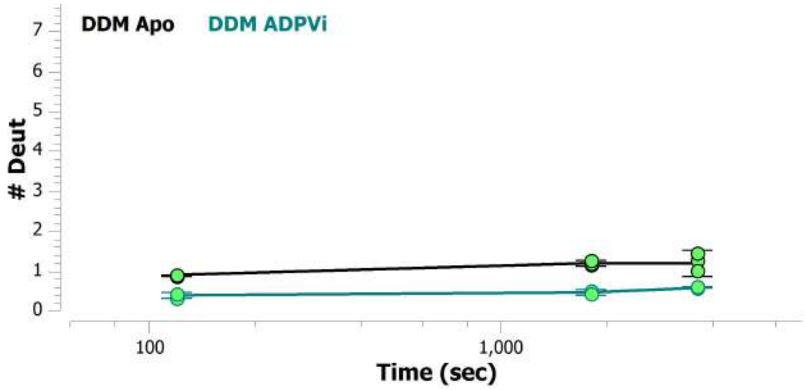

MSBA 548-558: IVVVEDGVIVE (#146)

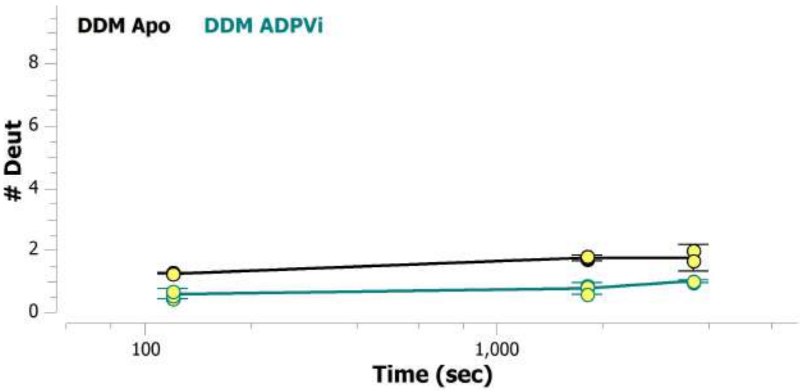


MSBA 548-565: IVVVEDGVIVERGTHNDL (#147)

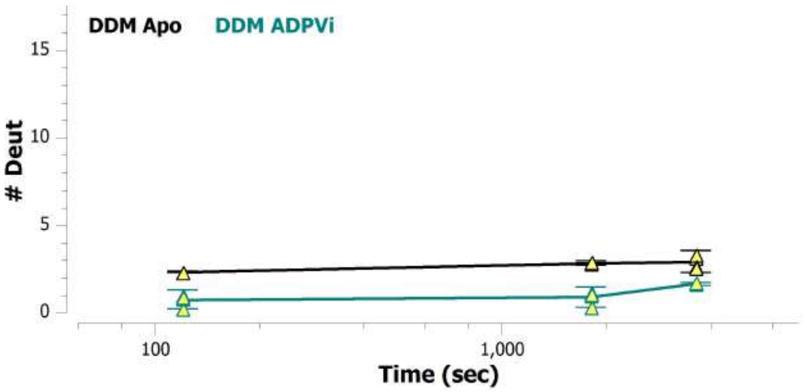


MSBA 548-572: IVVVEDGVIVERGTHNDLLEHRGVY (#148)

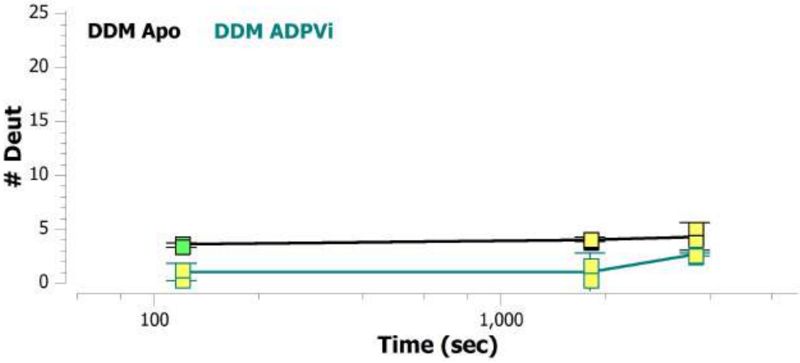


MSBA 550-556: VVEDGVI (#149)

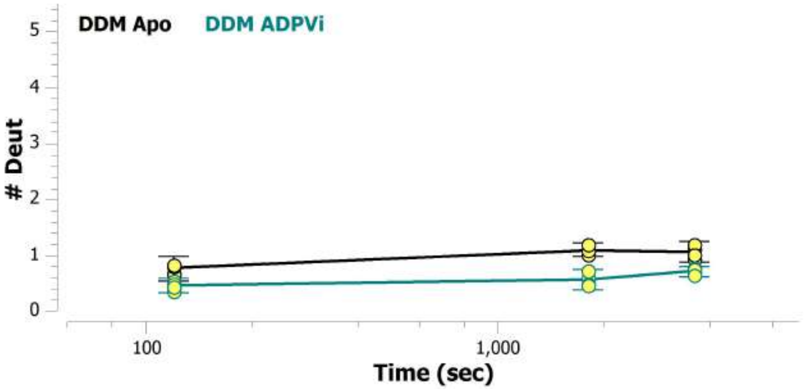


MSBA 551-556: VEDGVI (#150)

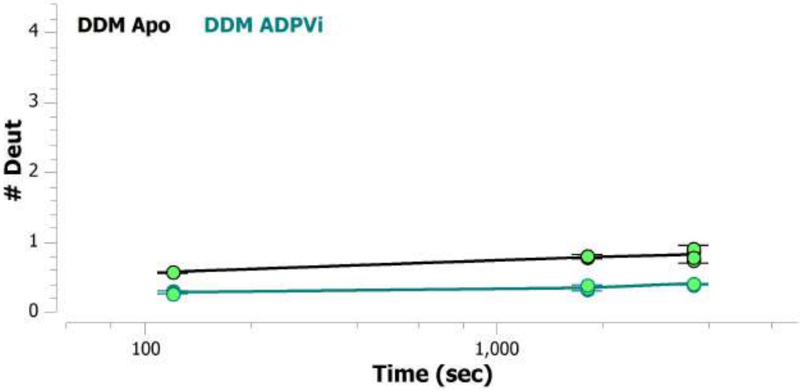


MSBA 551-558: VEDGVIVE (#151)

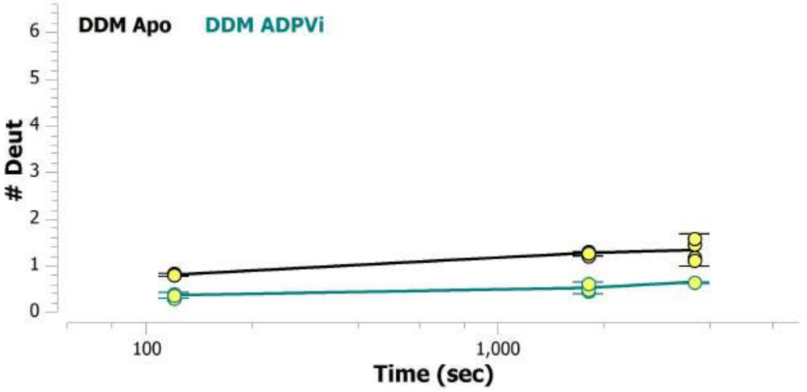

MSBA 553-558: DGVIVE (#101)

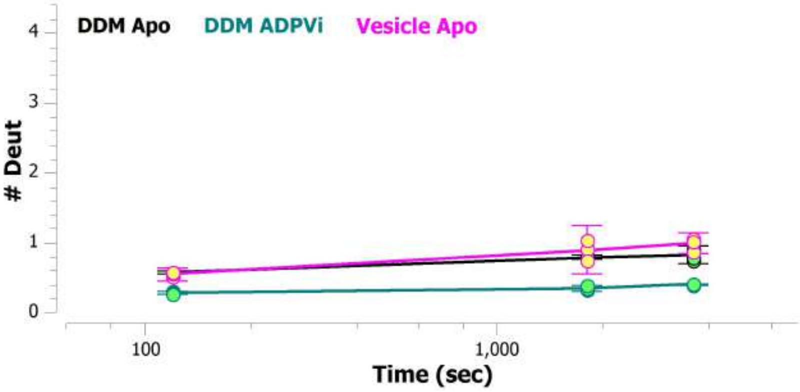


MSBA 553-565: DGVIVERGTHNDL (#152)

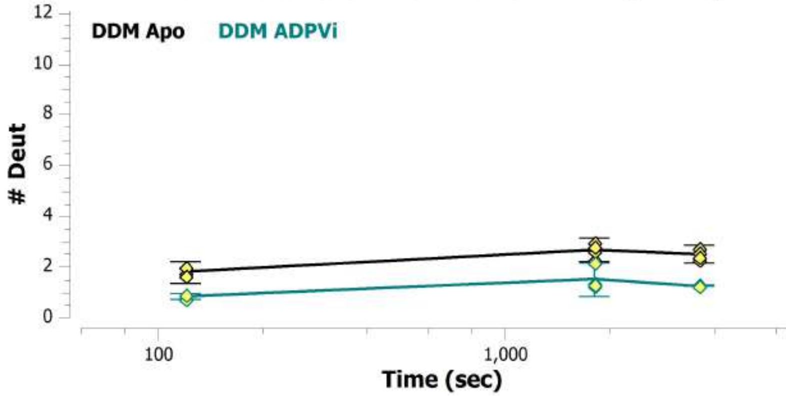


MSBA 553-572: DGVIVERGTHNDLLEHRGVY (#153)

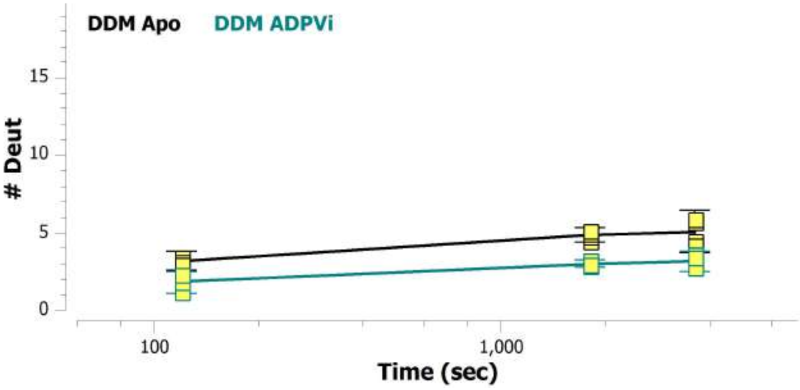


MSBA 554-565: GVIVERGTHNDL (#154)

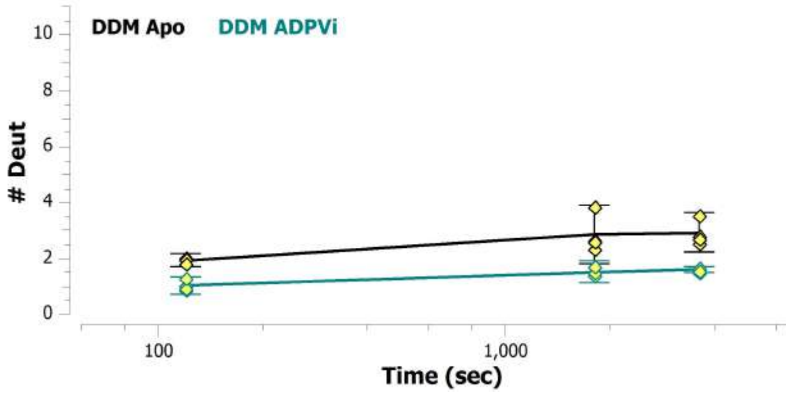


MSBA 554-566: GVIVERGTHNDLL (#155)

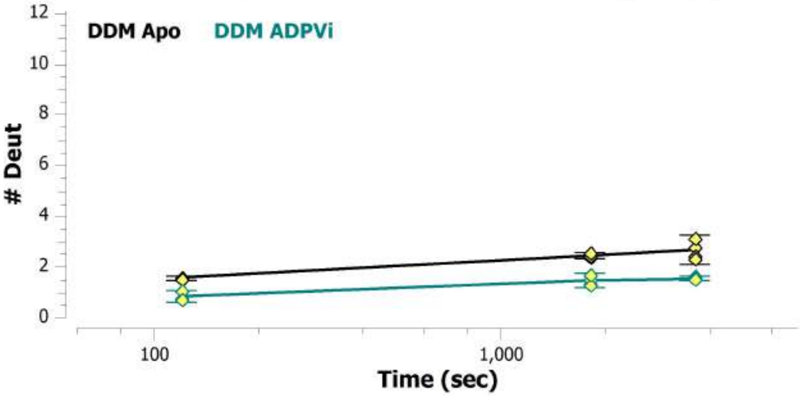


MSBA 555-565: VIVERGTHNDL (#156)

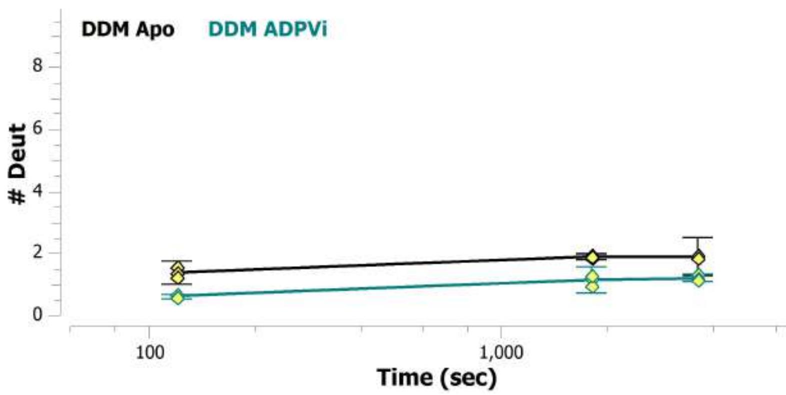

**MSBA 556-565: IVERGTHNDL (#157)**

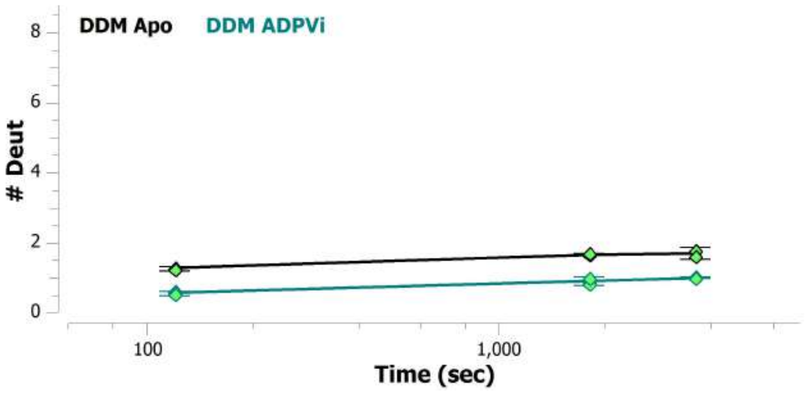


**MSBA 557-566: VERGTHNDLL (#102)**

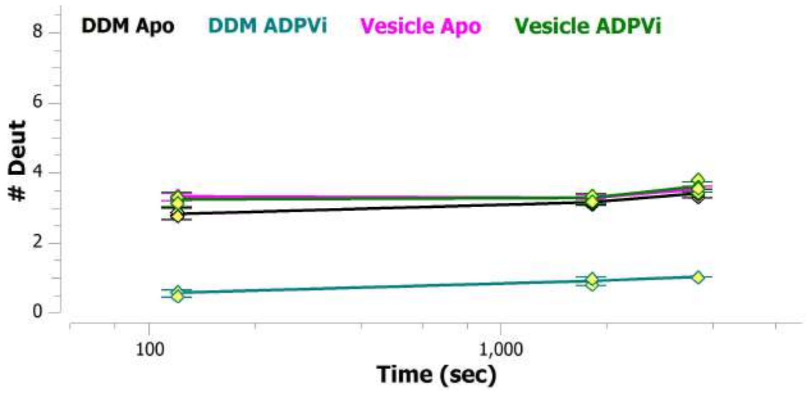


**MSBA 557-567: VERGTHNDLLE (#158)**

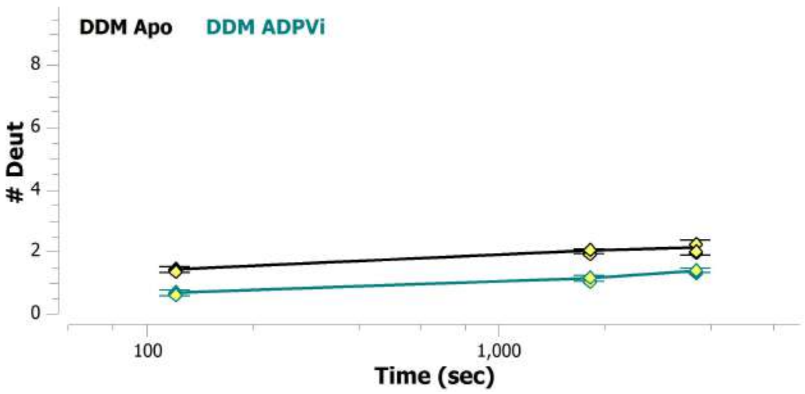


**MSBA 557-572: VERGTHNDLLEHRGVY (#103)**

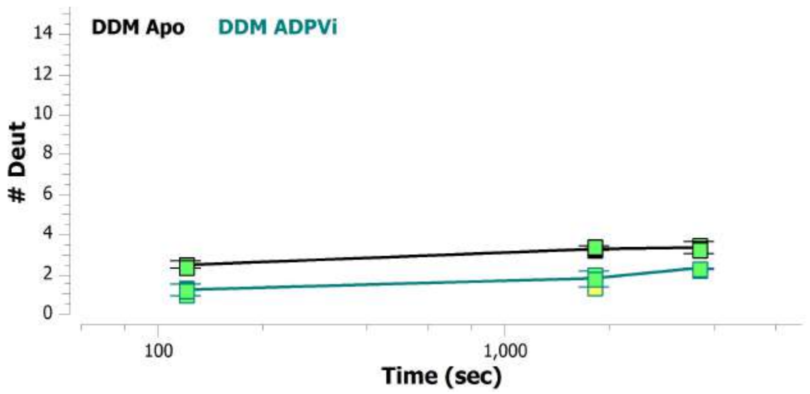


**MSBA 558-565: ERGTHNDL (#159)**

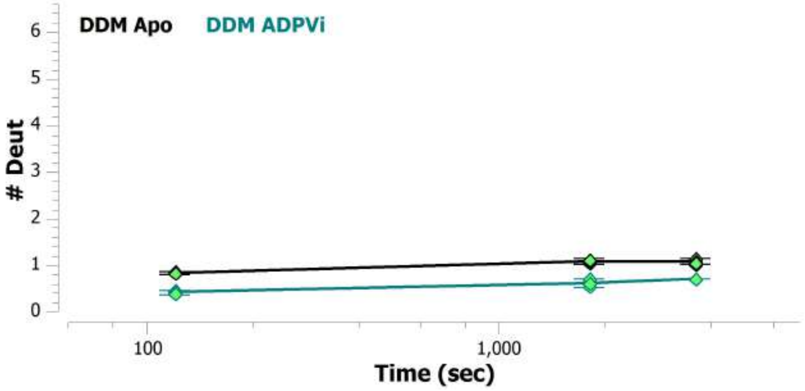


**MSBA 559-565: RGTHNDL (#160)**

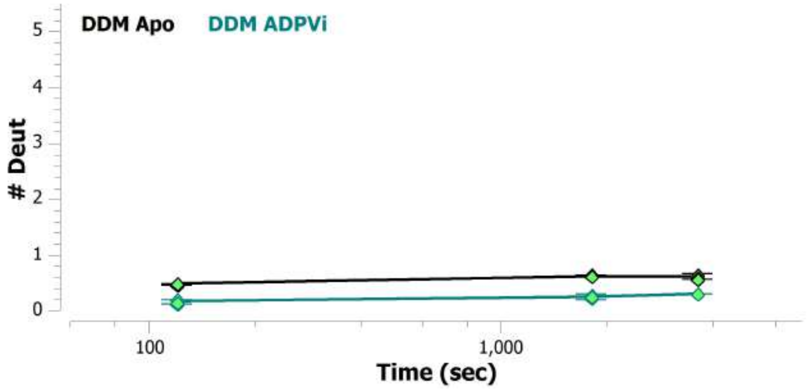

#### MSBA 559-566: RGTHNDLL (#161)

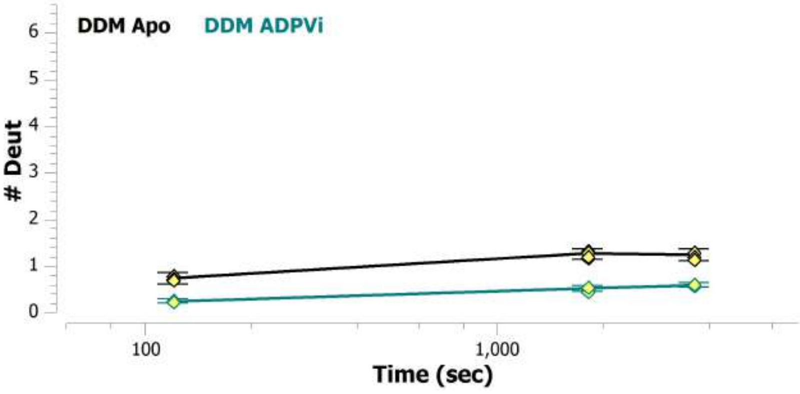


#### MSBA 559-572: RGTHNDLLEHRGVY (#162)

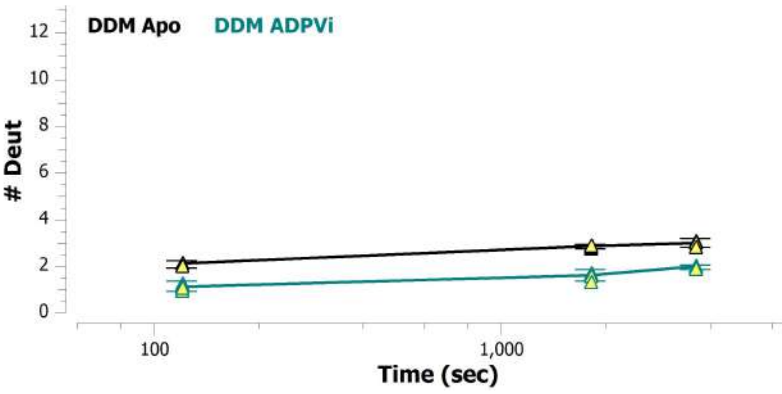


#### MSBA 566-572: LEHRGVY (#104)

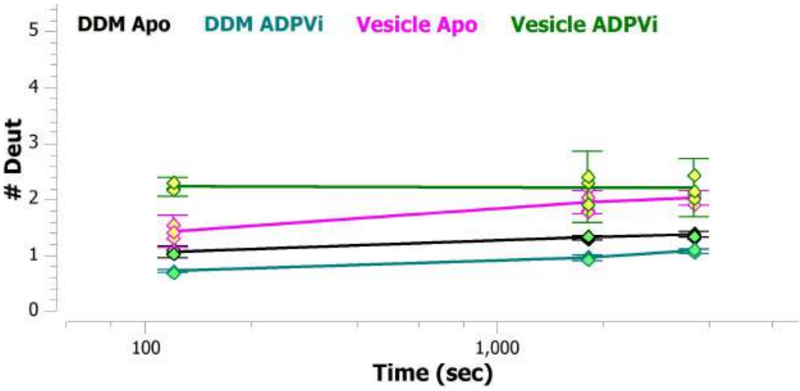


#### MSBA 567-572: EHRGVY (#163)

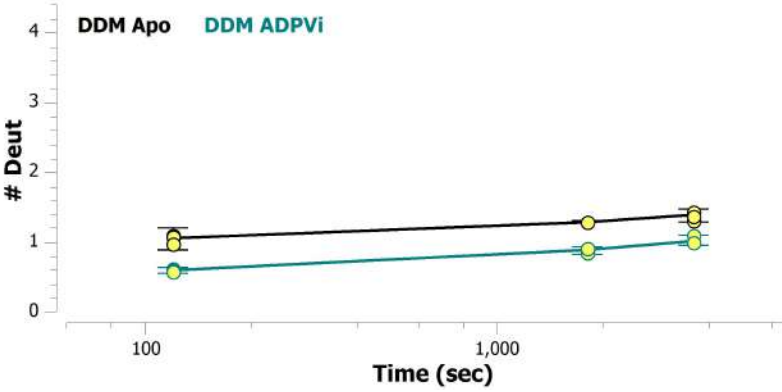


#### MSBA 65-71: VWMPLVV (#12)

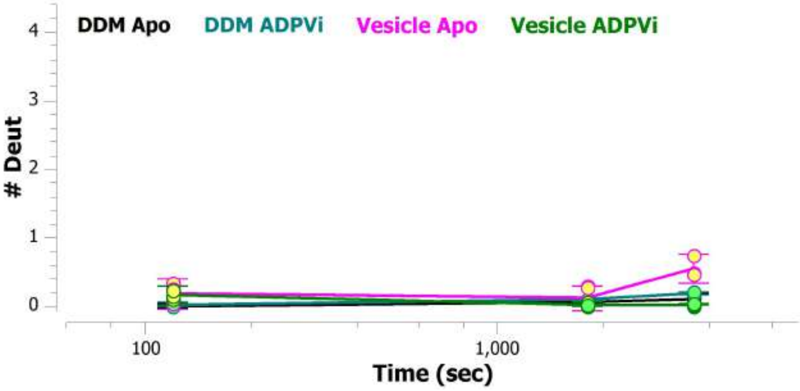


#### MSBA 65-73: VWMPLVVIG (#8)

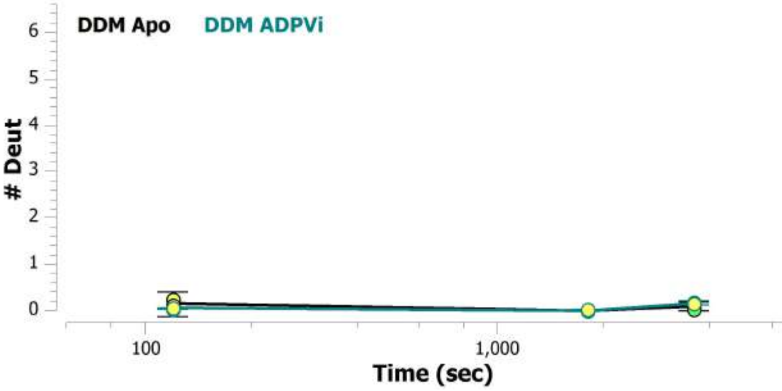

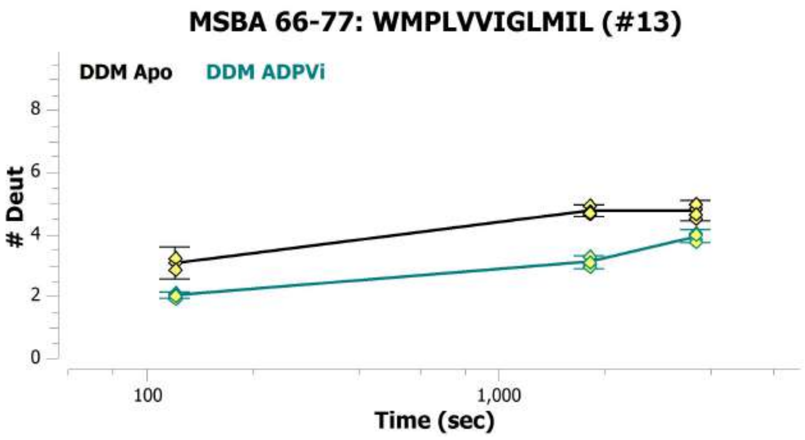
MSBA 66-77: WMPLVVIGLMIL (#13)
DDM Apo
DDM ADPVi
### Deut
0
2
4
6
8
100
1,000
Time (sec)

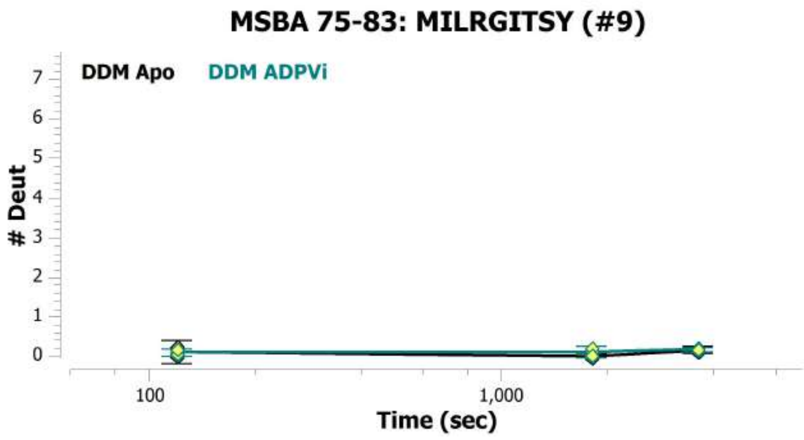
MSBA 75-83: MILRGITSY (#9)
DDM Apo
DDM ADPVi
### Deut
0
1
2
3
4
5
6
7
100
1,000
Time (sec)

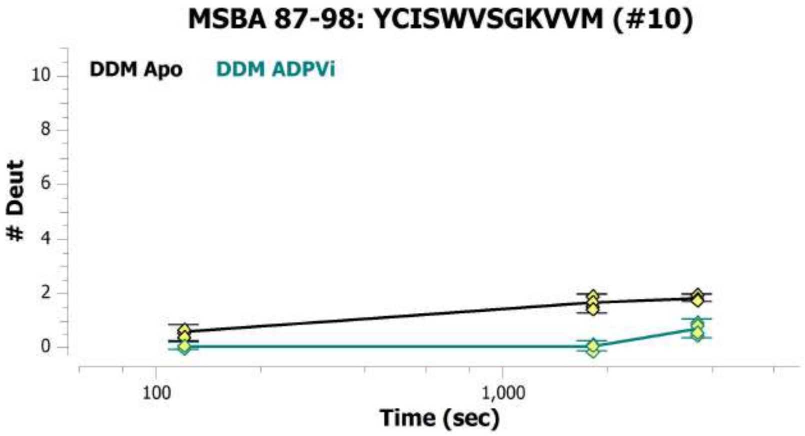
MSBA 87-98: YCISWVSGKVVM (#10)
DDM Apo
DDM ADPVi
### Deut
0
2
4
6
8
10
100
1,000
Time (sec)

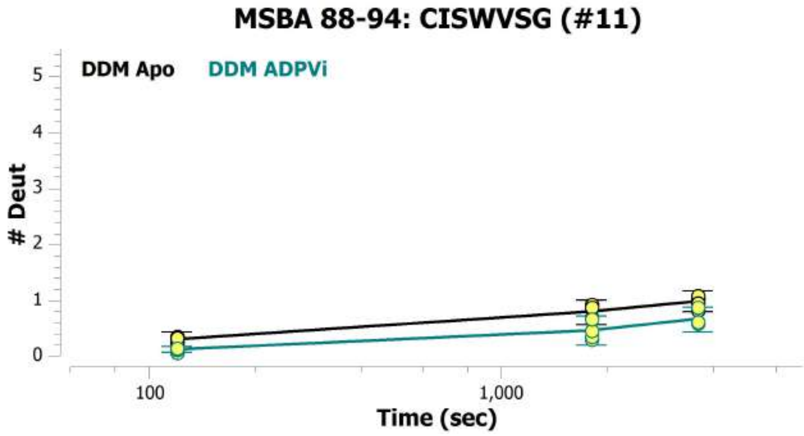
MSBA 88-94: CISWVSG (#11)
DDM Apo
DDM ADPVi
### Deut
0
1
2
3
4
5
100
1,000
Time (sec)

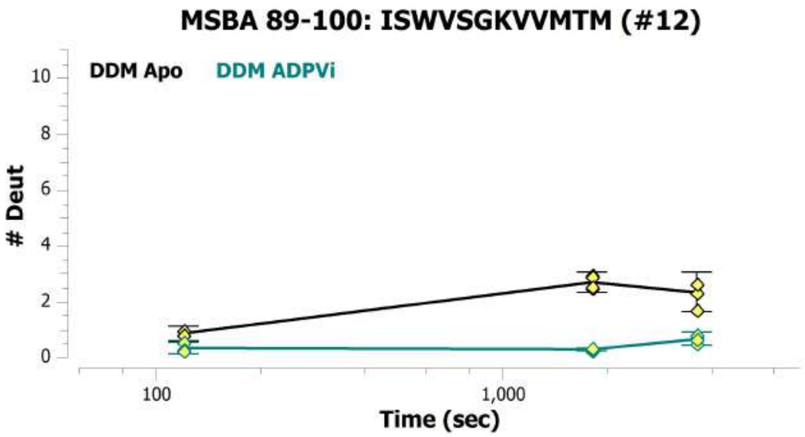
MSBA 89-100: ISWVSGKVVMTM (#12)
DDM Apo
DDM ADPVi
### Deut
0
2
4
6
8
10
100
1,000
Time (sec)

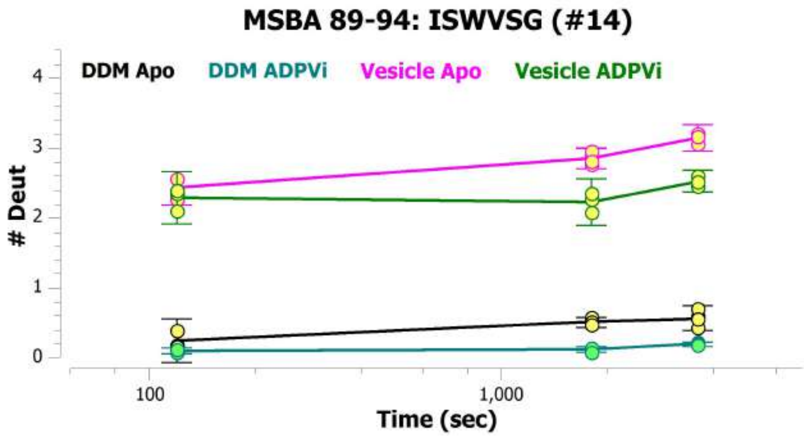
MSBA 89-94: ISWVSG (#14)
DDM Apo
DDM ADPVi
Vesicle Apo
Vesicle ADPVi
### Deut
0
1
2
3
4
100
1,000
Time (sec)

**MSBA 89-98: ISWVSGKVVM (#15)**

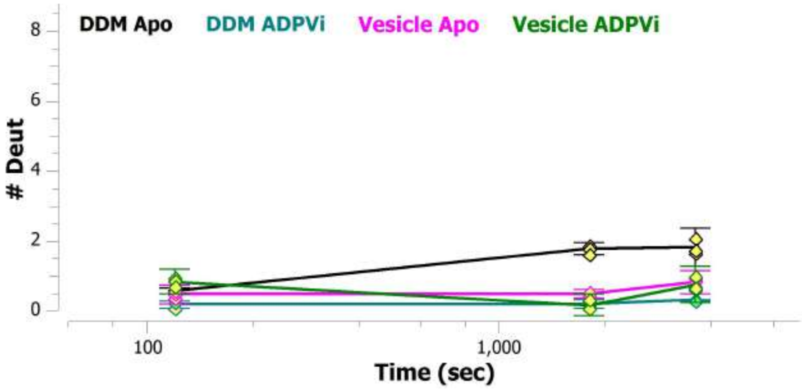


**MSBA 92-100: VSGKVVMTM (#15)**

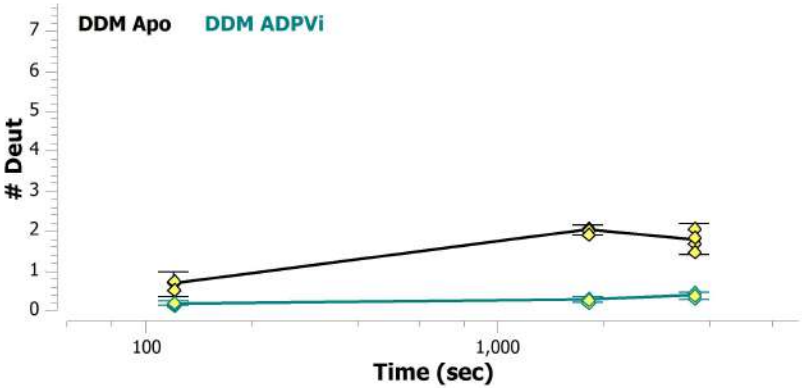


**MSBA 92-98: VSGKVVM (#13)**

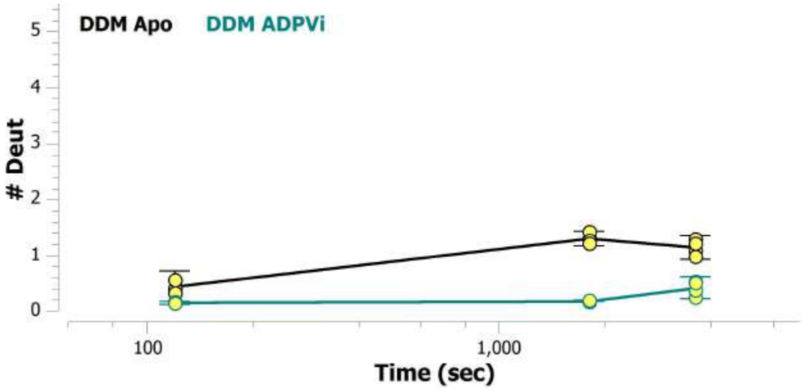


**MSBA 92-99: VSGKVVMT (#14)**

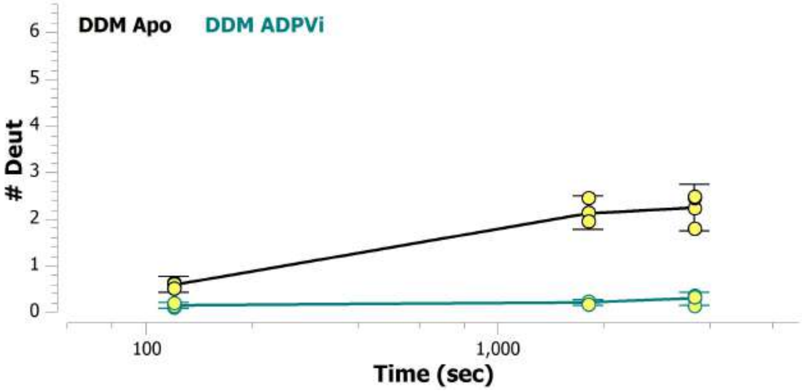


**MSBA 93-98: SGKVVM (#16)**

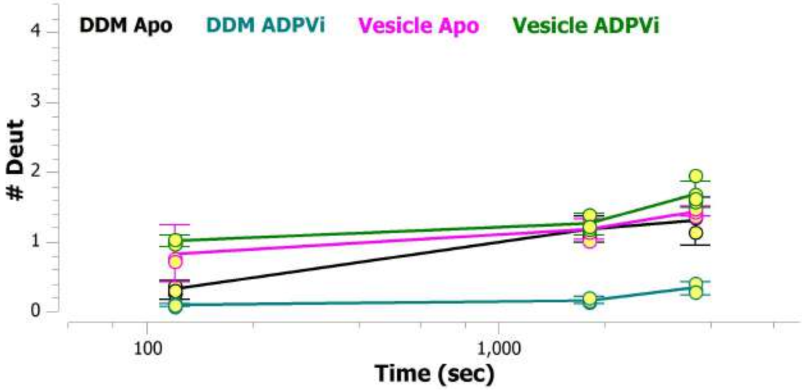


**MSBA 95-100: KVVMTM (#16)**

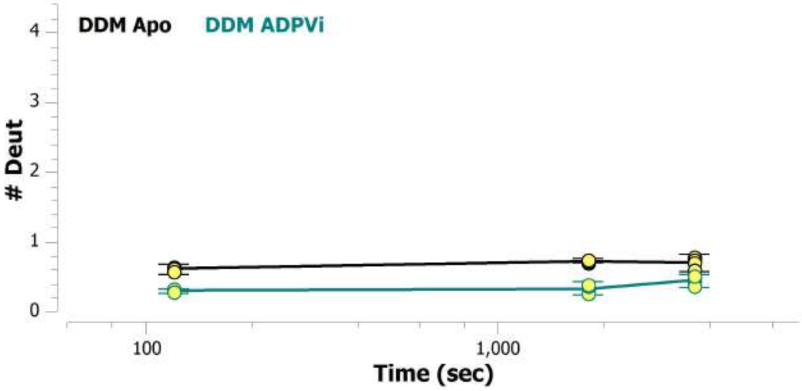

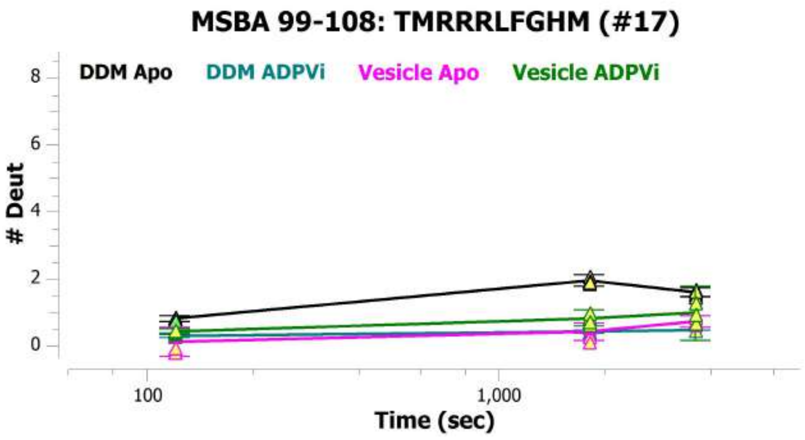
MSBA 99-108: TMRRRLFGHM (#17)
DDM Apo
DDM ADPVi
Vesicle Apo
Vesicle ADPVi
### Deut
Time (sec)
0
2
4
6
8
100
1,000

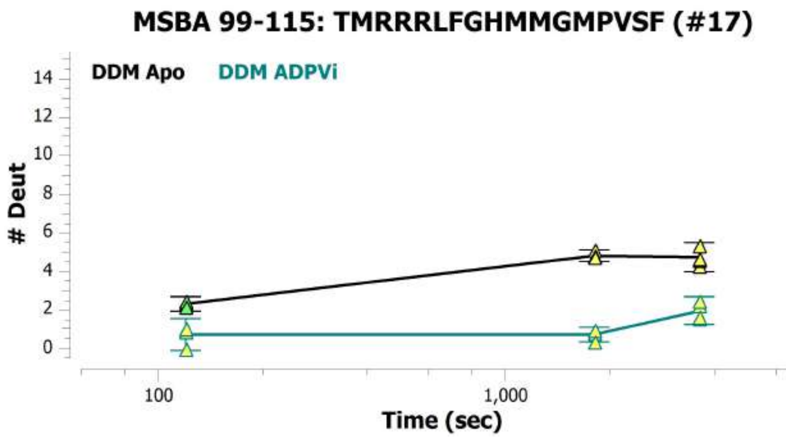
MSBA 99-115: TMRRRLFGHMMGMPVSF (#17)
DDM Apo
DDM ADPVi
### Deut
Time (sec)
0
2
4
6
8
10
12
14
100
1,000
